## Supplementary Figures for "Tradeoff Between Speed and Robustness in Primordium Initiation Mediated by Auxin-CUC1 Interaction"

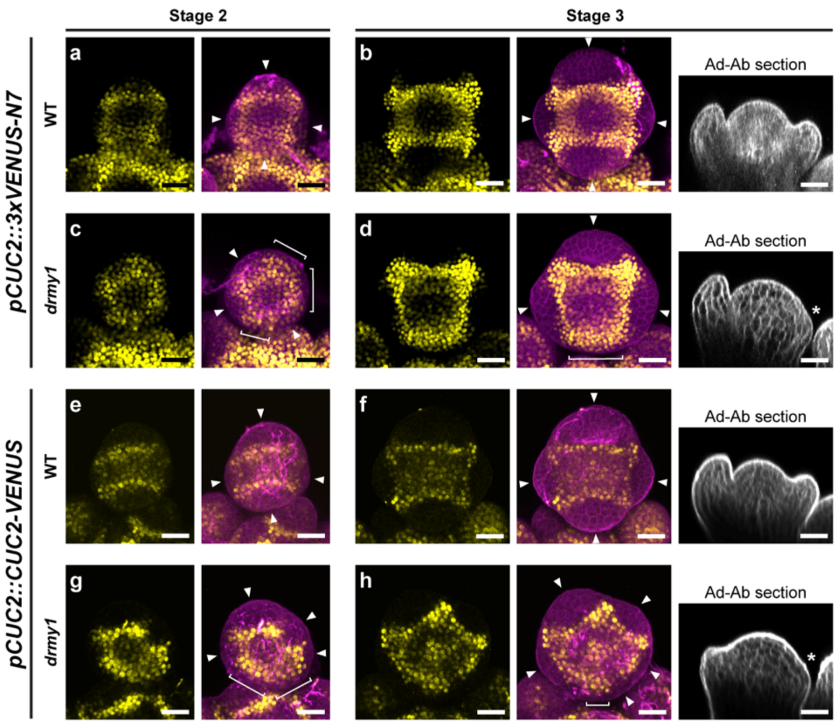


**Supplementary Fig. 1. CUC2 patterning is disrupted in *drmy1*.**

**(a-d)** *CUC2* expression pattern in WT **(a,b)** and *drmy1* **(c,d)**, before (stage 2) or after (stage 3) sepal initiation. Shown are images of *pCUC2::3xVENUS-N7* (yellow) by itself and merged with PI (magenta). For stage 3, optical sections along the abaxial-adaxial axis are also shown. Images are representative of n = 12 WT stage 2 buds, n = 6 WT stage 3 buds, n = 11 *drmy1* stage 2 buds, and n = 13 *drmy1* stage 3 buds. Note that plants are of Ler background and are heterozygous for the *pCUC2::3xVENUS-N7* transgene.

**(e-h)** *CUC2* protein accumulation pattern in WT **(e,f)** and *drmy1* **(g,h)**, before (stage 2) or after (stage 3) sepal initiation. Shown are images of *pCUC2::CUC2-VENUS* (yellow) by itself and merged with PI (magenta). For stage 3, optical sections along the abaxial-adaxial axis are shown. Images are representative of n = 15 WT stage 2 buds, n = 6 WT stage 3 buds, n = 18 *drmy1* stage 2 buds, and n = 14 *drmy1* stage 3 buds. Note that plants are of Ler background.

Note that in WT, *CUC2* is expressed in boundaries between the center of the floral meristem and four robustly positioned (incipient) sepal primordia (arrowheads). In *drmy1*, *CUC2* is expressed in variably positioned boundaries (arrowheads), and in the bud periphery corresponding to regions of no sepal outgrowth (brackets; asterisks). Scale bars, 25 μm.

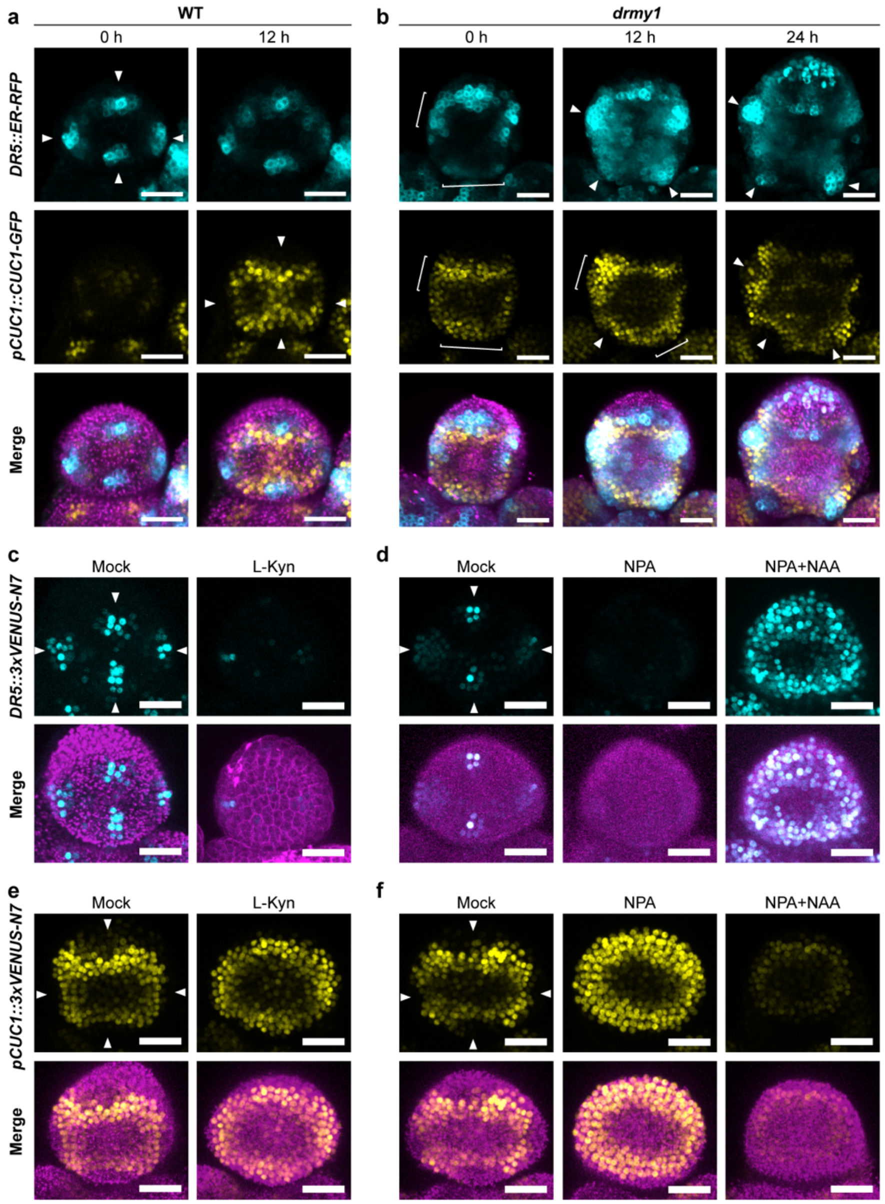


**Supplementary Fig. 2. Strong, robustly positioned auxin maxima are required for robust boundary expression of *CUC1***

**(a-b)** Live imaging of a WT **(a)** and *drmy1* **(b)** stage 2 bud carrying *DR5::ER-RFP* (cyan, top row) and *pCUC1::CUC1-GFP* (yellow, middle row). Chlorophyll (magenta) is shown together with both markers (bottom row). In WT, four robustly positioned auxin maxima appear in the four incipient sepal primordia, before boundary accumulation of CUC1 (arrowheads). In *drmy1*, weak and diffuse auxin signal frequently colocalizes with CUC1 which accumulates in the bud periphery (brackets). As auxin signal concentrates into variably positioned auxin maxima, CUC1 retreats from the bud periphery and accumulates in variably positioned boundaries (arrowheads). Images are representative of n = 3 buds per genotype.

**(c-d)** *DR5::3xVENUS-N7* in WT background treated with mock or 80 μM L-Kyn for 4 days **(c)**, or transiently treated with mock, 100 μM NPA, or 100 μM NPA + 20 μM NAA for 24 hours and observed 2 days later **(d)**. DR5 is shown on the top, and DR5 merged with chlorophyll is shown on the bottom. Note that strong, robustly positioned auxin maxima (arrowheads) are disrupted by L-Kyn, NPA, or NPA+NAA treatments. Images are representative of n=5 (mock in **(c)**), n=7 (L-Kyn), n=6 (mock in **(d)**), n=8 (NPA), and n=7 (NPA+NAA) buds.

**(e-f)** *pCUC1::3xVENUS-N7* in WT background treated with mock or 80 μM L-Kyn for 4 days **(e)**, or transiently treated with mock, 100 μM NPA, or 100 μM NPA + 20 μM NAA for 24 hours and observed 3 days later **(f)**. *pCUC1::3xVENUS-N7* is shown on the top, and *pCUC1::3xVENUS-N7* together with chlorophyll is shown on the bottom. Note that boundary expression of *CUC1* (arrowheads) are disrupted by the treatments. In L-Kyn and NPA, *CUC1* expression expands to the entire bud periphery, whereas in NPA+NAA, *CUC1* expression is restricted to a weak ring immediately inside the bud periphery. Images are representative of n=8 (mock in **(e)**), n=9 (L-Kyn), n=5 (mock in **(f)**), n=10 (NPA), and n=6 (NPA+NAA) buds. Scale bars, 25 μm.

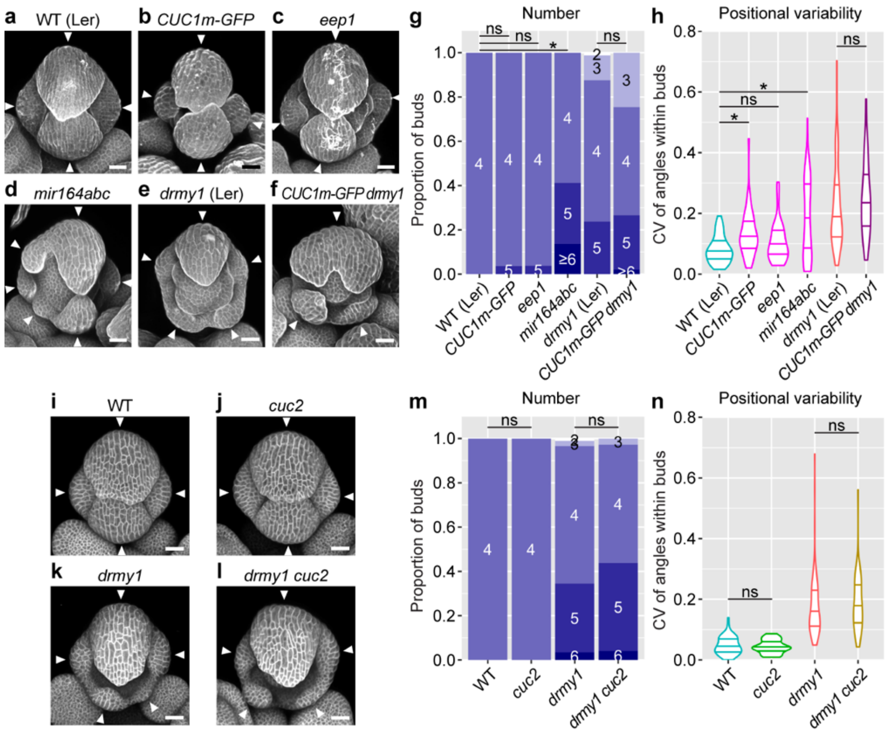


**Supplementary Fig. 3. Differential contribution of *CUC1* and *CUC2* to variability in sepal initiation.**

**(a-f)** Various lines upregulating *CUC1* show enhanced variability in sepal initiation, while further upregulation of *CUC1* in *drmy1* does not further enhance the variability. Shown are PI-stained buds of WT **(a)**, *CUC1m-GFP* **(b)**, *eep1* **(c)**, *mir164abc* **(d)**, *drmy1* **(e)**, and *CUC1m-GFP drmy1* **(f)**. Arrowheads point to sepal primordia. All lines were in Ler WT background except *mir164abc*, which was in a mixed Ler-Col background. Scale bars, 25 µm.

**(g-h)** Quantification of sepal primordium number **(g)** and positional variability **(h)**. Sample size: WT, n = 58 buds; *mCUC1-GFP*, n = 55 buds; *eep1*, n = 54 buds; *mir164abc*, n = 51 buds; *drmy1*, n = 80 buds; *mCUC1-GFP drmy1*, n = 49. For **(g)**, p-values from Fisher's contingency table tests: WT vs. *CUC1m-GFP*, p = 0.2347; WT vs *eep1*, p = 0.2302; WT vs. *mir164abc*, p = $7.512\times10^{-9}$; *drmy1* vs *CUC1m-GFP drmy1*, p = 0.1062. For **(h)**, p-values from Wilcoxon's rank sum tests: WT vs. *CUC1m-GFP*, p = $5.311\times10^{-5}$; WT vs. *eep1*, p = 0.0628; WT vs. *mir164abc*, p = $1.051\times10^{-3}$; *drmy1* vs *CUC1m-GFP*, p = 0.1035.

**(i-l)** Mutation of *CUC2* does not reduce variability in *drmy1*. Shown are WT **(i)**, *cuc2* **(j)**, *drmy1* **(k)**, and *drmy1 cuc2* **(l)**, all carrying a membrane marker *UBQ10::mCitrine-RCI2A*. Arrowheads point to sepal primordia. Scale bars, 25 µm.

**(m-n)** Quantification of sepal primordium number **(m)** and positional variability **(n)**. Data of WT and *drmy1* were reused from Fig. 2j-k. Sample size: WT, n = 66 buds; *cuc2*, n = 58 buds; *drmy1*, n = 87 buds; *drmy1 cuc2*, n = 73 buds. For **(m)**, p-values from Fisher's contingency table tests: WT vs. *cuc2*, p = 1; *drmy1* vs. *drmy1 cuc2*, p = 0.7234. For **(n)**, p-values from Wilcoxon's rank sum tests: WT vs. *cuc2*, p = 0.7790; *drmy1* vs. *drmy1 cuc2*, p = 0.2285.

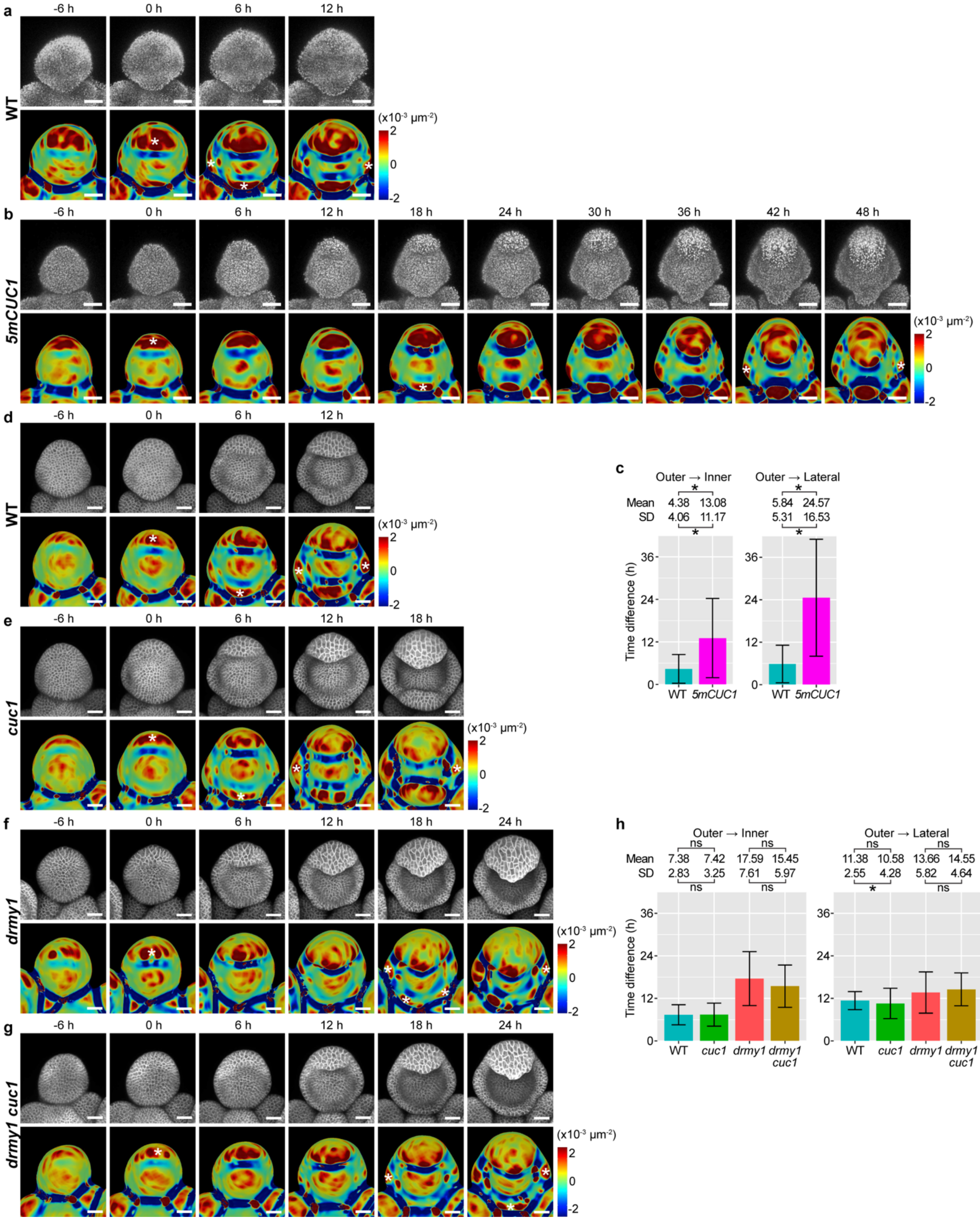
a
-6 h
0 h
6 h
12 h
WT
(x10⁻³ μm⁻²)
2
0
-2
b
-6 h
0 h
6 h
12 h
18 h
24 h
30 h
36 h
42 h
48 h
5mCUC1
(x10⁻³ μm⁻²)
2
0
-2
c
Outer → Inner
Mean 4.38 13.08
SD 4.06 11.17
Outer → Lateral
5.84 24.57
5.31 16.53
Time difference (h)
36
24
12
0
WT
5mCUC1
d
-6 h
0 h
6 h
12 h
WT
(x10⁻³ μm⁻²)
2
0
-2
e
-6 h
0 h
6 h
12 h
18 h
cuc1
(x10⁻³ μm⁻²)
2
0
-2
f
-6 h
0 h
6 h
12 h
18 h
24 h
drmy1
(x10⁻³ μm⁻²)
2
0
-2
g
-6 h
0 h
6 h
12 h
18 h
24 h
drmy1 cuc1
(x10⁻³ μm⁻²)
2
0
-2
h
Outer → Inner
ns
ns
Mean 7.38 7.42 17.59 15.45
SD 2.83 3.25 7.61 5.97
ns
ns
Outer → Lateral
ns
ns
11.38 10.58 13.66 14.55
2.55 4.28 5.82 4.64
ns
Time difference (h)
36
24
12
0
WT
cuc1
drmy1
drmy1 cuc1

**Supplementary Fig. 4. CUC1 overexpression is sufficient but not necessary for variable sepal initiation timing.**

**(a-c)** *CUC1* upregulation (*5mCUC1*) disrupts robustness in sepal initiation timing. Shown are representative 6 h-interval image series of sepal initiation in WT **(a)** and *5mCUC1* **(b)**. **(c)** shows quantification of time between outer and inner sepal initiation (left), and between outer and lateral sepal initiation (right). Note that *5mCUC1* causes a significant increase in time between outer sepal initiation and the initiation of all other sepals (mean). This time is also more variable among different buds of *5mCUC1* (SD). Mean and SD values are shown on top, and mean ± SD is shown in the graphs. Sample size for outer vs. inner, n = 48 buds for WT, n = 39 buds for *5mCUC1*. Sample size for outer vs. lateral, n = 38 buds for WT, n = 21 buds for *5mCUC1*. For mean, p-values from Wilcoxson's rank sum tests are: outer → inner, $3.714\times10^{-7}$; outer → lateral, $1.622\times10^{-5}$. For SD, p-values from Levene's tests are: outer → inner, $3.554\times10^{-3}$; outer → lateral, $1.209\times10^{-6}$.

**(d-h)** The *cuc1* mutation does not restore robust sepal initiation timing in *drmy1*. Shown are representative 6 h-interval image series of sepal initiation in WT **(d)**, *cuc1* **(e)**, *drmy1* **(f)**, and *drmy1 cuc1* **(g)** carrying a membrane marker *35S::mCitrine-RCI2A*. **(h)** shows quantification of time between outer and inner sepal initiation (left), and between outer and lateral sepal initiation (right). Note that while *drmy1* shows delayed inner sepal initiation relative to the outer sepal (mean), and more variability among different buds (SD), the *cuc1* mutation fails to rescue either aspect. Mean and SD values are shown on top, and mean ± SD is shown in the graphs. Sample size: n = 48 buds for WT, n = 38 buds for *cuc1*, n = 58 buds for *drmy1*, and n = 47 buds for *drmy1 cuc1*. For mean, p-values from Wilcoxson's rank sum tests are: outer → inner, WT vs. *cuc1*, 0.8877; *drmy1* vs. *drmy1 cuc1*, 0.1605; outer → lateral, WT vs. *cuc1*, 0.3617; *drmy1* vs. *drmy1 cuc1*, 0.4593. For SD, p-values from Levene's tests are: outer → inner, WT vs. *cuc1*, 0.3638, *drmy1* vs. *drmy1 cuc1*, 0.5248; outer → lateral, WT vs. *cuc1*, $1.082\times10^{-3}$; *drmy1* vs. *drmy1 cuc1*, 0.2119.

In **(a,b,d-g)**, top rows show the chlorophyll channel **(a,b)** or the mCitrine channel **(d-g)**, and bottom rows show Gaussian curvature heatmaps made from extracted surfaces. A sepal primordium is considered initiated (asterisks) when it forms a dark red band of positive Gaussian curvature (primordium) adjacent to a dark blue band of negative Gaussian curvature (boundary) (see Methods). Scale bars, 25 μm.

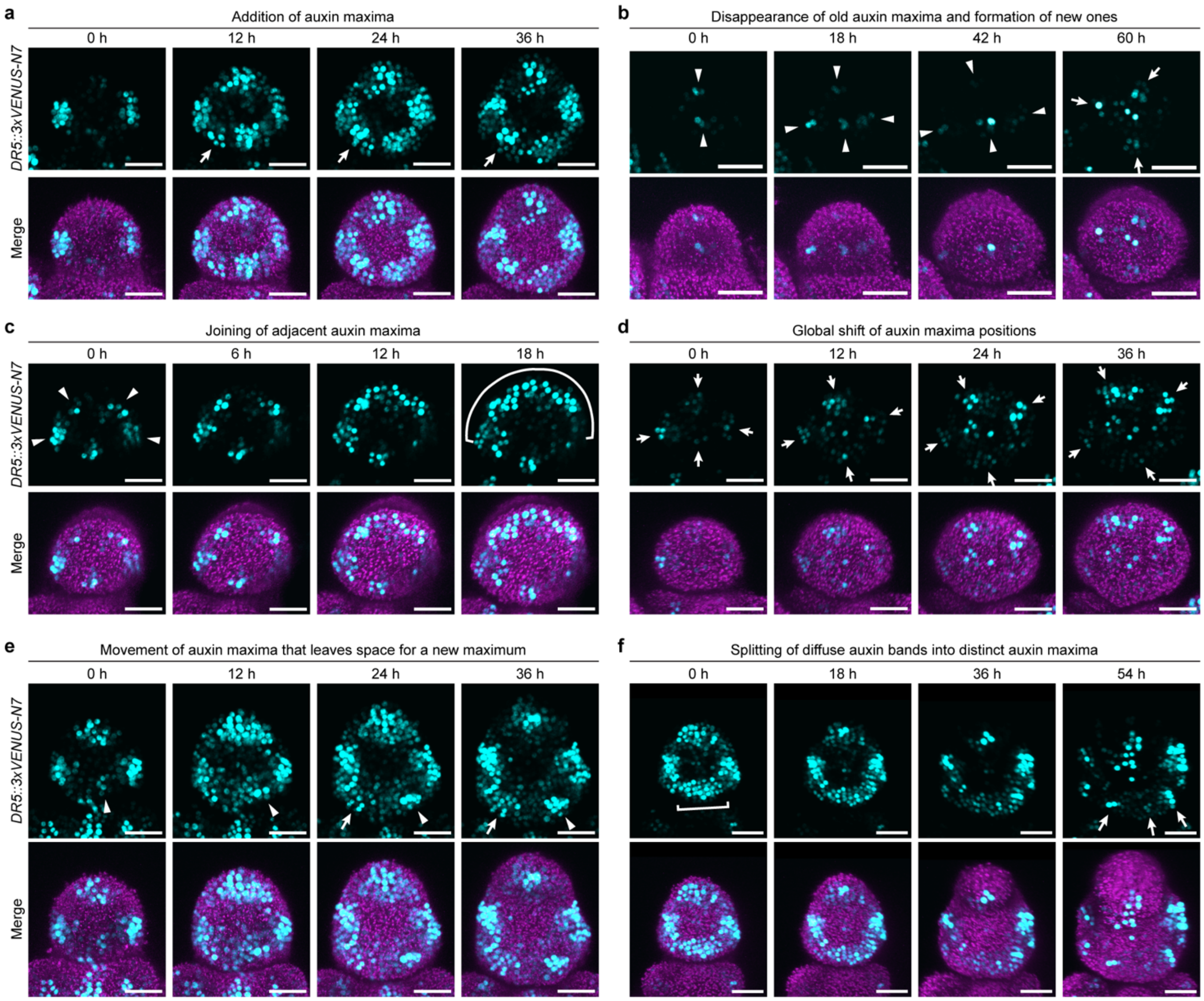


**Supplementary Fig. 5. Dynamic changes in auxin patterning in the developing buds of *5mCUC1*.**

Shown are DR5::3xVENUS-N7 (top rows), and DR5 merged with the chlorophyll channel (bottom rows), in stage 2 to stage 3 buds of *5mCUC1* live imaged through time. **(a)** Additional auxin maxima formed (arrow). **(b)** At the beginning of stage 2, four auxin maxima formed at seemingly robust positions (arrowheads). Three of them then disappeared, and three new auxin maxima formed (arrows). Signal for both DR5 and chlorophyll has been increased relative to other panels in order to reveal the pattern. **(c)** Joining of four auxin maxima (arrowheads) into one diffuse band (curved bracket). **(d)** Four auxin maxima that initially formed at robust positions shifted collectively (arrows). **(e)** The auxin maximum for the incipient inner sepal moved (arrowhead), leaving space for an additional auxin maximum (arrow). **(f)** A diffuse band of auxin signaling (bracket) split into three distinct auxin maxima (arrows). Scale bars, 25 μm.

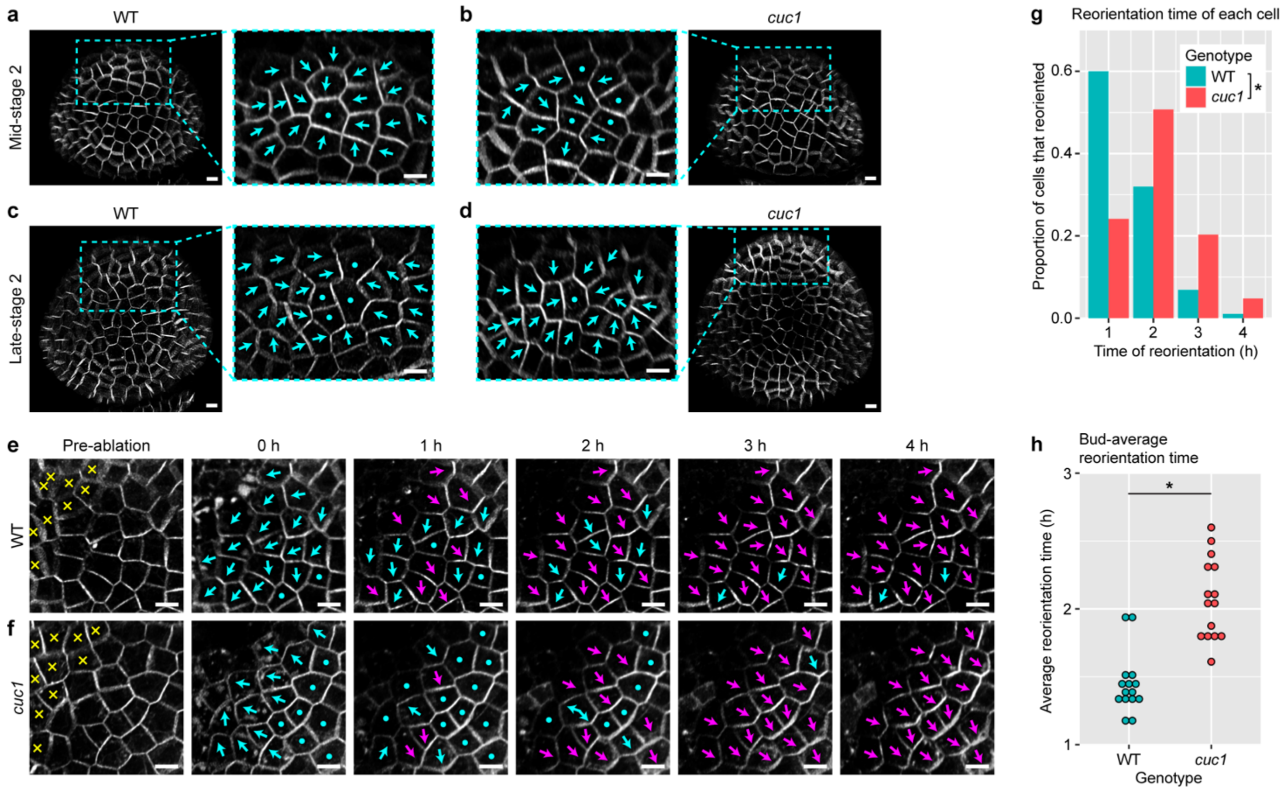


**Supplementary Fig. 6. CUC1 makes PIN1 more dynamic in the floral meristem.**

**(a-d)** PIN1-GFP images of WT **(a,c)** and *cuc1* mutant **(b,d)** buds at mid-stage 2 **(a,b)** and late-stage 2 **(c,d)**. Enlarged images show the incipient outer sepal region. Arrows label PIN1 polarity and dots label cells that are apolar or with unclear polarity. Note that in mid-stage 2, PIN1 is highly polarized towards the incipient outer sepal in WT, which is less clear in *cuc1*. In late-stage 2, PIN1 in both genotypes strongly polarize towards the incipient outer sepal. Scale bars, 5 µm.

**(e-h)** Reorientation of PIN1-GFP in response to ablation. **(e,f)** Representative WT **(e)** and *cuc1* mutant **(f)** buds with PIN1-GFP reporter that are ablated (yellow crosses) and live imaged every hour. Cyan arrows and dots label PIN1 polarity in cells that have not completed reorientation. Magenta arrows label PIN1 polarity in cells that have completed reorientation. Note that cells in WT complete reorientation faster than cells in *cuc1*. This is most obvious in the 1 h image. In this particular WT sample, there are two cells that did not reorient at 4 h (cyan arrows). Scale bars, 5 µm. **(g)** Quantification of reorientation timing in all cells. n = 478 for WT and n = 418 for *cuc1*. Asterisk represents statistical significance from Kolmogorov-Smirnov Test ($p < 2.2\times10^{-16}$). **(h)** Quantification of reorientation time averaged across all cells (for cells that reoriented within 4 h) in each bud, for buds that have at least 10 cells reoriented. Each dot represents a bud (n = 15 for both WT and *cuc1*). Asterisk represents statistically significant difference in a two-sided Student's t-test ($p = 5.788\times10^{-7}$).

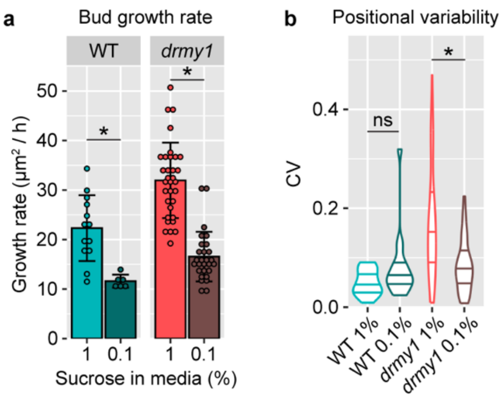


**Supplementary Fig. 7. Additional quantification for WT and *drmy1* buds cultured on normal or low sucrose media.**

**(a)** Culturing inflorescence samples on media containing 0.1% sucrose reduces bud growth rate by half compared to normal growth media containing 1% sucrose. Growth rate is measured as the areal increase in Z-projection ($\mu m^2$) divided by the duration of live imaging (h). Mean ± SD and individual data points were plotted. Sample size: WT 1%, n = 13; WT 0.1%, n = 6; *drmy1* 1%, n = 33; *drmy1* 0.1%, n = 28. Asterisks show statistically significant difference in Student's two-sided t tests (WT, $p = 6.908\times10^{-5}$; drmy1, $p = 3.703\times10^{-13}$). **(b)** Positional variability of sepal primordia initiated. Sample size: WT 1%, n = 22; WT 0.1%, n = 17; *drmy1* 1%, n = 63; *drmy1* 0.1%, n = 43. Asterisks indicate statistically significant differences in Wilcoxon's rank sum test ($p = 3.465\times10^{-5}$) and ns indicates no significant difference (p = 0.1032).

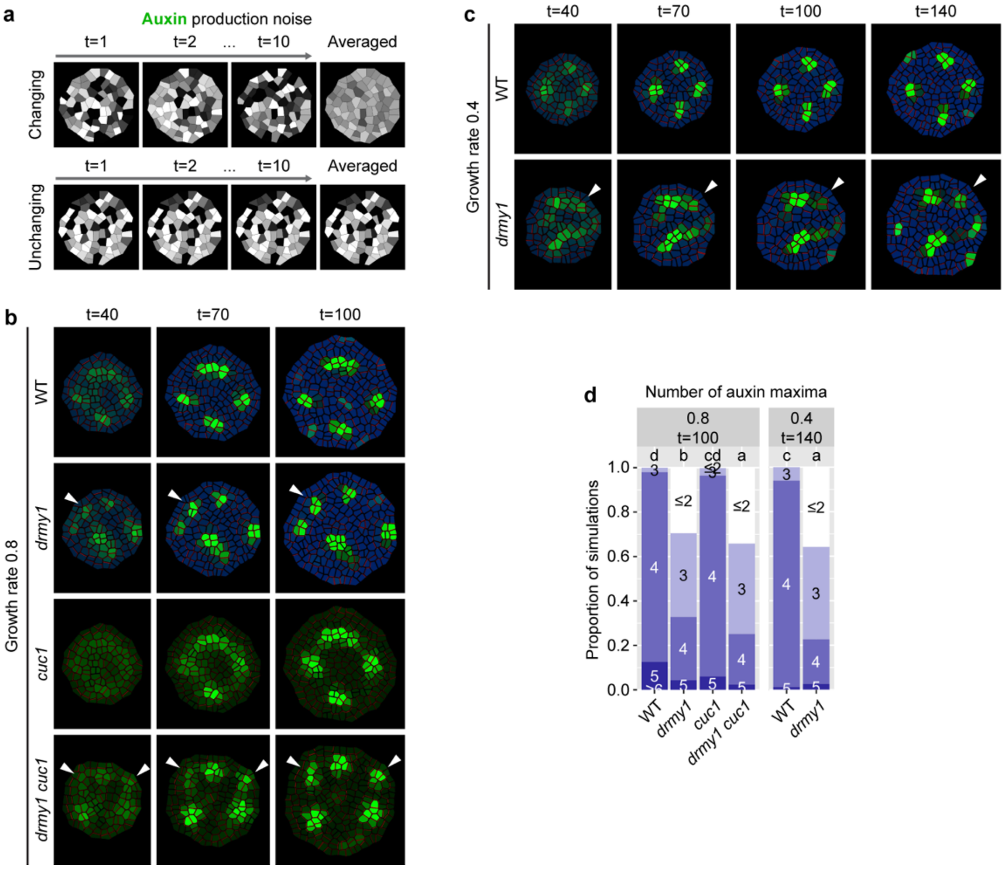


**Supplementary Fig. 8. The *cuc1* mutation and slow growth rescues auxin patterning in *drmy1* by promoting temporal noise averaging.**

**(a)** Temporal noise averaging. When PIN is less sensitive to auxin level fluctuations among neighboring cells, PIN repolarizes on a longer time scale, allowing auxin level fluctuations in each cell to average out through time, which rescues *drmy1*. Setting auxin production noise as temporally unchanging eliminates this averaging, and thus decreasing PIN1 repolarization should no longer rescue *drmy1*. **(b-c)** When noise is set temporally unchanging, in *drmy1* and *drmy1 cuc1* under moderate growth rate **(b)**, and *drmy1* under reduced growth rate **(c)**, sporadic auxin patches are stabilized to form auxin maxima (arrowheads). **(d)** Quantification of auxin maxima number across n=500 simulations. t=100 for growth rate 0.8 (compare to Fig. 6i). t=140 for growth rate 0.4 (compare to Fig. 7f). Letters show multiple comparison of mean auxin maxima number using Tukey's HSD.
