## Supplementary Information Legends for "Tradeoff Between Speed and Robustness in Primordium Initiation Mediated by Auxin-CUC1 Interaction"

**Supplementary Movie 1. *CUC1* expression pattern in WT vs *drmy1* buds.** Related to Fig. 1.
Shown are *pCUC1::3xVENUS-N7* (yellow, top row) and VENUS merged with the Chlorophyll channel (magenta, bottom). Time points are 6 hours apart. Scale bar, 25 µm.

**Supplementary Movie 2. CUC1 protein accumulation pattern in WT vs *drmy1* buds.** Related to Fig. 1.

Shown are *pCUC1::CUC1-GFP* (yellow, top row) and GFP merged with the Chlorophyll channel (magenta, bottom). Time points are 6 hours apart. Scale bar, 25 µm.

**Supplementary Movie 3. Auxin signaling pattern in a WT bud.** Related to Fig. 4.

Shown are *DR5::3xVENUS-N7* (cyan, left) and DR5 merged with the Chlorophyll channel (magenta, right). Time points are normalized relative to the beginning of stage 2 (0 h) and are 6 hours apart. Scale bar, 25 µm. Same for Supplementary Movies 4-7.

**Supplementary Movie 4. Auxin signaling pattern in a *5mCUC1* bud.** Related to Fig. 4.

**Supplementary Movie 5. Auxin signaling pattern in a *cuc1* bud.** Related to Fig. 4.

**Supplementary Movie 6. Auxin signaling pattern in a *drmy1* bud.** Related to Fig. 4.

**Supplementary Movie 7. Auxin signaling pattern in a *drmy1 cuc1* bud.** Related to Fig. 4.

**Supplementary Movie 8. Model of auxin pattern formation in WT, *cuc1*, *drmy1*, and *drmy1 cuc1* under moderate growth rate (0.8).** Related to Fig. 6.

**Supplementary Movie 9. Model of auxin pattern formation in WT and *drmy1* under low (0.4), intermediate (0.8), and high (1.2) growth rate.** Related to Fig. 7.

**Supplementary Movie 10. Model of auxin pattern formation in WT, *cuc1*, *drmy1*, and *drmy1 cuc1* under moderate growth rate (0.8), and WT and *drmy1* under reduced growth rate (0.4), when auxin noise is set constant (temporally unchanging).** Related to Supplementary Fig. 8.

**Supplementary Dataset 1. Data used in all graphs in XLSX format.**

**Supplementary Dataset 2. Computational model of auxin pattern formation in the floral meristem.** Also uploaded on GitHub: https://github.com/RoederLab/MassSpringAuxin
