## Supplementary Dataset 2 for "Tradeoff Between Speed and Robustness in Primordium Initiation Mediated by Auxin-CUC1 Interaction": create_run_20231010_demo.py

```
# Python script for creating the run file
# Shuyao Kong
# 2023-10-10

# Define variables to be tested as lists of strings
CUCs = ["1","0"]
SDauxs = ["0.1","1.0"]
plasts = ["0.4","0.8","1.2"]

# Other variables
initialCond = "/home/ahr75/Documents/Shuyao/oofs_2023-03-18/ext/02-CellModels/ext/04-MassSpring/ext/19-MassSpringAuxinQuadratic/Init_blank_72cells_2023-09-07.mdxm"
pinLowCUC = "Linear"
pinHighCUC= "Quadratic"
maxSteps = "140"
unchangingNoise = "False"
reps = 5

# Get today's date in format YYYYMMDD
import datetime
import os
today = datetime.datetime.today().strftime('%Y%m%d')
outdir = "/home/ahr75/Documents/Shuyao/oofs_output/" + today
if not os.path.exists(outdir):
    os.mkdir(outdir)

# Loop over variables and write file
import sys
f = open("/home/ahr75/Documents/Shuyao/oofs_output/run_"+today+".py", "w")
for rep in range(1,reps+1):
    for CUC in CUCs:
        for SDaux in SDauxs:
            for plast in plasts:
                runName = "Run_"+CUC+"_"+SDaux+"_"+pinLowCUC+"_"+pinHighCUC+"_"+plast+"_"+unchangingNoise+"_sim"+str(rep)+"_"+today
                runName = runName.replace(".","-")
                f.write("Process.Mesh__System__Open('Stack1', '"+initialCond+"', 'no', 'no')\n")
                f.write("Process.Model__CCF__01_Cell_Disk('10', '0.01', '.0001', '10', '10', '10', '0.1', 'Euler', 'Bi-conjugate Gradient Stabilized', '50', '1e-6', 'No', '.0001', 'Model/CCF/02 Mass Spring Solver', 'Model/CCF/10 Cell Tissue', 'Model/CCF/04 Mass Spring Growth', 'Model/CCF/05 Divide Cells', 'Model/CCF/06 Split Edges', 'Model/CCF/Auxin Gradient Model', '"+runName+"', '"+maxSteps+"', '"+outdir+"', '10', 'True', '"+unchangingNoise+"')\n")

f.close()
```
