## Supplementary Dataset 2 for "Tradeoff Between Speed and Robustness in Primordium Initiation Mediated by Auxin-CUC1 Interaction": CellDisk.cpp

```
//
// This file is part of MorphoDynamX - http://www.MorphoDynamX.org
// Copyright (C) 2012-2016 Richard S. Smith and collaborators.
//
// If you use MorphoDynamX in your work, please cite:
//   http://dx.doi.org/10.7554/eLife.05864
//
// MorphoDynamX is free software, and is licensed under under the terms of the 
// GNU General (GPL) Public License version 2.0, http://www.gnu.org/licenses.
//
#include <CellDisk.hpp>
#include <CCUtils.hpp>

namespace CellDisk
{

  // Initialize the process for polar auxin transport
  bool AuxinGradient::initialize(CellTissue &_tissue, CCIndexDataAttr &_indexAttr, MSCellDataAttr &_cellAttr, MSEdgeDataAttr &_msEdgeAttr, QStringList auxParms)
  {
    tissue = &_tissue;
    indexAttr = &_indexAttr;
    cellAttr = &_cellAttr;
    msEdgeAttr = &_msEdgeAttr;

    getParms(auxParms);

    return true;
  }


  //Initialize derivatives for the auxin transport
  void AuxinGradient::initDerivs(SolverT &solver, VertexAttr &vertexAttr)
  {

    for(CCIndex c : tissue->dualGraph().vertices()) {
      auto &cCD = (*cellAttr)[c];

      // Find perimeter for convenient access
      cCD.perimeter = 0;
      for(const auto &pr : neighborInteriors(tissue->dualGraph(), c)) {
        CCIndex w = pr.second;

        CCIndexData &wIdx = (*indexAttr)[w];
        cCD.perimeter += wIdx.measure;
      }
    }
  }


  // Calculate derivative for the solver
  void AuxinGradient::calcDerivatives(const SolverT &solver, VertexAttr &vertexAttr)
  {
    CCStructure &cs = tissue->cellStructure();
    CCStructure &dg = tissue->dualGraph();

    // For each cell
    for(CCIndex c : dg.vertices()) {
      MassSpringCellData &cCD = (*cellAttr)[c];
      CCIndexData &cIdx = (*indexAttr)[c];
      auto &cSD = vertexAttr[c];

      // Decide the function form of PIN repolarization depending on CUC concentration
      QString pinForm;
      if(cCD.cuc <= thresCUC) pinForm = pinLowCUC;
      else pinForm = pinHighCUC;

      // Preparations for calculating PIN1 distribution: sum up values for all neighboring cells
      double aTotal = 0.0;
      // Get the neighbor vertices and the edges (interiors) between
      for(const auto &pr : neighborInteriors(tissue->dualGraph(), c)) {  // c is the focal cell itself
        CCIndex n = pr.first;  // n is the neighboring cell
        CCIndex w = pr.second; // w is the cell wall in between
        CCSignedIndex e = orientedMembrane(cs, c, n); // Going from the focal cell to the neighboring cell
        CCIndexData &wIdx = (*indexAttr)[w];
        MassSpringCellData &nCD = (*cellAttr)[n];

        // Calculate the auxin factor depending on the repolarization function form
        double auxfct = nCD.aux; // Default is linear
        if(pinForm=="Quadratic") auxfct = pow(auxfct,2.0);
        else if(pinForm=="Cubic")auxfct = pow(auxfct,3.0);
        else if(pinForm=="Exp2") auxfct = pow(2.0,auxfct);
        else if(pinForm=="Exp3") auxfct = pow(3.0,auxfct);

        // Sum up the values for all neighboring cells
        if(e.orientation() == ccf::POS) aTotal += auxfct * wIdx.measure;
        else if (e.orientation() == ccf::NEG) aTotal += auxfct * wIdx.measure;
      }

      // Auxin production with noise
      cSD.dx[0] += auxProd * cCD.auxProdNoise;

      // CUC production determined by auxin concentration. Hill coefficient = 4
      cSD.dx[1] += cucProd / (1.0 + pow(cCD.aux/Kaux, 4));

      // Auxin decay
      cSD.dx[0] -= auxDecay * cCD.aux;

      // CUC decay
      cSD.dx[1] -= cucDecay * cCD.cuc;
  
      // Polar auxin transport
      for(const auto &pr : neighborInteriors(tissue->dualGraph(), c)) {

        CCIndex n = pr.first; // Neighboring cell
        CCIndex w = pr.second; // Wall in between (edge in the dual graph)
        MassSpringCellData &nCD = (*cellAttr)[n];
        auto &nSD = vertexAttr[n];
        CCIndexData &wIdx = (*indexAttr)[w];
        CCIndexData &nIdx = (*indexAttr)[n];
  
        if(aTotal <= 0)
          continue;
        CCSignedIndex e = orientedMembrane(cs, c, n); // Going from the focal cell to the neighboring cell
        MassSpringEdgeData &eED = (*msEdgeAttr)[~e];

        // Decide the function form of PIN repolarization depending on CUC concentration
        QString pinForm;
        if(cCD.cuc <= thresCUC) pinForm = pinLowCUC;
        else pinForm = pinHighCUC;
        double auxfct = nCD.aux; // Default is linear
        if(pinForm=="Quadratic") auxfct = pow(auxfct,2.0);
        else if(pinForm=="Cubic")auxfct = pow(auxfct,3.0);
        else if(pinForm=="Exp2") auxfct = pow(2.0,auxfct);
        else if(pinForm=="Exp3") auxfct = pow(3.0,auxfct);
        
        if(e.orientation() == ccf::POS){ // If it is positive direction

          // Determine the amount of PIN1 on this wall
          eED.pinP = (1.0-pinrepr) * eED.pinP + pinrepr * auxfct * wIdx.measure / aTotal;

          // Polar auxin transport
          cSD.dx[0] -= auxTrans * cCD.aux * eED.pinP / cIdx.measure;
          nSD.dx[0] += auxTrans * cCD.aux * eED.pinP / nIdx.measure;

	      }
	      else if(e.orientation() == ccf::NEG){ // If it is negative direction

          // Determine the amount of PIN1 on this wall
          eED.pinN = (1.0-pinrepr) * eED.pinN + pinrepr * auxfct * wIdx.measure / aTotal;

          // Polar auxin transport
          cSD.dx[0] -= auxTrans * cCD.aux * eED.pinN / cIdx.measure;
          nSD.dx[0] += auxTrans * cCD.aux * eED.pinN / nIdx.measure;
	      }
      }
    }
  }


  // Update noise in auxin production and PIN1 distribution
  void AuxinGradient::UpdateNoise() {

    CCStructure &dg = tissue->dualGraph();

    // Attrs for visualization
    CCIndexDoubleAttr &auxProdNoiseAttr = currentMesh()->signalAttr<double>("NoiseAuxProd");

    // Draw an auxin production rate for each cell from a Gaussian distribution
    for(CCIndex f : dg.vertices()){
      MassSpringCellData &pCD = (*cellAttr)[f];
      pCD.auxProdNoise = max(0.0, gaussRan(1.0, sdAuxinProd)); // Capped on the lower end (0) but uncapped on the higher end
      auxProdNoiseAttr[f] = pCD.auxProdNoise; // Update Attr for visualization
    }
  }


  // Calculate cell area
  void AuxinGradient::CalcCellArea(){
    for(CCIndex c : tissue->dualGraph().vertices()) {
      auto &cIdx = (*indexAttr)[c];
      auto &cCD = (*cellAttr)[c];
      cCD.area = cIdx.measure;
    }
  }


  // Dilute auxin and CUC based on cell growth
  void AuxinGradient::Dilute(){
    for(CCIndex c : tissue->dualGraph().vertices()) {
      auto &cIdx = (*indexAttr)[c];
      auto &cCD = (*cellAttr)[c];
      cCD.aux = cCD.aux * cCD.area / cIdx.measure;
      cCD.cuc = cCD.cuc * cCD.area / cIdx.measure;
    }
  }


  // Initialization of the main process
  bool CellDisk::initialize(QWidget *parent)
  {
  
    // Initialize the random number generator
    sran(time(0));

    // Get run control parameters
    runName = parm("Run Name");
    runMaxSteps = parm("Run Max Steps");
    outDir = parm("Output directory");
    outFreq = parm("Output frequency");
    stepCount = 0;

    // Get the current mesh
    mesh = currentMesh();
    if(!mesh) throw(QString("CellDisk::initialize No current mesh"));
      
    indexAttr = &mesh->indexAttr();
    cellAttr = &mesh->attributes().attrMap<CCIndex,MassSpringCellData>("DiskAuxinCellData");
    msEdgeAttr = &mesh->attributes().attrMap<CCIndex,MassSpringEdgeData>("Mass Spring Data");

    // Cell Division
    if(!getProcess(parm("Divide Process"), divideProcess))
      throw(QString("CellDisk::initialize Cannot make divide cell process"));
    divideProcess->initialize(parent);

    // Growth Process
    if(!getProcess(parm("Growth Process"), growthProcess))
      throw(QString("CellDisk::initialize Cannot make growth process"));
    growthProcess->initialize(parent);

    // Tissue process, growth process initializes
    if(!getProcess(parm("Tissue Process"), tissueProcess))
      throw(QString("CellDisk::initialize Cannot make tissue process"));
    tissueProcess->initialize(parent);
    
    // AuxinGradient: Initialize the solver derivatives provider
    if(!getProcess(parm("Auxin Gradient Model"), auxinProcess)){
      throw(QString("CellDiskAuxin::initialize Unable to make Auxin Gradient Model process"));
    }
    QStringList auxParms = QStringList();
    if(parm("Autorun")=="True"){
      auxParms = runName.replace("-",".").split('_').mid(1,5);
    }
    auxinProcess->initialize(tissue(), *indexAttr, *cellAttr, *msEdgeAttr, auxParms);

    // Initialize AuxinGradient
    clearDerivs();
    addDerivs(auxinProcess);
    auxinProcess->UpdateNoise();
    auxinProcess->CalcCellArea();
    
    // Mass-Spring: Initialize the solver process
    if(!getProcess(parm("Solver Process"), solverProcess))
      throw(QString("CellDisk::initialize Cannot make mass-spring solver process"));
    solverProcess->initialize(parent);

    // Get the split edges process
    if(!getProcess(parm("Split Edges Process"), splitEdgesProcess))
      throw(QString("CellDisk::initialize Cannot make splitEdges process"));

    // Create directory based on today's date
    QDir dirdate(outDir);
    if(!dirdate.exists()) dirdate.mkpath(".");
    QDir::setCurrent(outDir);

    // Create sub-directories in the output directory
    QDir dirrun(runName);
    if(dirrun.exists()) dirrun.removeRecursively(); // override if existing
    dirrun.mkpath(".");
    dirrun.mkpath("./Screenshots");
    dirrun.mkpath("./CCData");

    // Display "Faces > Polarity" to ease screenshots
    CCDrawParms &cdp = mesh->drawParms(mesh->ccName());
    cdp.setRenderChoice("Faces", "Polarity");
    cdp.setGroupVisible("Faces", true);
    updateState();
    updateViewer();

    return true;
  }


  bool CellDisk::step()
  {
    // Mass spring, return if not converged.
    // This means the program runs the Mass-Spring simulation until it converges, and then run the auxin, growth, and division simulations
    solverProcess->solve();
    double xNorm = solverProcess->calcXNorm();
    double dxNorm = solverProcess->calcDXNorm();
    if(dxNorm < parm("Converge Threshold").toDouble()){
      mdxInfo << "Converged, X Norm:" << xNorm << " Dx Norm:" << dxNorm << endl;
    }
    else{
      mdxInfo << "Mass spring did not converge, Dx Norm: " << dxNorm << endl;
      return true;
    }
    stepCount++;

    // Dilute morphogens because of cell growth
    auxinProcess->Dilute();

    // Update noise of auxin production, unless noise is set to be unchanging
    if(parm("Unchanging Noise")=="False") auxinProcess->UpdateNoise();

    // Initialize and solve the auxin system
    initSolver(&tissue().dualGraph());
    solve();
    mesh->updateProperties(tissue().tissueName());
    mesh->updateProperties(tissue().tissueDualName());

    // Calculate Attrs for rendering
    displayAttrs();

    // Split large-enough cells
    divideProcess->step();

    // Split edges
    splitEdgesProcess->run();

    // Grow
    growthProcess->step();

    // Re-initialize the solvers for MassSpring and AuxinGradient
    solverProcess->initialize(0);
    QStringList auxParms = runName.replace("-",".").split('_').mid(1,5);
    auxinProcess->initialize(tissue(), *indexAttr, *cellAttr, *msEdgeAttr, auxParms);
    mesh->updateAll(tissueProcess->tissue().tissueName());
    mesh->updateAll(tissueProcess->tissue().tissueDualName());

    // Re-calculate cell area after growth and division
    auxinProcess->CalcCellArea();

    // Output cell data and screenshot
    if(outFreq != "")
      if(stepCount % outFreq.toInt() == 0) outputToFile();

    // Check stopping condition
    return checkStop();

  }


  bool CellDisk::rewind(QWidget *parent)
  {
    // To rewind, we'll reload the mesh
    mesh = currentMesh();
    if(!mesh or mesh->file().isEmpty())
      throw(QString("No current mesh, cannot rewind"));
    MeshLoad meshLoad(*this);
    meshLoad.setParm("File Name", mesh->file());
    return meshLoad.run();
  }


  // Output cell data and screenshot
  void CellDisk::outputToFile(){

    // Take a screenshot
    QDir::setCurrent(outDir + "/" + runName + "/Screenshots");
    takeSnapshot("Screenshot_" + runName + "_" + QString::number(stepCount) + ".jpg", {1320,1320});
    mdxInfo << ">>>>>> Screenshot saved in:" << QDir::current().absolutePath() << endl;

    // Output Cell Complex Data
    QDir::setCurrent(outDir + "/" + runName + "/CCData");
    QFile CCDataFile("CCData_" + runName + "_" + QString::number(stepCount) + ".txt");
    if (CCDataFile.open(QIODevice::ReadWrite)) {
      QTextStream stm(&CCDataFile);

      // Access cell complex Attrs
      CCIndexDataAttr &ccAttr = mesh->indexAttr();
      CCStructure &cs = tissue().cellStructure();

      // Create header
      stm   << "CCIndex" <<"\t"<< "x"          <<"\t"<< "y"          <<"\t"<< "area"        <<"\t"<< "aux"   <<"\t"<< "cuc"   << endl;
      for(CCIndex f : cs.faces()) {
        QString fIndex = QString::fromStdString(ccf::to_string(f));
        auto fAttr = ccAttr[f];
        MassSpringCellData &fCD = (*cellAttr)[f];
        stm << fIndex    <<"\t"<< fAttr.pos[0] <<"\t"<< fAttr.pos[1] <<"\t"<< fAttr.measure <<"\t"<< fCD.aux <<"\t"<< fCD.cuc << endl; 
      }
    }
    mdxInfo << ">>>> " << runName << " step " << stepCount << " saved in" << QDir::current().absolutePath() << endl;

  }


  // Check stopping condition; if stop conditions are met, return false to terminate the program
  bool CellDisk::checkStop(){
    if(runMaxSteps != ""){
      if(stepCount >= runMaxSteps.toInt()) return false;
    }
    return true;
  }


  void CellDisk::displayAttrs(){

    // Update the edge attributes for MDX rendering
    CCSignedIndexDoubleAttr &polAttr = currentMesh()->attributes().attrMap<CCSignedIndex, double>("Polarity");
    
    const CCStructure &cs = tissue().cellStructure();
    for(CCIndex f : cs.faces()) {

      // Update PIN1 for display
      for(CCIndex e : cs.bounds(f)) {
        CCSignedIndex es(e, cs.ro(f, e));

        if(es.orientation() == ccf::POS){
          polAttr[es] = (*msEdgeAttr)[e].pinP;
        }
        else if(es.orientation() == ccf::NEG){
          polAttr[es] = (*msEdgeAttr)[e].pinN;
        }
      }
    }
  }


  bool CellDiskSolver::initialize(QWidget *parent)
  {
    // Mass spring process
    MassSpringDerivs *msDerivs;
    if(!getProcess(parm("Mass Spring Process"), msDerivs))
      throw QString("CellDiskSolver::initialize Cannot make mass-spring derivs process %2").arg(name()).arg(parm("Mass Spring Process"));
    msDerivs->initialize(parent);

    Mesh *mesh = currentMesh();
    msDerivs->cs = &mesh->ccStructure("Tissue");
    msDerivs->indexAttr = &mesh->indexAttr();

    clearDerivs();
    addDerivs(msDerivs);
    initSolver(msDerivs->cs);

    return true;
  }


  void MassSpringDerivs::initDerivs(Solver<Point3d> &solver, Solver<Point3d>::VertexAttr &vAttr, Solver<Point3d>::EdgeAttr &eAttr)
  {
    Mesh *mesh = currentMesh();
    if(!mesh)
      throw(QString("MassSpringDerivs::initDerivs No current mesh"));

    // Get parameters from GUI
    k = parm("Spring Constant").toDouble();
    pressure = parm("Pressure").toDouble();

    cs = &mesh->ccStructure("Tissue");
    indexAttr  = &mesh->indexAttr();
    msEdgeAttr = &mesh->attributes().attrMap<CCIndex, MassSpringEdgeData>("Mass Spring Data");

    // Locate border and mark it, also grab cell for orientation
    auto edges = cs->edges();

    #pragma omp parallel for
    for(uint i = 0; i < edges.size(); i++) {
      CCIndex e = edges[i];
      MassSpringEdgeData &eMS = (*msEdgeAttr)[e];

      auto cb = cs->cobounds(e);
      if(cb.size() == 1) { // On the border
        eMS.border = true;
        eMS.f = *cb.begin();
      } else {
        eMS.border = false;
        eMS.f = CCIndex::UNDEF;
      }
    }
  }


  void MassSpringDerivs::calcDerivatives(const SolverT &solver, CCIndex v, Point3d &values)
  {
    if(!cs)
      throw QString("MassSpringDerivs::calcDerivatives Cell complex not set").arg(name());
    if(!indexAttr)
      throw QString("MassSpringDerivs::calcDerivatives Index data attribute not set").arg(name());
    if(!msEdgeAttr)
      throw QString("MassSpringDerivs::calcDerivatives Edge data attribute not set").arg(name());

    CCIndexData &vIdx = (*indexAttr)[v];
    for(const Flip &flip : cs->matchV(CCIndex::BOTTOM, v, CCIndex::Q, CCIndex::Q)) {
      CCIndex e = flip.interior;
      CCIndex n = flip.otherFacet(v);
      CCIndexData &nIdx = (*indexAttr)[n];

      MassSpringEdgeData &eMS = (*msEdgeAttr)[e];
      Point3d dir =  nIdx.pos - vIdx.pos;
      double length = norm(dir);
      if(length > 0)
        dir /= length;
      values += (length - eMS.restLen)/eMS.restLen * k/eMS.elast * dir;

      // Pressure only on the outside edges
      if(eMS.border and pressure > 0) {
        // Find orientation
        int ro = cs->ro(eMS.f, e) * cs->ro(e, v);
        // Apply pressure
        values += Point3d(dir.y(), -dir.x(), 0) * (ro == ccf::POS ? 1.0 : -1.0) * pressure * length * 0.25; // 1/2 pressure applied twice
      }
    }
  }


  bool CellDiskGrowth::initialize(QWidget *parent)
  {
    // Get the current mesh
    mesh = currentMesh();
    if(!mesh) throw(QString("CellDisk::initialize No current mesh"));

    // Initialize the tissue
    if(!getProcess(parm("Tissue Process"), tissueProcess))
      throw(QString("CellDiskGrowth::initialize Cannot make tissue process"));
    tissueProcess->initialize(parent);

    return true;
  }


  bool CellDiskGrowth::step()
  {
    if(!mesh) throw(QString("CellDiskGrowth::step No current mesh"));

    if(!tissueProcess) throw(QString("CellDiskGrowth::step No tissue process"));

    CCStructure &cs = tissueProcess->tissue().cellStructure();
    auto &msEdgeAttr = mesh->attributes().attrMap<CCIndex, MassSpringEdgeData>("Mass Spring Data");
    auto &indexAttr = mesh->indexAttr();

    double growthRate = parm("Growth Rate").toDouble();
    double dt = parm("Dt").toDouble();

    auto edges = cs.edges();

    // Growth
    for(uint i = 0; i < edges.size(); i++) {
      CCIndex e = edges[i];
      MassSpringEdgeData &eMS = msEdgeAttr[e];
      CCIndexPair ep = cs.edgeBounds(e);
      CCIndexData &vIdx = indexAttr[ep.first];
      CCIndexData &wIdx = indexAttr[ep.second];
      double length = norm(vIdx.pos - wIdx.pos);
      // Strain-based growth
      if(length > eMS.restLen and growthRate > 0) {
        eMS.restLen *= 1.0 + (length - eMS.restLen)/eMS.restLen * growthRate * eMS.plast * dt;
      }
    }

    return true;
  }


  // Initialize to grab subdivider
  bool CellDiskDivide::initialize(QWidget *parent)
  {
    Mesh *mesh = currentMesh();
    if(!mesh)
      throw(QString("CellDiskDivide::step No current mesh"));

    // Call base initialize
    if(!CellTissueCell2dDivide::initialize(parent))
      return false;

    if(!getProcess(parm("Solver Process"), solverProcess))
      throw(QString("CellDiskDivide::initialize Cannot make solver process"));
    solverProcess->initialize(parent);

    if(!getProcess(parm("Tissue Process"), tissueProcess))
      throw(QString("CellDisk::initialize Cannot make tissue process"));

    // Setup subdivision object
    MSEdgeDataAttr *eAttr = &mesh->attributes().attrMap<CCIndex,MassSpringEdgeData>("Mass Spring Data");
    MSCellDataAttr *cAttr = &mesh->attributes().attrMap<CCIndex,MassSpringCellData>("DiskAuxinCellData");
    subdiv.mdx = *CellTissueCell2dDivide::subdivider();
    subdiv.ms = MassSpringSubdivide(eAttr, cAttr, &mesh->indexAttr());

    // Set noise in division (overriding)
    setParm("Center Noise", "1.0");
    setParm("Wall Noise", "1.0");
    setParm("Junction Distance Noise", "1.0");

    return true;
  }


  // Run a step of cell division
  bool CellDiskDivide::step() 
  { 
    return CellTissueCell2dDivide::step(currentMesh(), &subdiv);
  }


  // Function to retrieve the Cell Tissue
  CellTissue &CellDisk::tissue()
  {
    if(!tissueProcess) throw(QString("CellDisk::tissue Tissue process empty"));
    return tissueProcess->tissue();
  }


  // Rendering
  bool CellDiskRender::defaultDrawChoices(Mesh &mesh, const QString &ccName)
  {
    cellAttr = &mesh.attributes().attrMap<CCIndex,MassSpringCellData>("DiskAuxinCellData");

    // Call base method
    RenderPolarity::defaultDrawChoices(mesh, ccName);

    if(!polColors) {
      mdxInfo << "CellDiskRender::defaultDrawChoices Polarity color map empty" << endl;
      return false;
    }

    // Now redo the colorMap if required, default should be 4 colors
    if(polColors->size() < 6) {
      polColors->colors.resize(10);
      polColors->colors[0] = Colorb(  0,  31,   0, 255);
      polColors->colors[1] = Colorb(  0, 255,   0, 255); // Auxin, dark green to green
      polColors->colors[2] = Colorb(  0,   0,   0, 255);
      polColors->colors[3] = Colorb(255,   0,   0, 255); // PIN1, black to red
      polColors->colors[4] = Colorb(  0,   0,   0, 255);
      polColors->colors[5] = Colorb(  0,   0, 255, 255); // CUC1, black to blue

      polColors->makeRangeMap();
      polColors->setName(0, "Auxin");
      polColors->setBounds(0, Point2d(0.0, 20.0));
      polColors->setUnit(0, "");
      polColors->setIndices(0, Point2i(0, 1));
      polColors->setName(1, "PIN");
      polColors->setBounds(1, Point2d(0.0, 0.75));
      polColors->setUnit(1, "");
      polColors->setIndices(1, Point2i(2, 3));
      polColors->setName(2, "CUC");
      polColors->setBounds(2, Point2d(0.0, 10.0));
      polColors->setUnit(2, "");
      polColors->setIndices(2, Point2i(4, 5));
      mdxInfo << "Reset color map complete" << endl;
    } 

    return true;
  }


  // Set the face color based on morphogen concentration
  Colorb CellDiskRender::setFacePolarityColor(CCIndex c, VizAttribute<CCIndex> &auxAttr)
  {
    if(!cellAttr) { // can't throw, called from OMP
      mdxInfo << "CellDiskRender::setFaceSignalColor cellAttr is null" << endl;
      return Colorb();
    }
    MassSpringCellData &cCD = (*cellAttr)[c];
    
    const ColorMap::ChannelMap &cmap1 = polColors->channelMap(0); // Auxin colormap
    const ColorMap::ChannelMap &cmap2 = polColors->channelMap(2); // CUC colormap
    Colorf visCol1 = Colorf(polColors->getRawColor(cCD.aux, cmap1.bounds, cmap1.indices));
    Colorf visCol2 = Colorf(polColors->getRawColor(cCD.cuc, cmap2.bounds, cmap2.indices));

    // composite the colours together in even measure
    Colorf color = composite(visCol1, visCol2, comp::LIGHTEN);

    return Colorb(color);

  }


 // Process to get auxin and CUC concentration
  bool getConc::initialize(QWidget *parent)
  {
    Mesh *mesh = currentMesh();
    if(!mesh) throw(QString("getConc::initialize No current mesh"));

    QString ccName = mesh->ccName();
    if(ccName.isEmpty()) throw(QString("getConc::initialize Error, no cell complex selected"));

    CCStructure &cs = mesh->ccStructure(ccName);
    CCIndexDataAttr &indexAttr = mesh->indexAttr();
    MSCellDataAttr *cellAttr = &mesh->attributes().attrMap<CCIndex,MassSpringCellData>("DiskAuxinCellData");

    for(CCIndex c : cs.faces()) {
      CCIndexData &cIdx = indexAttr[c];
      if(cIdx.selected) {
        auto &cCD = (*cellAttr)[c];
        mdxInfo << "Selected cell has auxin " << cCD.aux << " and CUC " << cCD.cuc << endl;
      }
    }
    return true;
  }
  bool getConc::run()
  {
    return true;
  }


  // Process to set auxin and CUC concentration
  bool setConc::initialize(QWidget *parent)
  {
    AmountAux = parm("Amount Auxin");
    AmountCuc = parm("Amount CUC");
    return true;
  }
  bool setConc::run()
  {
    Mesh *mesh = currentMesh();
    if(!mesh)
      throw(QString("CellGrid::run No current mesh"));

    QString ccName = mesh->ccName();
    if(ccName.isEmpty())
      throw(QString("setConc::run Error, no cell complex selected"));

    CCStructure &cs = mesh->ccStructure(ccName);
    CCIndexDataAttr &indexAttr = mesh->indexAttr();
    MSCellDataAttr *cellAttr = &mesh->attributes().attrMap<CCIndex,MassSpringCellData>("DiskAuxinCellData");

    for(CCIndex c : cs.faces()) {
      CCIndexData &cIdx = indexAttr[c];
      if(cIdx.selected) {
        auto &cCD = (*cellAttr)[c];
        if(!AmountAux.isEmpty()) cCD.aux = AmountAux.toDouble();
        if(!AmountCuc.isEmpty()) cCD.cuc = AmountCuc.toDouble();
        mdxInfo << "Auxin conc set to " << cCD.aux << " and CUC conc set to " << cCD.cuc << endl;
      }
    }
    mesh->updateProperties(ccName);

    return true;
  }


  REGISTER_PROCESS(CellDisk);
  REGISTER_PROCESS(CellDiskDivide);
  REGISTER_PROCESS(CellDiskGrowth);
  REGISTER_PROCESS(CellDiskTissue);
  REGISTER_PROCESS(CellDiskSolver);
  REGISTER_PROCESS(CellDiskSplitEdges);
  REGISTER_PROCESS(MassSpringDerivs);
  
  REGISTER_PROCESS(AuxinGradient);
  REGISTER_PROCESS(CellDiskRender);

  REGISTER_PROCESS(getConc);
  REGISTER_PROCESS(setConc);
    
}
```
