## Supplementary Dataset 2 for "Tradeoff Between Speed and Robustness in Primordium Initiation Mediated by Auxin-CUC1 Interaction": CellDisk.hpp

```
//
// This file is part of MorphoDynamX - http://www.MorphoDynamX.org
// Copyright (C) 2012-2016 Richard S. Smith and collaborators.
//
// If you use MorphoDynamX in your work, please cite:
//   http://dx.doi.org/10.7554/eLife.05864
//
// MorphoDynamX is free software, and is licensed under under the terms of the 
// GNU General (GPL) Public License version 2.0, http://www.gnu.org/licenses.
//
#ifndef CELL_APEX_HPP
#define CELL_APEX_HPP

#include <Attributes.hpp>
#include <Function.hpp>
#include <QDir>

#include <Contour.hpp>

#include <Mesh.hpp>
#include <Process.hpp>
#include <Random.hpp>
#include <MeshProcessSystem.hpp>
#include <MeshProcessSystemRender.hpp>

#include <Solver.hpp>

#include <MDXProcessTissue.hpp>
#include <MDXProcessCellDivide.hpp>
#include <MeshProcessStructure.hpp>

using namespace mdx;

namespace CellDisk
{
  class CellDiskAuxin;
  class CellDiskDivide;
  class CellDiskTissue;
  class CellDiskGrowth;
  class CellDiskSolver;
  class CellDiskSplitEdges;
  class MassSpringDerivs;
  

  // Structure to store the cell data
  struct MassSpringCellData
  { 
    double aux = 0; // Auxin concentration
    double auxProdNoise = 1.0;// Noise in auxin production, as a multiplicative factor of Gaussian distribution
    double cuc = 0; // CUC1 concentration
    double perimeter = 0;
    double area = 0;

    // Unused
    double amount = 0;
    uint cellType = 0;
    bool fixedConc = false;

    MassSpringCellData() {}

    bool operator==(const MassSpringCellData &other) const
    {
      if(aux == other.aux and auxProdNoise == other.auxProdNoise and cuc == other.cuc
          and perimeter == other.perimeter 
          and area == other.area)
        return true;
      return false;
    }

  };
  typedef AttrMap<CCIndex,MassSpringCellData> MSCellDataAttr;
  
  
  // Structure to store the edge data
  struct MassSpringEdgeData
  {
    double restLen; // Rest length
    CCIndex f; // Cell if edge is on border for quick lookup
    bool border; // Is edge on the border?
    double pinP; // Total amount of PIN1 localized to this wall, in the positive direction (Note: not concentration)
    double pinN; // Total amount of PIN1 localized to this wall, in the negative direction (Note: not concentration)
    double pinPNoise; // Unused
    double pinNNoise; // Unused
    double plast; // Plasticity of the cell wall
    double elast; // Elasticity of the cell wall

    MassSpringEdgeData() : 
      restLen(1.0), f(0), border(false), pinP(0.0), pinN(0.0), pinPNoise(1.0), pinNNoise(1.0), plast(1.0), elast(1.0) {}

    bool operator==(const MassSpringEdgeData &other)
    {
      if(restLen == other.restLen and f == other.f and border == other.border
        and pinP == other.pinP and pinN == other.pinN
        and plast == other.plast and elast == other.elast)
        return true;
      return false;
    }

  };
  typedef AttrMap<CCIndex,MassSpringEdgeData> MSEdgeDataAttr;
  
  
  // Process defining the differential equations of molecular interactions (auxin and CUC)
  class AuxinGradient : public Process, public SolverDerivs<Point2d>
  {
  public:
 
    // Constructor
    AuxinGradient(const Process &process) : Process(process) 
    {
      setName("Model/CCF/Auxin Gradient Model");
      setDesc("Transport on a cellular grid");
      setIcon(QIcon(":/images/CellGrid.png"));

      // Auxin dynamics parameters
      addParm("Auxin Production", "Mean auxin production per cell averaged through time", "1.0");
      addParm("Auxin Transport", "Polar auxin transport coefficient", "400.0");
      addParm("Auxin Decay", "Auxin decay coefficient", "0.2");
      addParm("SD Auxin Production", "Gaussian model: SD of auxin production", "0.1");

      // CUC dynamics parameters
      addParm("CUC Prod", "CUC Production coefficient", "1.0");
      addParm("CUC Decay", "CUC decay coefficient", "0.2");

      // Auxin-CUC interaction parameters
      addParm("K auxin", "Half saturation level of auxin on repressing CUC production", "5.0");
      addParm("Thres CUC", "Threshold of CUC, where PIN1 changes repolarization function form", "2.0");
      addParm("PIN low CUC", "Repolarization function form of PIN1 when CUC is below the threshold", "Linear", QStringList() << "Linear" << "Quadratic" << "Cubic" << "Exp2" << "Exp3");
      addParm("PIN high CUC", "Repolarization function form of PIN1 when CUC is above the threshold", "Quadratic", QStringList() << "Linear" << "Quadratic" << "Cubic" << "Exp2" << "Exp3");
      addParm("PIN repolarize speed", "Repolarization speed of PIN1 in the presence of CUC1", "0.01");

      // Growth parameters
      addParm("Plasticity", "Baseline plasticity", "0.8");
      addParm("Elasticity", "Baseline elasticity", "0.8");

    }
    using Process::initialize;
    bool initialize(CellTissue &tissue, CCIndexDataAttr &indexAttr, MSCellDataAttr &cellAttr, MSEdgeDataAttr &msEdgeAttr, QStringList auxParms);

    // Reimplemented solver methods, Jacobian is tricky for up-the-gradient PIN
    void initDerivs(SolverT &solver, VertexAttr &vertexData);
    void calcDerivatives(const SolverT &solver, VertexAttr &vertexData);

    void setValues(const SolverT &solver, CCIndex c, const VectorT &values)
    {
      MassSpringCellData &cCD = (*cellAttr)[c];
      cCD.aux = values[0];
      cCD.cuc = values[1];
    }
  
    void getValues(const SolverT &solver, CCIndex c, VectorT &values)
    {
      MassSpringCellData &cCD = (*cellAttr)[c];
      values[0] = cCD.aux;
      values[1] = cCD.cuc;
    }

    void getParms(QStringList auxParms)
    {
      // Auxin dynamics parameters
      auxProd = parm("Auxin Production").toDouble();
      auxTrans = parm("Auxin Transport").toDouble();
      auxDecay = parm("Auxin Decay").toDouble();

      // CUC dynamics parameters
      cucDecay = parm("CUC Decay").toDouble();

      // Growth parameters
      elast = parm("Elasticity").toDouble();

      // Auxin-CUC interaction parameters
      Kaux = parm("K auxin").toDouble();
      thresCUC = parm("Thres CUC").toDouble();
      pinrepr = parm("PIN repolarize speed").toDouble();

      // Autorun parameters
      if(auxParms.empty()){ // Not autorun
        cucProd = parm("CUC Prod").toDouble();
        sdAuxinProd = parm("SD Auxin Production").toDouble();
        pinLowCUC = parm("PIN low CUC");
        pinHighCUC = parm("PIN high CUC");
        plast = parm("Plasticity").toDouble();
      }
      else{ 
        // Autorun, run parameters are given by auxParms
        // CUC Prod, SD aux, SD PIN, PIN low CUC, PIN high CUC, plasticity
        cucProd = auxParms[0].toDouble();
        sdAuxinProd = auxParms[1].toDouble();
        pinLowCUC = auxParms[2];
        pinHighCUC = auxParms[3];
        plast = auxParms[4].toDouble();
      }

      // Update plasticity and elasticity for all cells (the same value)
      CCStructure &cs = tissue->cellStructure();
      for(CCIndex e : cs.edges()){
        MassSpringEdgeData &pED = (*msEdgeAttr)[e];
        pED.plast = plast;
        pED.elast = elast;
      }

    }

    void UpdateNoise(); // Update noise in auxin production and PIN1 distribution

    void CalcCellArea(); // Calculate cell area to determine how much auxin and CUC are diluted because of cell expansion

    void Dilute(); // Dilute auxin and CUC because of cell expansion

  private:
    CellTissue *tissue = 0;
    CCIndexDataAttr *indexAttr = 0;
    MSCellDataAttr *cellAttr = 0;
    MSEdgeDataAttr *msEdgeAttr = 0;

    // Parameters for auxin dynamics
    double auxProd = 0.0;
    double auxTrans = 0.0;
    double auxDecay = 0.0;
    double sdAuxinProd = 0.0;

    // Parameters for CUC dynamics
    double cucProd = 0.0;
    double cucDecay = 0.0;

    // Parameters for auxin-CUC interaction
    double Kaux = 0.0; // Half saturation level of auxin, for both growth and CUC expression
    double thresCUC = 0.0;
    QString pinLowCUC = "Linear";
    QString pinHighCUC = "Quadratic";
    double pinrepr = 0.01;

    // Parameters for growth
    double plast = 0.8; // Baseline plasticity
    double elast = 0.8; // Baseline elasticity

  };


  // Class for subdivision of cells
  class MassSpringSubdivide : virtual public Subdivide
  {
  public:
    MassSpringSubdivide() {}
    MassSpringSubdivide(MSEdgeDataAttr *msEdgeAttr, MSCellDataAttr *msCellAttr, CCIndexDataAttr *indexAttr) 
     : msEdgeData(msEdgeAttr), msCellData(msCellAttr), indexAttr(indexAttr) {} // Initializer list
    void splitCellUpdate(Dimension dim, const CCStructure &cs, const CCStructure::SplitStruct& ss,
                   CCIndex otherP = CCIndex(), CCIndex otherN = CCIndex(), double interpPos = 0.5) const
    {

      if(dim == 1) {
      
        // Propagate rest length
        MassSpringEdgeData &rMS = (*msEdgeData)[ss.parent];
        MassSpringEdgeData &pMS = (*msEdgeData)[ss.childP];
        MassSpringEdgeData &nMS = (*msEdgeData)[ss.childN];

        nMS.restLen = (1.0 - interpPos) * rMS.restLen;
        pMS.restLen = interpPos * rMS.restLen;
        
        // Inherit PIN amount, plasticity, and elasticity
        nMS.pinP = (1.0 - interpPos) * rMS.pinP;
        nMS.pinN = (1.0 - interpPos) * rMS.pinN;
        nMS.plast = rMS.plast;
        nMS.elast = rMS.elast;
        pMS.pinP = interpPos * rMS.pinP;
        pMS.pinN = interpPos * rMS.pinN;
        pMS.plast = rMS.plast;
        pMS.elast = rMS.elast;
        
      } else if(dim  == 2) {
      
        // Assign rest length to new wall, just use actual length
        MassSpringEdgeData &eMS = (*msEdgeData)[ss.membrane];
        CCIndexPair ep = cs.edgeBounds(ss.membrane);
        eMS.restLen = norm((*indexAttr)[ep.first].pos - (*indexAttr)[ep.second].pos);
        
        // Initialize PIN concentration, plasticity, and elasticity
        eMS.pinP = 0.01; // cannot be set to 0
        eMS.pinN = 0.01; // cannot be set to 0
        eMS.plast = 1.0;
        eMS.elast = 1.0;
        
        // The two childs inherit the same Auxin concentration, CUC concentration, and noise
        MassSpringCellData &cR = (*msCellData)[ss.parent];
        MassSpringCellData &cP = (*msCellData)[ss.childP];
        MassSpringCellData &cN = (*msCellData)[ss.childN];
        cP.aux = cR.aux;
        cN.aux = cR.aux;
        cP.auxProdNoise = cR.auxProdNoise;
        cN.auxProdNoise = cR.auxProdNoise;
        cP.cuc = cR.cuc;
        cN.cuc = cR.cuc;
      }
    }

    MSEdgeDataAttr *msEdgeData;
    MSCellDataAttr *msCellData;
    CCIndexDataAttr *indexAttr;
  };

  class CellDiskSubdivide : virtual public Subdivide
  {
  public:
    CellDiskSubdivide() {}

    void splitCellUpdate(Dimension dim, const CCStructure &cs, const CCStructure::SplitStruct& ss, 
        CCIndex otherP = CCIndex(), CCIndex otherN = CCIndex(),  double interpPos = 0.5)
    {
      mdx.splitCellUpdate(dim, cs, ss, otherP, otherN, interpPos);
      ms.splitCellUpdate(dim, cs, ss, otherP, otherN, interpPos);
    }

    MDXSubdivide mdx;
    MassSpringSubdivide ms;
  };
 
 
 
  // Main process (MassSpringCells merged with GridAuxin)
  class CellDisk : public Process, public Solver<Point2d>
  {
  public:
    CellDisk(const Process &process) : Process(process) 
    {
      setName("Model/CCF/01 Cell Disk");
      setDesc("Growing cellular apex");
      setIcon(QIcon(":/images/CellDisk.png"));
      addParm("Converge Threshold", "Convergence threshold for mass-spring system", ".0001");
      addParm("Solver Process", "Name of the process for the Meinhardt Solver", "Model/CCF/02 Mass Spring Solver");
      addParm("Tissue Process", "Name of process for Cell Tissue", "Model/CCF/10 Cell Tissue");
      addParm("Growth Process", "Name of the process for Growth", "Model/CCF/04 Mass Spring Growth");
      addParm("Divide Process", "Name of the process for Cell Division", "Model/CCF/05 Divide Cells");
      addParm("Split Edges Process", "Name of the process to split edges", "Model/CCF/06 Split Edges");
      addParm("Auxin Gradient Model", "Name of process for auxin model derivatives", "Model/CCF/Auxin Gradient Model");
      
      // Run control parameters
      addParm("Run Name", "Name of the run", "New Run");
      addParm("Run Max Steps", "(Int) Maximum number of steps after which the run stops. Leave blank to disable this feature", "150");
      addParm("Output directory", "Write cell data and screenshot to this directory", "/home/ahr75/Documents/Shuyao/oofs_output/" + QDateTime::currentDateTime().toString("yyyyMMdd"));
      addParm("Output frequency", "(Int) Write output each this number of steps. Leave blank to disable output", "10");
      addParm("Autorun", "Take auxin parameters from the run name?", "False", QStringList() << "True" << "False");
      addParm("Unchanging Noise", "Auxin production noise is unchanged through time?", "False", QStringList() << "True" << "False");
    }
    bool initialize(QWidget *parent);
    bool step();
    bool rewind(QWidget *parent);
    
    // Custom functions
    bool checkStop();
    void displayAttrs();
    void outputToFile();
    
    CellTissue &tissue(); // Function to retrieve the Cell Tissue

  private:
    Mesh *mesh = 0;

    CellDiskSolver *solverProcess = 0;
    CellDiskTissue *tissueProcess = 0;
    CellDiskGrowth *growthProcess = 0;
    CellDiskDivide *divideProcess = 0;
    CellDiskSplitEdges *splitEdgesProcess = 0;
    
    CCIndexDataAttr *indexAttr = 0;
    MSCellDataAttr *cellAttr = 0;
    MSEdgeDataAttr *msEdgeAttr = 0;
    AuxinGradient *auxinProcess = 0;

    // Run control parameters
    QString runName = "New Run";
    QString runMaxSteps = "";
    QString outDir = "./";
    QString outFreq = "";
    int stepCount = 0;

  };

  
  
  

  class CellDiskSolver :  public Process, public Solver<Point3d>
  {
  public:
    CellDiskSolver(const Process &process) : Process(process) 
    {
      setName("Model/CCF/02 Mass Spring Solver");
      addParm("Mass Spring Process", "Name of derivs process for mass-spring mechanics", "Model/CCF/03 Mass Spring Derivs");
      addParm("Tissue Process", "Name of process for Cell Tissue", "Model/CCF/10 Cell Tissue");
    }
    bool initialize(QWidget *parent);
  };

  class MassSpringDerivs : public Process, public SolverDerivs<Point3d>
  {
  public:
    MassSpringDerivs(const Process &process): Process(process) 
    {
      setName("Model/CCF/03 Mass Spring Derivs");
      insertParm("Spring Constant", "Stiffness of springs", "1.0", 1);
      insertParm("Pressure", "Pressure", "1.0", 2);
    }

    // Reimplemented solver methods
    void initDerivs(SolverT &solver, VertexAttr &vAttr, EdgeAttr &eAttr);
    void calcDerivatives(const SolverT &solver, CCIndex v, VectorT &values);

    void setValues(const SolverT &solver, CCIndex v, const VectorT &values)
    {
      (*indexAttr)[v].pos = values;
    }
  
    void getValues(const SolverT &solver, CCIndex v, VectorT &values)
    {
      values = (*indexAttr)[v].pos;
    }

    CCStructure const *cs = 0;
    CCIndexDataAttr *indexAttr = 0;
    MSEdgeDataAttr *msEdgeAttr = 0;

    double k = 1.0;
    double pressure = 0.5;
  };

  class CellDiskGrowth : public Process
  {
  public:
    CellDiskGrowth(const Process &process): Process(process) 
    {
      setName("Model/CCF/04 Mass Spring Growth");
      addParm("Growth Rate", "The rate of growth", "1.0");
      addParm("Dt", "Time step for growth", "0.1");

      addParm("Tissue Process", "Name of the process for the Cell Tissue", "Model/CCF/10 Cell Tissue");
    }

    // Initialize to grab subdivider
    bool initialize(QWidget *parent);

    // Run a step of cell division
    bool step();

  private:
    Mesh *mesh = 0;
    CellDiskTissue *tissueProcess = 0;
  };

  class CellDiskDivide : public CellTissueCell2dDivide
  {
  public:
    CellDiskDivide(const Process &process) : CellTissueCell2dDivide(process)
    {
      setName("Model/CCF/05 Divide Cells");
      addParm("Solver Process", "Name of the process for the Meinhardt solver", "Model/CCF/02 Mass Spring Solver");
      addParm("Tissue Process", "Name of the process for the Cell Tissue", "Model/CCF/10 Cell Tissue");
    }

    // Initialize to grab subdivider
    bool initialize(QWidget *parent);

    // Run a step of cell division
    bool step();

    CellDiskSubdivide *subdivider() { return &subdiv; }

  private:
    CellDiskSubdivide subdiv;

    CellDiskSolver *solverProcess = 0;
    CellDiskTissue *tissueProcess = 0;
  };

  class CellDiskSplitEdges : public SplitEdges
  {
  public:
    CellDiskSplitEdges(const Process &process) : SplitEdges(process) 
    {
      setName("Model/CCF/06 Split Edges");
    }
  };

  class CellDiskTissue : public CellTissueProcess
  {
  public:
    CellDiskTissue(const Process &process) : CellTissueProcess(process) 
    {
      setName("Model/CCF/10 Cell Tissue");
    }
  };
  
  class CellDiskRender : public RenderPolarity
  {
  public:
    CellDiskRender(const Process &process) : RenderPolarity(process)
    {
      setName("Model/CCF/20 Render");
    }
    bool defaultDrawChoices(Mesh &mesh, const QString &ccName);
    Colorb setFacePolarityColor(CCIndex f, VizAttribute<CCIndex> &signalAttr);

    MSCellDataAttr *cellAttr = 0;
  };


  class getConc : public Process
  {
  public:
    getConc(const Process &process) : Process(process) 
    {
      setName("Model/CCF/Get Concentration");
      setDesc("Get the concentration of auxin and CUC of selected cells.");
    }
    bool initialize(QWidget *parent);
    bool run();
  };


  class setConc : public Process
  {
  public:
    setConc(const Process &process) : Process(process) 
    {
      setName("Model/CCF/Set Concentration");
      setDesc("Set the concentration of auxin and CUC of selected cells.");
      addParm("Amount Auxin", "Auxin concentration to set; or leave blank", "");
      addParm("Amount CUC", "CUC concentration to set; or leave blank", "");
    }
    bool initialize(QWidget *parent);
    bool run();

  private:
    QString AmountAux;
    QString AmountCuc;
  };

}
#endif 
```
