## Supplementary Dataset 2 for "Tradeoff Between Speed and Robustness in Primordium Initiation Mediated by Auxin-CUC1 Interaction": Makefile

```
# Name of model/process
NAME = CellDisk
MODELPATH = "Model/CCF/01 Cell Disk"
SOLIB = usrLib$(NAME).so

# List all object files here
OBJECTS = $(NAME).o qrc_$(NAME).o

# default target
target: $(SOLIB)

# Call MorphoDynamX to get makefile
include $(shell mdx --resource)/MDXProcess.mk

# Add extra compile flags here
CXXFLAGS += -Wno-unused-local-typedefs -Wno-unused-parameter -Wno-unused-value -fopenmp

# Add extra libraries here
LIBS+=

# Add extra link flags here
LD_FLAGS+= -fopenmp

# Model dependencies
$(NAME).o: $(NAME).cpp $(NAME).hpp Makefile

$(SOLIB): $(OBJECTS)

# Run the model
run: target
	mdx --model $(MODELPATH) --addlibrary $(SOLIB) $(NAME).mdxv
```
