## Supplementary Dataset 2 for "Tradeoff Between Speed and Robustness in Primordium Initiation Mediated by Auxin-CUC1 Interaction": MorphoDynamX.py

```
######## New MorphoDynamX session v2.0 r2-1395: 2023-03-19 10:01:01
Process.Tools__System__Load_View('/home/sk3245/Documents/oofs_2023-03-18/ext/02-CellModels/ext/04-MassSpring/ext/02-MassSpringAuxin/CellDisk.mdxv')
Process.Model__CCF__01_Cell_Disk('.0001', '50', 'Model/CCF/02 Mass Spring Solver', 'Model/CCF/10 Cell Tissue', 'Model/CCF/04 Mass Spring Growth', 'Model/CCF/05 Divide Cells', 'Model/CCF/06 Split Edges')
Process.Mesh__System__Save('', '/tmp/VLsIK6Sg/CellDisk.mdxm', 'no')
```
