## Supplementary Dataset 2 for "Tradeoff Between Speed and Robustness in Primordium Initiation Mediated by Auxin-CUC1 Interaction": run_20230612.py

```
Process.Mesh__System__Open('Stack1', '/home/ahr75/Documents/Shuyao/oofs_output/Init_round5_polarity_2023-06-09.mdxm', 'no', 'no')
Process.Model__CCF__Auxin_Gradient_Model('1.0', '1000.0', '1.0', '0.2', '0.0', '0.2', '0.2', 'Markov', '1', '2', '1.0', '2', '0.1', '10.0', '4.0', '0.67', '2', '0.2', '0.6', '0.2', '0.6', '0.4', '0.0', '0.7', '1e-6', 'Yes', 'No', 'No', 'No')
Process.Model__CCF__01_Cell_Disk('10', '0.01', '.0001', '10', '10', '10', '0.1', 'Euler', 'Bi-conjugate Gradient Stabilized', '50', '1e-6', 'Yes', '.0001', 'Model/CCF/02 Mass Spring Solver', 'Model/CCF/10 Cell Tissue', 'Model/CCF/04 Mass Spring Growth', 'Model/CCF/05 Divide Cells', 'Model/CCF/06 Split Edges', 'Model/CCF/Auxin Gradient Model', 'CUC0-0_1Markov2_sim6_20230612', '300', '300', '/home/ahr75/Documents/Shuyao/oofs_output/20230612', '10', 'True')
Process.Mesh__System__Open('Stack1', '/home/ahr75/Documents/Shuyao/oofs_output/Init_round5_polarity_2023-06-09.mdxm', 'no', 'no')
Process.Model__CCF__Auxin_Gradient_Model('1.0', '1000.0', '1.0', '0.2', '0.0', '0.2', '0.2', 'Markov', '1', '2', '1.0', '2', '0.1', '10.0', '4.0', '0.67', '2', '0.2', '0.6', '0.2', '0.6', '0.4', '0.0', '0.7', '1e-6', 'Yes', 'No', 'No', 'No')
Process.Model__CCF__01_Cell_Disk('10', '0.01', '.0001', '10', '10', '10', '0.1', 'Euler', 'Bi-conjugate Gradient Stabilized', '50', '1e-6', 'Yes', '.0001', 'Model/CCF/02 Mass Spring Solver', 'Model/CCF/10 Cell Tissue', 'Model/CCF/04 Mass Spring Growth', 'Model/CCF/05 Divide Cells', 'Model/CCF/06 Split Edges', 'Model/CCF/Auxin Gradient Model', 'CUC0-0_1Markov2_sim7_20230612', '300', '300', '/home/ahr75/Documents/Shuyao/oofs_output/20230612', '10', 'True')
Process.Mesh__System__Open('Stack1', '/home/ahr75/Documents/Shuyao/oofs_output/Init_round5_polarity_2023-06-09.mdxm', 'no', 'no')
Process.Model__CCF__Auxin_Gradient_Model('1.0', '1000.0', '1.0', '0.2', '0.0', '0.2', '0.2', 'Markov', '1', '2', '1.0', '2', '0.1', '10.0', '4.0', '0.67', '2', '0.2', '0.6', '0.2', '0.6', '0.4', '0.0', '0.7', '1e-6', 'Yes', 'No', 'No', 'No')
Process.Model__CCF__01_Cell_Disk('10', '0.01', '.0001', '10', '10', '10', '0.1', 'Euler', 'Bi-conjugate Gradient Stabilized', '50', '1e-6', 'Yes', '.0001', 'Model/CCF/02 Mass Spring Solver', 'Model/CCF/10 Cell Tissue', 'Model/CCF/04 Mass Spring Growth', 'Model/CCF/05 Divide Cells', 'Model/CCF/06 Split Edges', 'Model/CCF/Auxin Gradient Model', 'CUC0-0_1Markov2_sim8_20230612', '300', '300', '/home/ahr75/Documents/Shuyao/oofs_output/20230612', '10', 'True')
Process.Mesh__System__Open('Stack1', '/home/ahr75/Documents/Shuyao/oofs_output/Init_round5_polarity_2023-06-09.mdxm', 'no', 'no')
Process.Model__CCF__Auxin_Gradient_Model('1.0', '1000.0', '1.0', '0.2', '0.0', '0.2', '0.2', 'Markov', '1', '2', '1.0', '2', '0.1', '10.0', '4.0', '0.67', '2', '0.2', '0.6', '0.2', '0.6', '0.4', '0.0', '0.7', '1e-6', 'Yes', 'No', 'No', 'No')
Process.Model__CCF__01_Cell_Disk('10', '0.01', '.0001', '10', '10', '10', '0.1', 'Euler', 'Bi-conjugate Gradient Stabilized', '50', '1e-6', 'Yes', '.0001', 'Model/CCF/02 Mass Spring Solver', 'Model/CCF/10 Cell Tissue', 'Model/CCF/04 Mass Spring Growth', 'Model/CCF/05 Divide Cells', 'Model/CCF/06 Split Edges', 'Model/CCF/Auxin Gradient Model', 'CUC0-0_1Markov2_sim9_20230612', '300', '300', '/home/ahr75/Documents/Shuyao/oofs_output/20230612', '10', 'True')
Process.Mesh__System__Open('Stack1', '/home/ahr75/Documents/Shuyao/oofs_output/Init_round5_polarity_2023-06-09.mdxm', 'no', 'no')
Process.Model__CCF__Auxin_Gradient_Model('1.0', '1000.0', '1.0', '0.2', '0.0', '0.2', '0.2', 'Markov', '5', '10', '1.0', '2', '0.1', '10.0', '4.0', '0.67', '2', '0.2', '0.6', '0.2', '0.6', '0.4', '0.0', '0.7', '1e-6', 'Yes', 'No', 'No', 'No')
Process.Model__CCF__01_Cell_Disk('10', '0.01', '.0001', '10', '10', '10', '0.1', 'Euler', 'Bi-conjugate Gradient Stabilized', '50', '1e-6', 'Yes', '.0001', 'Model/CCF/02 Mass Spring Solver', 'Model/CCF/10 Cell Tissue', 'Model/CCF/04 Mass Spring Growth', 'Model/CCF/05 Divide Cells', 'Model/CCF/06 Split Edges', 'Model/CCF/Auxin Gradient Model', 'CUC0-0_5Markov10_sim6_20230612', '300', '300', '/home/ahr75/Documents/Shuyao/oofs_output/20230612', '10', 'True')
Process.Mesh__System__Open('Stack1', '/home/ahr75/Documents/Shuyao/oofs_output/Init_round5_polarity_2023-06-09.mdxm', 'no', 'no')
Process.Model__CCF__Auxin_Gradient_Model('1.0', '1000.0', '1.0', '0.2', '0.0', '0.2', '0.2', 'Markov', '5', '10', '1.0', '2', '0.1', '10.0', '4.0', '0.67', '2', '0.2', '0.6', '0.2', '0.6', '0.4', '0.0', '0.7', '1e-6', 'Yes', 'No', 'No', 'No')
Process.Model__CCF__01_Cell_Disk('10', '0.01', '.0001', '10', '10', '10', '0.1', 'Euler', 'Bi-conjugate Gradient Stabilized', '50', '1e-6', 'Yes', '.0001', 'Model/CCF/02 Mass Spring Solver', 'Model/CCF/10 Cell Tissue', 'Model/CCF/04 Mass Spring Growth', 'Model/CCF/05 Divide Cells', 'Model/CCF/06 Split Edges', 'Model/CCF/Auxin Gradient Model', 'CUC0-0_5Markov10_sim7_20230612', '300', '300', '/home/ahr75/Documents/Shuyao/oofs_output/20230612', '10', 'True')
Process.Mesh__System__Open('Stack1', '/home/ahr75/Documents/Shuyao/oofs_output/Init_round5_polarity_2023-06-09.mdxm', 'no', 'no')
Process.Model__CCF__Auxin_Gradient_Model('1.0', '1000.0', '1.0', '0.2', '0.0', '0.2', '0.2', 'Markov', '5', '10', '1.0', '2', '0.1', '10.0', '4.0', '0.67', '2', '0.2', '0.6', '0.2', '0.6', '0.4', '0.0', '0.7', '1e-6', 'Yes', 'No', 'No', 'No')
Process.Model__CCF__01_Cell_Disk('10', '0.01', '.0001', '10', '10', '10', '0.1', 'Euler', 'Bi-conjugate Gradient Stabilized', '50', '1e-6', 'Yes', '.0001', 'Model/CCF/02 Mass Spring Solver', 'Model/CCF/10 Cell Tissue', 'Model/CCF/04 Mass Spring Growth', 'Model/CCF/05 Divide Cells', 'Model/CCF/06 Split Edges', 'Model/CCF/Auxin Gradient Model', 'CUC0-0_5Markov10_sim8_20230612', '300', '300', '/home/ahr75/Documents/Shuyao/oofs_output/20230612', '10', 'True')
Process.Mesh__System__Open('Stack1', '/home/ahr75/Documents/Shuyao/oofs_output/Init_round5_polarity_2023-06-09.mdxm', 'no', 'no')
Process.Model__CCF__Auxin_Gradient_Model('1.0', '1000.0', '1.0', '0.2', '0.0', '0.2', '0.2', 'Markov', '5', '10', '1.0', '2', '0.1', '10.0', '4.0', '0.67', '2', '0.2', '0.6', '0.2', '0.6', '0.4', '0.0', '0.7', '1e-6', 'Yes', 'No', 'No', 'No')
Process.Model__CCF__01_Cell_Disk('10', '0.01', '.0001', '10', '10', '10', '0.1', 'Euler', 'Bi-conjugate Gradient Stabilized', '50', '1e-6', 'Yes', '.0001', 'Model/CCF/02 Mass Spring Solver', 'Model/CCF/10 Cell Tissue', 'Model/CCF/04 Mass Spring Growth', 'Model/CCF/05 Divide Cells', 'Model/CCF/06 Split Edges', 'Model/CCF/Auxin Gradient Model', 'CUC0-0_5Markov10_sim9_20230612', '300', '300', '/home/ahr75/Documents/Shuyao/oofs_output/20230612', '10', 'True')
Process.Mesh__System__Open('Stack1', '/home/ahr75/Documents/Shuyao/oofs_output/Init_round5_polarity_2023-06-09.mdxm', 'no', 'no')
Process.Model__CCF__Auxin_Gradient_Model('1.0', '1000.0', '1.0', '0.2', '0.0', '0.2', '0.2', 'Gaussian', '1', '2', '1.0', '2', '0.1', '10.0', '4.0', '0.67', '2', '0.2', '0.6', '0.2', '0.6', '0.4', '0.0', '0.7', '1e-6', 'Yes', 'No', 'No', 'No')
Process.Model__CCF__01_Cell_Disk('10', '0.01', '.0001', '10', '10', '10', '0.1', 'Euler', 'Bi-conjugate Gradient Stabilized', '50', '1e-6', 'Yes', '.0001', 'Model/CCF/02 Mass Spring Solver', 'Model/CCF/10 Cell Tissue', 'Model/CCF/04 Mass Spring Growth', 'Model/CCF/05 Divide Cells', 'Model/CCF/06 Split Edges', 'Model/CCF/Auxin Gradient Model', 'CUC0-0_1-0Gaussian2_sim6_20230612', '300', '300', '/home/ahr75/Documents/Shuyao/oofs_output/20230612', '10', 'True')
Process.Mesh__System__Open('Stack1', '/home/ahr75/Documents/Shuyao/oofs_output/Init_round5_polarity_2023-06-09.mdxm', 'no', 'no')
Process.Model__CCF__Auxin_Gradient_Model('1.0', '1000.0', '1.0', '0.2', '0.0', '0.2', '0.2', 'Gaussian', '1', '2', '1.0', '2', '0.1', '10.0', '4.0', '0.67', '2', '0.2', '0.6', '0.2', '0.6', '0.4', '0.0', '0.7', '1e-6', 'Yes', 'No', 'No', 'No')
Process.Model__CCF__01_Cell_Disk('10', '0.01', '.0001', '10', '10', '10', '0.1', 'Euler', 'Bi-conjugate Gradient Stabilized', '50', '1e-6', 'Yes', '.0001', 'Model/CCF/02 Mass Spring Solver', 'Model/CCF/10 Cell Tissue', 'Model/CCF/04 Mass Spring Growth', 'Model/CCF/05 Divide Cells', 'Model/CCF/06 Split Edges', 'Model/CCF/Auxin Gradient Model', 'CUC0-0_1-0Gaussian2_sim7_20230612', '300', '300', '/home/ahr75/Documents/Shuyao/oofs_output/20230612', '10', 'True')
Process.Mesh__System__Open('Stack1', '/home/ahr75/Documents/Shuyao/oofs_output/Init_round5_polarity_2023-06-09.mdxm', 'no', 'no')
Process.Model__CCF__Auxin_Gradient_Model('1.0', '1000.0', '1.0', '0.2', '0.0', '0.2', '0.2', 'Gaussian', '1', '2', '1.0', '2', '0.1', '10.0', '4.0', '0.67', '2', '0.2', '0.6', '0.2', '0.6', '0.4', '0.0', '0.7', '1e-6', 'Yes', 'No', 'No', 'No')
Process.Model__CCF__01_Cell_Disk('10', '0.01', '.0001', '10', '10', '10', '0.1', 'Euler', 'Bi-conjugate Gradient Stabilized', '50', '1e-6', 'Yes', '.0001', 'Model/CCF/02 Mass Spring Solver', 'Model/CCF/10 Cell Tissue', 'Model/CCF/04 Mass Spring Growth', 'Model/CCF/05 Divide Cells', 'Model/CCF/06 Split Edges', 'Model/CCF/Auxin Gradient Model', 'CUC0-0_1-0Gaussian2_sim8_20230612', '300', '300', '/home/ahr75/Documents/Shuyao/oofs_output/20230612', '10', 'True')
Process.Mesh__System__Open('Stack1', '/home/ahr75/Documents/Shuyao/oofs_output/Init_round5_polarity_2023-06-09.mdxm', 'no', 'no')
Process.Model__CCF__Auxin_Gradient_Model('1.0', '1000.0', '1.0', '0.2', '0.0', '0.2', '0.2', 'Gaussian', '1', '2', '1.0', '2', '0.1', '10.0', '4.0', '0.67', '2', '0.2', '0.6', '0.2', '0.6', '0.4', '0.0', '0.7', '1e-6', 'Yes', 'No', 'No', 'No')
Process.Model__CCF__01_Cell_Disk('10', '0.01', '.0001', '10', '10', '10', '0.1', 'Euler', 'Bi-conjugate Gradient Stabilized', '50', '1e-6', 'Yes', '.0001', 'Model/CCF/02 Mass Spring Solver', 'Model/CCF/10 Cell Tissue', 'Model/CCF/04 Mass Spring Growth', 'Model/CCF/05 Divide Cells', 'Model/CCF/06 Split Edges', 'Model/CCF/Auxin Gradient Model', 'CUC0-0_1-0Gaussian2_sim9_20230612', '300', '300', '/home/ahr75/Documents/Shuyao/oofs_output/20230612', '10', 'True')
Process.Mesh__System__Open('Stack1', '/home/ahr75/Documents/Shuyao/oofs_output/Init_round5_polarity_2023-06-09.mdxm', 'no', 'no')
Process.Model__CCF__Auxin_Gradient_Model('1.0', '1000.0', '1.0', '0.2', '0.0', '0.2', '0.2', 'Gaussian', '1', '2', '1.0', '100', '0.1', '10.0', '4.0', '0.67', '2', '0.2', '0.6', '0.2', '0.6', '0.4', '0.0', '0.7', '1e-6', 'Yes', 'No', 'No', 'No')
Process.Model__CCF__01_Cell_Disk('10', '0.01', '.0001', '10', '10', '10', '0.1', 'Euler', 'Bi-conjugate Gradient Stabilized', '50', '1e-6', 'Yes', '.0001', 'Model/CCF/02 Mass Spring Solver', 'Model/CCF/10 Cell Tissue', 'Model/CCF/04 Mass Spring Growth', 'Model/CCF/05 Divide Cells', 'Model/CCF/06 Split Edges', 'Model/CCF/Auxin Gradient Model', 'CUC0-0_1-0Gaussian100_sim6_20230612', '300', '300', '/home/ahr75/Documents/Shuyao/oofs_output/20230612', '10', 'True')
Process.Mesh__System__Open('Stack1', '/home/ahr75/Documents/Shuyao/oofs_output/Init_round5_polarity_2023-06-09.mdxm', 'no', 'no')
Process.Model__CCF__Auxin_Gradient_Model('1.0', '1000.0', '1.0', '0.2', '0.0', '0.2', '0.2', 'Gaussian', '1', '2', '1.0', '100', '0.1', '10.0', '4.0', '0.67', '2', '0.2', '0.6', '0.2', '0.6', '0.4', '0.0', '0.7', '1e-6', 'Yes', 'No', 'No', 'No')
Process.Model__CCF__01_Cell_Disk('10', '0.01', '.0001', '10', '10', '10', '0.1', 'Euler', 'Bi-conjugate Gradient Stabilized', '50', '1e-6', 'Yes', '.0001', 'Model/CCF/02 Mass Spring Solver', 'Model/CCF/10 Cell Tissue', 'Model/CCF/04 Mass Spring Growth', 'Model/CCF/05 Divide Cells', 'Model/CCF/06 Split Edges', 'Model/CCF/Auxin Gradient Model', 'CUC0-0_1-0Gaussian100_sim7_20230612', '300', '300', '/home/ahr75/Documents/Shuyao/oofs_output/20230612', '10', 'True')
Process.Mesh__System__Open('Stack1', '/home/ahr75/Documents/Shuyao/oofs_output/Init_round5_polarity_2023-06-09.mdxm', 'no', 'no')
Process.Model__CCF__Auxin_Gradient_Model('1.0', '1000.0', '1.0', '0.2', '0.0', '0.2', '0.2', 'Gaussian', '1', '2', '1.0', '100', '0.1', '10.0', '4.0', '0.67', '2', '0.2', '0.6', '0.2', '0.6', '0.4', '0.0', '0.7', '1e-6', 'Yes', 'No', 'No', 'No')
Process.Model__CCF__01_Cell_Disk('10', '0.01', '.0001', '10', '10', '10', '0.1', 'Euler', 'Bi-conjugate Gradient Stabilized', '50', '1e-6', 'Yes', '.0001', 'Model/CCF/02 Mass Spring Solver', 'Model/CCF/10 Cell Tissue', 'Model/CCF/04 Mass Spring Growth', 'Model/CCF/05 Divide Cells', 'Model/CCF/06 Split Edges', 'Model/CCF/Auxin Gradient Model', 'CUC0-0_1-0Gaussian100_sim8_20230612', '300', '300', '/home/ahr75/Documents/Shuyao/oofs_output/20230612', '10', 'True')
Process.Mesh__System__Open('Stack1', '/home/ahr75/Documents/Shuyao/oofs_output/Init_round5_polarity_2023-06-09.mdxm', 'no', 'no')
Process.Model__CCF__Auxin_Gradient_Model('1.0', '1000.0', '1.0', '0.2', '0.0', '0.2', '0.2', 'Gaussian', '1', '2', '1.0', '100', '0.1', '10.0', '4.0', '0.67', '2', '0.2', '0.6', '0.2', '0.6', '0.4', '0.0', '0.7', '1e-6', 'Yes', 'No', 'No', 'No')
Process.Model__CCF__01_Cell_Disk('10', '0.01', '.0001', '10', '10', '10', '0.1', 'Euler', 'Bi-conjugate Gradient Stabilized', '50', '1e-6', 'Yes', '.0001', 'Model/CCF/02 Mass Spring Solver', 'Model/CCF/10 Cell Tissue', 'Model/CCF/04 Mass Spring Growth', 'Model/CCF/05 Divide Cells', 'Model/CCF/06 Split Edges', 'Model/CCF/Auxin Gradient Model', 'CUC0-0_1-0Gaussian100_sim9_20230612', '300', '300', '/home/ahr75/Documents/Shuyao/oofs_output/20230612', '10', 'True')
Process.Mesh__System__Open('Stack1', '/home/ahr75/Documents/Shuyao/oofs_output/Init_round5_polarity_2023-06-09.mdxm', 'no', 'no')
Process.Model__CCF__Auxin_Gradient_Model('1.0', '1000.0', '1.0', '0.2', '1.0', '0.2', '0.2', 'Markov', '1', '2', '1.0', '2', '0.1', '10.0', '4.0', '0.67', '2', '0.2', '0.6', '0.2', '0.6', '0.4', '0.0', '0.7', '1e-6', 'Yes', 'No', 'No', 'No')
Process.Model__CCF__01_Cell_Disk('10', '0.01', '.0001', '10', '10', '10', '0.1', 'Euler', 'Bi-conjugate Gradient Stabilized', '50', '1e-6', 'Yes', '.0001', 'Model/CCF/02 Mass Spring Solver', 'Model/CCF/10 Cell Tissue', 'Model/CCF/04 Mass Spring Growth', 'Model/CCF/05 Divide Cells', 'Model/CCF/06 Split Edges', 'Model/CCF/Auxin Gradient Model', 'CUC1-0_1Markov2_sim6_20230612', '300', '300', '/home/ahr75/Documents/Shuyao/oofs_output/20230612', '10', 'True')
Process.Mesh__System__Open('Stack1', '/home/ahr75/Documents/Shuyao/oofs_output/Init_round5_polarity_2023-06-09.mdxm', 'no', 'no')
Process.Model__CCF__Auxin_Gradient_Model('1.0', '1000.0', '1.0', '0.2', '1.0', '0.2', '0.2', 'Markov', '1', '2', '1.0', '2', '0.1', '10.0', '4.0', '0.67', '2', '0.2', '0.6', '0.2', '0.6', '0.4', '0.0', '0.7', '1e-6', 'Yes', 'No', 'No', 'No')
Process.Model__CCF__01_Cell_Disk('10', '0.01', '.0001', '10', '10', '10', '0.1', 'Euler', 'Bi-conjugate Gradient Stabilized', '50', '1e-6', 'Yes', '.0001', 'Model/CCF/02 Mass Spring Solver', 'Model/CCF/10 Cell Tissue', 'Model/CCF/04 Mass Spring Growth', 'Model/CCF/05 Divide Cells', 'Model/CCF/06 Split Edges', 'Model/CCF/Auxin Gradient Model', 'CUC1-0_1Markov2_sim7_20230612', '300', '300', '/home/ahr75/Documents/Shuyao/oofs_output/20230612', '10', 'True')
Process.Mesh__System__Open('Stack1', '/home/ahr75/Documents/Shuyao/oofs_output/Init_round5_polarity_2023-06-09.mdxm', 'no', 'no')
Process.Model__CCF__Auxin_Gradient_Model('1.0', '1000.0', '1.0', '0.2', '1.0', '0.2', '0.2', 'Markov', '1', '2', '1.0', '2', '0.1', '10.0', '4.0', '0.67', '2', '0.2', '0.6', '0.2', '0.6', '0.4', '0.0', '0.7', '1e-6', 'Yes', 'No', 'No', 'No')
Process.Model__CCF__01_Cell_Disk('10', '0.01', '.0001', '10', '10', '10', '0.1', 'Euler', 'Bi-conjugate Gradient Stabilized', '50', '1e-6', 'Yes', '.0001', 'Model/CCF/02 Mass Spring Solver', 'Model/CCF/10 Cell Tissue', 'Model/CCF/04 Mass Spring Growth', 'Model/CCF/05 Divide Cells', 'Model/CCF/06 Split Edges', 'Model/CCF/Auxin Gradient Model', 'CUC1-0_1Markov2_sim8_20230612', '300', '300', '/home/ahr75/Documents/Shuyao/oofs_output/20230612', '10', 'True')
Process.Mesh__System__Open('Stack1', '/home/ahr75/Documents/Shuyao/oofs_output/Init_round5_polarity_2023-06-09.mdxm', 'no', 'no')
Process.Model__CCF__Auxin_Gradient_Model('1.0', '1000.0', '1.0', '0.2', '1.0', '0.2', '0.2', 'Markov', '1', '2', '1.0', '2', '0.1', '10.0', '4.0', '0.67', '2', '0.2', '0.6', '0.2', '0.6', '0.4', '0.0', '0.7', '1e-6', 'Yes', 'No', 'No', 'No')
Process.Model__CCF__01_Cell_Disk('10', '0.01', '.0001', '10', '10', '10', '0.1', 'Euler', 'Bi-conjugate Gradient Stabilized', '50', '1e-6', 'Yes', '.0001', 'Model/CCF/02 Mass Spring Solver', 'Model/CCF/10 Cell Tissue', 'Model/CCF/04 Mass Spring Growth', 'Model/CCF/05 Divide Cells', 'Model/CCF/06 Split Edges', 'Model/CCF/Auxin Gradient Model', 'CUC1-0_1Markov2_sim9_20230612', '300', '300', '/home/ahr75/Documents/Shuyao/oofs_output/20230612', '10', 'True')
Process.Mesh__System__Open('Stack1', '/home/ahr75/Documents/Shuyao/oofs_output/Init_round5_polarity_2023-06-09.mdxm', 'no', 'no')
Process.Model__CCF__Auxin_Gradient_Model('1.0', '1000.0', '1.0', '0.2', '1.0', '0.2', '0.2', 'Markov', '5', '10', '1.0', '2', '0.1', '10.0', '4.0', '0.67', '2', '0.2', '0.6', '0.2', '0.6', '0.4', '0.0', '0.7', '1e-6', 'Yes', 'No', 'No', 'No')
Process.Model__CCF__01_Cell_Disk('10', '0.01', '.0001', '10', '10', '10', '0.1', 'Euler', 'Bi-conjugate Gradient Stabilized', '50', '1e-6', 'Yes', '.0001', 'Model/CCF/02 Mass Spring Solver', 'Model/CCF/10 Cell Tissue', 'Model/CCF/04 Mass Spring Growth', 'Model/CCF/05 Divide Cells', 'Model/CCF/06 Split Edges', 'Model/CCF/Auxin Gradient Model', 'CUC1-0_5Markov10_sim6_20230612', '300', '300', '/home/ahr75/Documents/Shuyao/oofs_output/20230612', '10', 'True')
Process.Mesh__System__Open('Stack1', '/home/ahr75/Documents/Shuyao/oofs_output/Init_round5_polarity_2023-06-09.mdxm', 'no', 'no')
Process.Model__CCF__Auxin_Gradient_Model('1.0', '1000.0', '1.0', '0.2', '1.0', '0.2', '0.2', 'Markov', '5', '10', '1.0', '2', '0.1', '10.0', '4.0', '0.67', '2', '0.2', '0.6', '0.2', '0.6', '0.4', '0.0', '0.7', '1e-6', 'Yes', 'No', 'No', 'No')
Process.Model__CCF__01_Cell_Disk('10', '0.01', '.0001', '10', '10', '10', '0.1', 'Euler', 'Bi-conjugate Gradient Stabilized', '50', '1e-6', 'Yes', '.0001', 'Model/CCF/02 Mass Spring Solver', 'Model/CCF/10 Cell Tissue', 'Model/CCF/04 Mass Spring Growth', 'Model/CCF/05 Divide Cells', 'Model/CCF/06 Split Edges', 'Model/CCF/Auxin Gradient Model', 'CUC1-0_5Markov10_sim7_20230612', '300', '300', '/home/ahr75/Documents/Shuyao/oofs_output/20230612', '10', 'True')
Process.Mesh__System__Open('Stack1', '/home/ahr75/Documents/Shuyao/oofs_output/Init_round5_polarity_2023-06-09.mdxm', 'no', 'no')
Process.Model__CCF__Auxin_Gradient_Model('1.0', '1000.0', '1.0', '0.2', '1.0', '0.2', '0.2', 'Markov', '5', '10', '1.0', '2', '0.1', '10.0', '4.0', '0.67', '2', '0.2', '0.6', '0.2', '0.6', '0.4', '0.0', '0.7', '1e-6', 'Yes', 'No', 'No', 'No')
Process.Model__CCF__01_Cell_Disk('10', '0.01', '.0001', '10', '10', '10', '0.1', 'Euler', 'Bi-conjugate Gradient Stabilized', '50', '1e-6', 'Yes', '.0001', 'Model/CCF/02 Mass Spring Solver', 'Model/CCF/10 Cell Tissue', 'Model/CCF/04 Mass Spring Growth', 'Model/CCF/05 Divide Cells', 'Model/CCF/06 Split Edges', 'Model/CCF/Auxin Gradient Model', 'CUC1-0_5Markov10_sim8_20230612', '300', '300', '/home/ahr75/Documents/Shuyao/oofs_output/20230612', '10', 'True')
Process.Mesh__System__Open('Stack1', '/home/ahr75/Documents/Shuyao/oofs_output/Init_round5_polarity_2023-06-09.mdxm', 'no', 'no')
Process.Model__CCF__Auxin_Gradient_Model('1.0', '1000.0', '1.0', '0.2', '1.0', '0.2', '0.2', 'Markov', '5', '10', '1.0', '2', '0.1', '10.0', '4.0', '0.67', '2', '0.2', '0.6', '0.2', '0.6', '0.4', '0.0', '0.7', '1e-6', 'Yes', 'No', 'No', 'No')
Process.Model__CCF__01_Cell_Disk('10', '0.01', '.0001', '10', '10', '10', '0.1', 'Euler', 'Bi-conjugate Gradient Stabilized', '50', '1e-6', 'Yes', '.0001', 'Model/CCF/02 Mass Spring Solver', 'Model/CCF/10 Cell Tissue', 'Model/CCF/04 Mass Spring Growth', 'Model/CCF/05 Divide Cells', 'Model/CCF/06 Split Edges', 'Model/CCF/Auxin Gradient Model', 'CUC1-0_5Markov10_sim9_20230612', '300', '300', '/home/ahr75/Documents/Shuyao/oofs_output/20230612', '10', 'True')
Process.Mesh__System__Open('Stack1', '/home/ahr75/Documents/Shuyao/oofs_output/Init_round5_polarity_2023-06-09.mdxm', 'no', 'no')
Process.Model__CCF__Auxin_Gradient_Model('1.0', '1000.0', '1.0', '0.2', '1.0', '0.2', '0.2', 'Gaussian', '1', '2', '1.0', '2', '0.1', '10.0', '4.0', '0.67', '2', '0.2', '0.6', '0.2', '0.6', '0.4', '0.0', '0.7', '1e-6', 'Yes', 'No', 'No', 'No')
Process.Model__CCF__01_Cell_Disk('10', '0.01', '.0001', '10', '10', '10', '0.1', 'Euler', 'Bi-conjugate Gradient Stabilized', '50', '1e-6', 'Yes', '.0001', 'Model/CCF/02 Mass Spring Solver', 'Model/CCF/10 Cell Tissue', 'Model/CCF/04 Mass Spring Growth', 'Model/CCF/05 Divide Cells', 'Model/CCF/06 Split Edges', 'Model/CCF/Auxin Gradient Model', 'CUC1-0_1-0Gaussian2_sim6_20230612', '300', '300', '/home/ahr75/Documents/Shuyao/oofs_output/20230612', '10', 'True')
Process.Mesh__System__Open('Stack1', '/home/ahr75/Documents/Shuyao/oofs_output/Init_round5_polarity_2023-06-09.mdxm', 'no', 'no')
Process.Model__CCF__Auxin_Gradient_Model('1.0', '1000.0', '1.0', '0.2', '1.0', '0.2', '0.2', 'Gaussian', '1', '2', '1.0', '2', '0.1', '10.0', '4.0', '0.67', '2', '0.2', '0.6', '0.2', '0.6', '0.4', '0.0', '0.7', '1e-6', 'Yes', 'No', 'No', 'No')
Process.Model__CCF__01_Cell_Disk('10', '0.01', '.0001', '10', '10', '10', '0.1', 'Euler', 'Bi-conjugate Gradient Stabilized', '50', '1e-6', 'Yes', '.0001', 'Model/CCF/02 Mass Spring Solver', 'Model/CCF/10 Cell Tissue', 'Model/CCF/04 Mass Spring Growth', 'Model/CCF/05 Divide Cells', 'Model/CCF/06 Split Edges', 'Model/CCF/Auxin Gradient Model', 'CUC1-0_1-0Gaussian2_sim7_20230612', '300', '300', '/home/ahr75/Documents/Shuyao/oofs_output/20230612', '10', 'True')
Process.Mesh__System__Open('Stack1', '/home/ahr75/Documents/Shuyao/oofs_output/Init_round5_polarity_2023-06-09.mdxm', 'no', 'no')
Process.Model__CCF__Auxin_Gradient_Model('1.0', '1000.0', '1.0', '0.2', '1.0', '0.2', '0.2', 'Gaussian', '1', '2', '1.0', '2', '0.1', '10.0', '4.0', '0.67', '2', '0.2', '0.6', '0.2', '0.6', '0.4', '0.0', '0.7', '1e-6', 'Yes', 'No', 'No', 'No')
Process.Model__CCF__01_Cell_Disk('10', '0.01', '.0001', '10', '10', '10', '0.1', 'Euler', 'Bi-conjugate Gradient Stabilized', '50', '1e-6', 'Yes', '.0001', 'Model/CCF/02 Mass Spring Solver', 'Model/CCF/10 Cell Tissue', 'Model/CCF/04 Mass Spring Growth', 'Model/CCF/05 Divide Cells', 'Model/CCF/06 Split Edges', 'Model/CCF/Auxin Gradient Model', 'CUC1-0_1-0Gaussian2_sim8_20230612', '300', '300', '/home/ahr75/Documents/Shuyao/oofs_output/20230612', '10', 'True')
Process.Mesh__System__Open('Stack1', '/home/ahr75/Documents/Shuyao/oofs_output/Init_round5_polarity_2023-06-09.mdxm', 'no', 'no')
Process.Model__CCF__Auxin_Gradient_Model('1.0', '1000.0', '1.0', '0.2', '1.0', '0.2', '0.2', 'Gaussian', '1', '2', '1.0', '2', '0.1', '10.0', '4.0', '0.67', '2', '0.2', '0.6', '0.2', '0.6', '0.4', '0.0', '0.7', '1e-6', 'Yes', 'No', 'No', 'No')
Process.Model__CCF__01_Cell_Disk('10', '0.01', '.0001', '10', '10', '10', '0.1', 'Euler', 'Bi-conjugate Gradient Stabilized', '50', '1e-6', 'Yes', '.0001', 'Model/CCF/02 Mass Spring Solver', 'Model/CCF/10 Cell Tissue', 'Model/CCF/04 Mass Spring Growth', 'Model/CCF/05 Divide Cells', 'Model/CCF/06 Split Edges', 'Model/CCF/Auxin Gradient Model', 'CUC1-0_1-0Gaussian2_sim9_20230612', '300', '300', '/home/ahr75/Documents/Shuyao/oofs_output/20230612', '10', 'True')
Process.Mesh__System__Open('Stack1', '/home/ahr75/Documents/Shuyao/oofs_output/Init_round5_polarity_2023-06-09.mdxm', 'no', 'no')
Process.Model__CCF__Auxin_Gradient_Model('1.0', '1000.0', '1.0', '0.2', '1.0', '0.2', '0.2', 'Gaussian', '1', '2', '1.0', '100', '0.1', '10.0', '4.0', '0.67', '2', '0.2', '0.6', '0.2', '0.6', '0.4', '0.0', '0.7', '1e-6', 'Yes', 'No', 'No', 'No')
Process.Model__CCF__01_Cell_Disk('10', '0.01', '.0001', '10', '10', '10', '0.1', 'Euler', 'Bi-conjugate Gradient Stabilized', '50', '1e-6', 'Yes', '.0001', 'Model/CCF/02 Mass Spring Solver', 'Model/CCF/10 Cell Tissue', 'Model/CCF/04 Mass Spring Growth', 'Model/CCF/05 Divide Cells', 'Model/CCF/06 Split Edges', 'Model/CCF/Auxin Gradient Model', 'CUC1-0_1-0Gaussian100_sim6_20230612', '300', '300', '/home/ahr75/Documents/Shuyao/oofs_output/20230612', '10', 'True')
Process.Mesh__System__Open('Stack1', '/home/ahr75/Documents/Shuyao/oofs_output/Init_round5_polarity_2023-06-09.mdxm', 'no', 'no')
Process.Model__CCF__Auxin_Gradient_Model('1.0', '1000.0', '1.0', '0.2', '1.0', '0.2', '0.2', 'Gaussian', '1', '2', '1.0', '100', '0.1', '10.0', '4.0', '0.67', '2', '0.2', '0.6', '0.2', '0.6', '0.4', '0.0', '0.7', '1e-6', 'Yes', 'No', 'No', 'No')
Process.Model__CCF__01_Cell_Disk('10', '0.01', '.0001', '10', '10', '10', '0.1', 'Euler', 'Bi-conjugate Gradient Stabilized', '50', '1e-6', 'Yes', '.0001', 'Model/CCF/02 Mass Spring Solver', 'Model/CCF/10 Cell Tissue', 'Model/CCF/04 Mass Spring Growth', 'Model/CCF/05 Divide Cells', 'Model/CCF/06 Split Edges', 'Model/CCF/Auxin Gradient Model', 'CUC1-0_1-0Gaussian100_sim7_20230612', '300', '300', '/home/ahr75/Documents/Shuyao/oofs_output/20230612', '10', 'True')
Process.Mesh__System__Open('Stack1', '/home/ahr75/Documents/Shuyao/oofs_output/Init_round5_polarity_2023-06-09.mdxm', 'no', 'no')
Process.Model__CCF__Auxin_Gradient_Model('1.0', '1000.0', '1.0', '0.2', '1.0', '0.2', '0.2', 'Gaussian', '1', '2', '1.0', '100', '0.1', '10.0', '4.0', '0.67', '2', '0.2', '0.6', '0.2', '0.6', '0.4', '0.0', '0.7', '1e-6', 'Yes', 'No', 'No', 'No')
Process.Model__CCF__01_Cell_Disk('10', '0.01', '.0001', '10', '10', '10', '0.1', 'Euler', 'Bi-conjugate Gradient Stabilized', '50', '1e-6', 'Yes', '.0001', 'Model/CCF/02 Mass Spring Solver', 'Model/CCF/10 Cell Tissue', 'Model/CCF/04 Mass Spring Growth', 'Model/CCF/05 Divide Cells', 'Model/CCF/06 Split Edges', 'Model/CCF/Auxin Gradient Model', 'CUC1-0_1-0Gaussian100_sim8_20230612', '300', '300', '/home/ahr75/Documents/Shuyao/oofs_output/20230612', '10', 'True')
Process.Mesh__System__Open('Stack1', '/home/ahr75/Documents/Shuyao/oofs_output/Init_round5_polarity_2023-06-09.mdxm', 'no', 'no')
Process.Model__CCF__Auxin_Gradient_Model('1.0', '1000.0', '1.0', '0.2', '1.0', '0.2', '0.2', 'Gaussian', '1', '2', '1.0', '100', '0.1', '10.0', '4.0', '0.67', '2', '0.2', '0.6', '0.2', '0.6', '0.4', '0.0', '0.7', '1e-6', 'Yes', 'No', 'No', 'No')
Process.Model__CCF__01_Cell_Disk('10', '0.01', '.0001', '10', '10', '10', '0.1', 'Euler', 'Bi-conjugate Gradient Stabilized', '50', '1e-6', 'Yes', '.0001', 'Model/CCF/02 Mass Spring Solver', 'Model/CCF/10 Cell Tissue', 'Model/CCF/04 Mass Spring Growth', 'Model/CCF/05 Divide Cells', 'Model/CCF/06 Split Edges', 'Model/CCF/Auxin Gradient Model', 'CUC1-0_1-0Gaussian100_sim9_20230612', '300', '300', '/home/ahr75/Documents/Shuyao/oofs_output/20230612', '10', 'True')
Process.Mesh__System__Open('Stack1', '/home/ahr75/Documents/Shuyao/oofs_output/Init_round5_polarity_2023-06-09.mdxm', 'no', 'no')
Process.Model__CCF__Auxin_Gradient_Model('1.0', '1000.0', '1.0', '0.2', '10.0', '0.2', '0.2', 'Markov', '1', '2', '1.0', '2', '0.1', '10.0', '4.0', '0.67', '2', '0.2', '0.6', '0.2', '0.6', '0.4', '0.0', '0.7', '1e-6', 'Yes', 'No', 'No', 'No')
Process.Model__CCF__01_Cell_Disk('10', '0.01', '.0001', '10', '10', '10', '0.1', 'Euler', 'Bi-conjugate Gradient Stabilized', '50', '1e-6', 'Yes', '.0001', 'Model/CCF/02 Mass Spring Solver', 'Model/CCF/10 Cell Tissue', 'Model/CCF/04 Mass Spring Growth', 'Model/CCF/05 Divide Cells', 'Model/CCF/06 Split Edges', 'Model/CCF/Auxin Gradient Model', 'CUC10-0_1Markov2_sim6_20230612', '300', '300', '/home/ahr75/Documents/Shuyao/oofs_output/20230612', '10', 'True')
Process.Mesh__System__Open('Stack1', '/home/ahr75/Documents/Shuyao/oofs_output/Init_round5_polarity_2023-06-09.mdxm', 'no', 'no')
Process.Model__CCF__Auxin_Gradient_Model('1.0', '1000.0', '1.0', '0.2', '10.0', '0.2', '0.2', 'Markov', '1', '2', '1.0', '2', '0.1', '10.0', '4.0', '0.67', '2', '0.2', '0.6', '0.2', '0.6', '0.4', '0.0', '0.7', '1e-6', 'Yes', 'No', 'No', 'No')
Process.Model__CCF__01_Cell_Disk('10', '0.01', '.0001', '10', '10', '10', '0.1', 'Euler', 'Bi-conjugate Gradient Stabilized', '50', '1e-6', 'Yes', '.0001', 'Model/CCF/02 Mass Spring Solver', 'Model/CCF/10 Cell Tissue', 'Model/CCF/04 Mass Spring Growth', 'Model/CCF/05 Divide Cells', 'Model/CCF/06 Split Edges', 'Model/CCF/Auxin Gradient Model', 'CUC10-0_1Markov2_sim7_20230612', '300', '300', '/home/ahr75/Documents/Shuyao/oofs_output/20230612', '10', 'True')
Process.Mesh__System__Open('Stack1', '/home/ahr75/Documents/Shuyao/oofs_output/Init_round5_polarity_2023-06-09.mdxm', 'no', 'no')
Process.Model__CCF__Auxin_Gradient_Model('1.0', '1000.0', '1.0', '0.2', '10.0', '0.2', '0.2', 'Markov', '1', '2', '1.0', '2', '0.1', '10.0', '4.0', '0.67', '2', '0.2', '0.6', '0.2', '0.6', '0.4', '0.0', '0.7', '1e-6', 'Yes', 'No', 'No', 'No')
Process.Model__CCF__01_Cell_Disk('10', '0.01', '.0001', '10', '10', '10', '0.1', 'Euler', 'Bi-conjugate Gradient Stabilized', '50', '1e-6', 'Yes', '.0001', 'Model/CCF/02 Mass Spring Solver', 'Model/CCF/10 Cell Tissue', 'Model/CCF/04 Mass Spring Growth', 'Model/CCF/05 Divide Cells', 'Model/CCF/06 Split Edges', 'Model/CCF/Auxin Gradient Model', 'CUC10-0_1Markov2_sim8_20230612', '300', '300', '/home/ahr75/Documents/Shuyao/oofs_output/20230612', '10', 'True')
Process.Mesh__System__Open('Stack1', '/home/ahr75/Documents/Shuyao/oofs_output/Init_round5_polarity_2023-06-09.mdxm', 'no', 'no')
Process.Model__CCF__Auxin_Gradient_Model('1.0', '1000.0', '1.0', '0.2', '10.0', '0.2', '0.2', 'Markov', '1', '2', '1.0', '2', '0.1', '10.0', '4.0', '0.67', '2', '0.2', '0.6', '0.2', '0.6', '0.4', '0.0', '0.7', '1e-6', 'Yes', 'No', 'No', 'No')
Process.Model__CCF__01_Cell_Disk('10', '0.01', '.0001', '10', '10', '10', '0.1', 'Euler', 'Bi-conjugate Gradient Stabilized', '50', '1e-6', 'Yes', '.0001', 'Model/CCF/02 Mass Spring Solver', 'Model/CCF/10 Cell Tissue', 'Model/CCF/04 Mass Spring Growth', 'Model/CCF/05 Divide Cells', 'Model/CCF/06 Split Edges', 'Model/CCF/Auxin Gradient Model', 'CUC10-0_1Markov2_sim9_20230612', '300', '300', '/home/ahr75/Documents/Shuyao/oofs_output/20230612', '10', 'True')
Process.Mesh__System__Open('Stack1', '/home/ahr75/Documents/Shuyao/oofs_output/Init_round5_polarity_2023-06-09.mdxm', 'no', 'no')
Process.Model__CCF__Auxin_Gradient_Model('1.0', '1000.0', '1.0', '0.2', '10.0', '0.2', '0.2', 'Markov', '5', '10', '1.0', '2', '0.1', '10.0', '4.0', '0.67', '2', '0.2', '0.6', '0.2', '0.6', '0.4', '0.0', '0.7', '1e-6', 'Yes', 'No', 'No', 'No')
Process.Model__CCF__01_Cell_Disk('10', '0.01', '.0001', '10', '10', '10', '0.1', 'Euler', 'Bi-conjugate Gradient Stabilized', '50', '1e-6', 'Yes', '.0001', 'Model/CCF/02 Mass Spring Solver', 'Model/CCF/10 Cell Tissue', 'Model/CCF/04 Mass Spring Growth', 'Model/CCF/05 Divide Cells', 'Model/CCF/06 Split Edges', 'Model/CCF/Auxin Gradient Model', 'CUC10-0_5Markov10_sim6_20230612', '300', '300', '/home/ahr75/Documents/Shuyao/oofs_output/20230612', '10', 'True')
Process.Mesh__System__Open('Stack1', '/home/ahr75/Documents/Shuyao/oofs_output/Init_round5_polarity_2023-06-09.mdxm', 'no', 'no')
Process.Model__CCF__Auxin_Gradient_Model('1.0', '1000.0', '1.0', '0.2', '10.0', '0.2', '0.2', 'Markov', '5', '10', '1.0', '2', '0.1', '10.0', '4.0', '0.67', '2', '0.2', '0.6', '0.2', '0.6', '0.4', '0.0', '0.7', '1e-6', 'Yes', 'No', 'No', 'No')
Process.Model__CCF__01_Cell_Disk('10', '0.01', '.0001', '10', '10', '10', '0.1', 'Euler', 'Bi-conjugate Gradient Stabilized', '50', '1e-6', 'Yes', '.0001', 'Model/CCF/02 Mass Spring Solver', 'Model/CCF/10 Cell Tissue', 'Model/CCF/04 Mass Spring Growth', 'Model/CCF/05 Divide Cells', 'Model/CCF/06 Split Edges', 'Model/CCF/Auxin Gradient Model', 'CUC10-0_5Markov10_sim7_20230612', '300', '300', '/home/ahr75/Documents/Shuyao/oofs_output/20230612', '10', 'True')
Process.Mesh__System__Open('Stack1', '/home/ahr75/Documents/Shuyao/oofs_output/Init_round5_polarity_2023-06-09.mdxm', 'no', 'no')
Process.Model__CCF__Auxin_Gradient_Model('1.0', '1000.0', '1.0', '0.2', '10.0', '0.2', '0.2', 'Markov', '5', '10', '1.0', '2', '0.1', '10.0', '4.0', '0.67', '2', '0.2', '0.6', '0.2', '0.6', '0.4', '0.0', '0.7', '1e-6', 'Yes', 'No', 'No', 'No')
Process.Model__CCF__01_Cell_Disk('10', '0.01', '.0001', '10', '10', '10', '0.1', 'Euler', 'Bi-conjugate Gradient Stabilized', '50', '1e-6', 'Yes', '.0001', 'Model/CCF/02 Mass Spring Solver', 'Model/CCF/10 Cell Tissue', 'Model/CCF/04 Mass Spring Growth', 'Model/CCF/05 Divide Cells', 'Model/CCF/06 Split Edges', 'Model/CCF/Auxin Gradient Model', 'CUC10-0_5Markov10_sim8_20230612', '300', '300', '/home/ahr75/Documents/Shuyao/oofs_output/20230612', '10', 'True')
Process.Mesh__System__Open('Stack1', '/home/ahr75/Documents/Shuyao/oofs_output/Init_round5_polarity_2023-06-09.mdxm', 'no', 'no')
Process.Model__CCF__Auxin_Gradient_Model('1.0', '1000.0', '1.0', '0.2', '10.0', '0.2', '0.2', 'Markov', '5', '10', '1.0', '2', '0.1', '10.0', '4.0', '0.67', '2', '0.2', '0.6', '0.2', '0.6', '0.4', '0.0', '0.7', '1e-6', 'Yes', 'No', 'No', 'No')
Process.Model__CCF__01_Cell_Disk('10', '0.01', '.0001', '10', '10', '10', '0.1', 'Euler', 'Bi-conjugate Gradient Stabilized', '50', '1e-6', 'Yes', '.0001', 'Model/CCF/02 Mass Spring Solver', 'Model/CCF/10 Cell Tissue', 'Model/CCF/04 Mass Spring Growth', 'Model/CCF/05 Divide Cells', 'Model/CCF/06 Split Edges', 'Model/CCF/Auxin Gradient Model', 'CUC10-0_5Markov10_sim9_20230612', '300', '300', '/home/ahr75/Documents/Shuyao/oofs_output/20230612', '10', 'True')
Process.Mesh__System__Open('Stack1', '/home/ahr75/Documents/Shuyao/oofs_output/Init_round5_polarity_2023-06-09.mdxm', 'no', 'no')
Process.Model__CCF__Auxin_Gradient_Model('1.0', '1000.0', '1.0', '0.2', '10.0', '0.2', '0.2', 'Gaussian', '1', '2', '1.0', '2', '0.1', '10.0', '4.0', '0.67', '2', '0.2', '0.6', '0.2', '0.6', '0.4', '0.0', '0.7', '1e-6', 'Yes', 'No', 'No', 'No')
Process.Model__CCF__01_Cell_Disk('10', '0.01', '.0001', '10', '10', '10', '0.1', 'Euler', 'Bi-conjugate Gradient Stabilized', '50', '1e-6', 'Yes', '.0001', 'Model/CCF/02 Mass Spring Solver', 'Model/CCF/10 Cell Tissue', 'Model/CCF/04 Mass Spring Growth', 'Model/CCF/05 Divide Cells', 'Model/CCF/06 Split Edges', 'Model/CCF/Auxin Gradient Model', 'CUC10-0_1-0Gaussian2_sim6_20230612', '300', '300', '/home/ahr75/Documents/Shuyao/oofs_output/20230612', '10', 'True')
Process.Mesh__System__Open('Stack1', '/home/ahr75/Documents/Shuyao/oofs_output/Init_round5_polarity_2023-06-09.mdxm', 'no', 'no')
Process.Model__CCF__Auxin_Gradient_Model('1.0', '1000.0', '1.0', '0.2', '10.0', '0.2', '0.2', 'Gaussian', '1', '2', '1.0', '2', '0.1', '10.0', '4.0', '0.67', '2', '0.2', '0.6', '0.2', '0.6', '0.4', '0.0', '0.7', '1e-6', 'Yes', 'No', 'No', 'No')
Process.Model__CCF__01_Cell_Disk('10', '0.01', '.0001', '10', '10', '10', '0.1', 'Euler', 'Bi-conjugate Gradient Stabilized', '50', '1e-6', 'Yes', '.0001', 'Model/CCF/02 Mass Spring Solver', 'Model/CCF/10 Cell Tissue', 'Model/CCF/04 Mass Spring Growth', 'Model/CCF/05 Divide Cells', 'Model/CCF/06 Split Edges', 'Model/CCF/Auxin Gradient Model', 'CUC10-0_1-0Gaussian2_sim7_20230612', '300', '300', '/home/ahr75/Documents/Shuyao/oofs_output/20230612', '10', 'True')
Process.Mesh__System__Open('Stack1', '/home/ahr75/Documents/Shuyao/oofs_output/Init_round5_polarity_2023-06-09.mdxm', 'no', 'no')
Process.Model__CCF__Auxin_Gradient_Model('1.0', '1000.0', '1.0', '0.2', '10.0', '0.2', '0.2', 'Gaussian', '1', '2', '1.0', '2', '0.1', '10.0', '4.0', '0.67', '2', '0.2', '0.6', '0.2', '0.6', '0.4', '0.0', '0.7', '1e-6', 'Yes', 'No', 'No', 'No')
Process.Model__CCF__01_Cell_Disk('10', '0.01', '.0001', '10', '10', '10', '0.1', 'Euler', 'Bi-conjugate Gradient Stabilized', '50', '1e-6', 'Yes', '.0001', 'Model/CCF/02 Mass Spring Solver', 'Model/CCF/10 Cell Tissue', 'Model/CCF/04 Mass Spring Growth', 'Model/CCF/05 Divide Cells', 'Model/CCF/06 Split Edges', 'Model/CCF/Auxin Gradient Model', 'CUC10-0_1-0Gaussian2_sim8_20230612', '300', '300', '/home/ahr75/Documents/Shuyao/oofs_output/20230612', '10', 'True')
Process.Mesh__System__Open('Stack1', '/home/ahr75/Documents/Shuyao/oofs_output/Init_round5_polarity_2023-06-09.mdxm', 'no', 'no')
Process.Model__CCF__Auxin_Gradient_Model('1.0', '1000.0', '1.0', '0.2', '10.0', '0.2', '0.2', 'Gaussian', '1', '2', '1.0', '2', '0.1', '10.0', '4.0', '0.67', '2', '0.2', '0.6', '0.2', '0.6', '0.4', '0.0', '0.7', '1e-6', 'Yes', 'No', 'No', 'No')
Process.Model__CCF__01_Cell_Disk('10', '0.01', '.0001', '10', '10', '10', '0.1', 'Euler', 'Bi-conjugate Gradient Stabilized', '50', '1e-6', 'Yes', '.0001', 'Model/CCF/02 Mass Spring Solver', 'Model/CCF/10 Cell Tissue', 'Model/CCF/04 Mass Spring Growth', 'Model/CCF/05 Divide Cells', 'Model/CCF/06 Split Edges', 'Model/CCF/Auxin Gradient Model', 'CUC10-0_1-0Gaussian2_sim9_20230612', '300', '300', '/home/ahr75/Documents/Shuyao/oofs_output/20230612', '10', 'True')
Process.Mesh__System__Open('Stack1', '/home/ahr75/Documents/Shuyao/oofs_output/Init_round5_polarity_2023-06-09.mdxm', 'no', 'no')
Process.Model__CCF__Auxin_Gradient_Model('1.0', '1000.0', '1.0', '0.2', '10.0', '0.2', '0.2', 'Gaussian', '1', '2', '1.0', '100', '0.1', '10.0', '4.0', '0.67', '2', '0.2', '0.6', '0.2', '0.6', '0.4', '0.0', '0.7', '1e-6', 'Yes', 'No', 'No', 'No')
Process.Model__CCF__01_Cell_Disk('10', '0.01', '.0001', '10', '10', '10', '0.1', 'Euler', 'Bi-conjugate Gradient Stabilized', '50', '1e-6', 'Yes', '.0001', 'Model/CCF/02 Mass Spring Solver', 'Model/CCF/10 Cell Tissue', 'Model/CCF/04 Mass Spring Growth', 'Model/CCF/05 Divide Cells', 'Model/CCF/06 Split Edges', 'Model/CCF/Auxin Gradient Model', 'CUC10-0_1-0Gaussian100_sim6_20230612', '300', '300', '/home/ahr75/Documents/Shuyao/oofs_output/20230612', '10', 'True')
Process.Mesh__System__Open('Stack1', '/home/ahr75/Documents/Shuyao/oofs_output/Init_round5_polarity_2023-06-09.mdxm', 'no', 'no')
Process.Model__CCF__Auxin_Gradient_Model('1.0', '1000.0', '1.0', '0.2', '10.0', '0.2', '0.2', 'Gaussian', '1', '2', '1.0', '100', '0.1', '10.0', '4.0', '0.67', '2', '0.2', '0.6', '0.2', '0.6', '0.4', '0.0', '0.7', '1e-6', 'Yes', 'No', 'No', 'No')
Process.Model__CCF__01_Cell_Disk('10', '0.01', '.0001', '10', '10', '10', '0.1', 'Euler', 'Bi-conjugate Gradient Stabilized', '50', '1e-6', 'Yes', '.0001', 'Model/CCF/02 Mass Spring Solver', 'Model/CCF/10 Cell Tissue', 'Model/CCF/04 Mass Spring Growth', 'Model/CCF/05 Divide Cells', 'Model/CCF/06 Split Edges', 'Model/CCF/Auxin Gradient Model', 'CUC10-0_1-0Gaussian100_sim7_20230612', '300', '300', '/home/ahr75/Documents/Shuyao/oofs_output/20230612', '10', 'True')
Process.Mesh__System__Open('Stack1', '/home/ahr75/Documents/Shuyao/oofs_output/Init_round5_polarity_2023-06-09.mdxm', 'no', 'no')
Process.Model__CCF__Auxin_Gradient_Model('1.0', '1000.0', '1.0', '0.2', '10.0', '0.2', '0.2', 'Gaussian', '1', '2', '1.0', '100', '0.1', '10.0', '4.0', '0.67', '2', '0.2', '0.6', '0.2', '0.6', '0.4', '0.0', '0.7', '1e-6', 'Yes', 'No', 'No', 'No')
Process.Model__CCF__01_Cell_Disk('10', '0.01', '.0001', '10', '10', '10', '0.1', 'Euler', 'Bi-conjugate Gradient Stabilized', '50', '1e-6', 'Yes', '.0001', 'Model/CCF/02 Mass Spring Solver', 'Model/CCF/10 Cell Tissue', 'Model/CCF/04 Mass Spring Growth', 'Model/CCF/05 Divide Cells', 'Model/CCF/06 Split Edges', 'Model/CCF/Auxin Gradient Model', 'CUC10-0_1-0Gaussian100_sim8_20230612', '300', '300', '/home/ahr75/Documents/Shuyao/oofs_output/20230612', '10', 'True')
Process.Mesh__System__Open('Stack1', '/home/ahr75/Documents/Shuyao/oofs_output/Init_round5_polarity_2023-06-09.mdxm', 'no', 'no')
Process.Model__CCF__Auxin_Gradient_Model('1.0', '1000.0', '1.0', '0.2', '10.0', '0.2', '0.2', 'Gaussian', '1', '2', '1.0', '100', '0.1', '10.0', '4.0', '0.67', '2', '0.2', '0.6', '0.2', '0.6', '0.4', '0.0', '0.7', '1e-6', 'Yes', 'No', 'No', 'No')
Process.Model__CCF__01_Cell_Disk('10', '0.01', '.0001', '10', '10', '10', '0.1', 'Euler', 'Bi-conjugate Gradient Stabilized', '50', '1e-6', 'Yes', '.0001', 'Model/CCF/02 Mass Spring Solver', 'Model/CCF/10 Cell Tissue', 'Model/CCF/04 Mass Spring Growth', 'Model/CCF/05 Divide Cells', 'Model/CCF/06 Split Edges', 'Model/CCF/Auxin Gradient Model', 'CUC10-0_1-0Gaussian100_sim9_20230612', '300', '300', '/home/ahr75/Documents/Shuyao/oofs_output/20230612', '10', 'True')
```
