## Supplementary Dataset 2 for "Tradeoff Between Speed and Robustness in Primordium Initiation Mediated by Auxin-CUC1 Interaction": Modeling_analysis_2023-10-10_demo.R

```
#######################################################################################

# Analysis of modeling output downstream of parsing
# Using demo data

# Shuyao Kong
# 2023-10-10

library(ggplot2)
library(dplyr)
library(tidyr)
library(stringr)
library(egg)
library(progress)
setwd("~/Roeder_Lab/Modeling/MassSpringAuxin/2023-10-10_Demo/")

#######################################################################################


# Load the parsed data
load("RData_20231010.RData")
sums.auxMax$budTimeID <- as.character(sums.auxMax$budTimeID)
tmp <- str_split(sums.auxMax$budTimeID, pattern = "_", simplify = T)
sums.auxMax$budID <- paste(tmp[,1], tmp[,2], tmp[,3], tmp[,4], tmp[,5], tmp[,6], tmp[,7], tmp[,8], tmp[,9], sep = "_")
sums.auxMax$time <- tmp[,10]
sums.auxMax$parm.cuc <- tmp[,2]
sums.auxMax$parm.SDaux <- tmp[,3]
sums.auxMax$parm.plast <- tmp[,6]
sums.auxMax <- sums.auxMax[sums.auxMax$num.cells >= 2,] # Filter out auxin maxima that has just 1 cell


# Name the genotypes
sums.auxMax <- unite(sums.auxMax, col = "genotype", c(parm.cuc,parm.SDaux))
sums.auxMax$genotype[sums.auxMax$genotype=="1_0.1"] <- "WT"
sums.auxMax$genotype[sums.auxMax$genotype=="1_1.0"] <- "drmy1"
sums.auxMax$genotype[sums.auxMax$genotype=="0_0.1"] <- "cuc1"
sums.auxMax$genotype[sums.auxMax$genotype=="0_1.0"] <- "drmy1cuc1"
sums.auxMax$genotype <- factor(sums.auxMax$genotype, levels = c("WT", "cuc1", "drmy1", "drmy1cuc1"))
sums.auxMax <- sums.auxMax[!is.na(sums.auxMax$genotype),]


# Summary of sample size (how many buds)
tmp <- sums.auxMax[,c("budID","genotype","parm.plast")] %>% unique
table(tmp[,c("genotype","parm.plast")])


# Select the time points of interest:
sums.auxMax.t <- rbind(sums.auxMax[sums.auxMax$parm.plast=="0.4" & sums.auxMax$time=="140",],
                       sums.auxMax[sums.auxMax$parm.plast=="0.8" & sums.auxMax$time=="100",],
                       sums.auxMax[sums.auxMax$parm.plast=="1.2" & sums.auxMax$time=="90",])


# Plotting auxin maxima intensity, WT, cuc1, drmy1, drmy1cuc1 (growth rate 0.8)
tmp <- sums.auxMax.t[sums.auxMax.t$parm.plast=="0.8" & sums.auxMax.t$time=="100",]
tmp <- tmp %>% group_by(budTimeID, genotype) %>%
  mutate(aux.conc = aux.amount / num.cells) %>%
  summarize(aux.conc.mean = mean(aux.conc))
tmp.meansd <- tmp %>% group_by(genotype) %>%
  summarize(conc.mean = mean(aux.conc.mean), conc.sd = sd(aux.conc.mean)) %>%
  mutate(ymin = conc.mean-conc.sd, ymax = conc.mean+conc.sd)
g <- ggplot(tmp.meansd, aes(x=genotype, y=conc.mean, ymin=ymin, ymax=ymax, fill=genotype)) +
  geom_bar(stat = "identity", width=0.8) +
  geom_errorbar(width=0.4) +
  scale_fill_manual(values = c("#00BFC4","#00B81F","#F8766D","#BB9D00")) +
  scale_y_continuous(limits = c(0,40), expand = c(0,0)) +
  labs(title = "Auxin maxima conc")
panel <- set_panel_size(p = g, file = "Plots/auxConc_plast_0-8_t100.png",
                        width = unit(3, "cm"), height = unit(6, "cm"))
png(); print(panel); dev.off()


# Plotting the number of auxin maxima, WT, cuc1, drmy1, drmy1cuc1 (growth rate 0.8)
interpolate <- function(x){
  if(length(x)==1) return(x/2)
  return( (c(0,x[1:(length(x)-1)]) + x) / 2 )
}
sums.auxMax.t.0.8 <- sums.auxMax.t[sums.auxMax.t$parm.plast=="0.8",]
tmp <- sums.auxMax.t.0.8 %>% group_by(budID, genotype, parm.plast) %>% summarize(num = n())
tmp$num <- as.character(tmp$num)
tmp$num[tmp$num %in% c("1","2")] <- "≤2"
tmp$num[tmp$num %in% c("6","7","8","9","10")] <- "≥6"
tmp$num <- factor(tmp$num, levels = c("≤2","3","4","5","≥6"))
tmp.lab <- tmp %>%
  group_by(genotype, parm.plast) %>%
  do(data.frame(value = names(rev(table(.$num))),
                Cumsum = cumsum(rev(table(.$num)) / sum(rev(table(.$num)))))) %>%
  mutate(y = interpolate(Cumsum)) %>%
  mutate(Color = ifelse(value %in% c("≤2","3"), "black", "white")) %>%
  .[!is.na(.$y),] %>%
  .[.$y != 0 & .$y != 1,]

g <- ggplot(tmp, aes(x = genotype, fill = num)) +
  geom_bar(position = "fill") +
  geom_text(inherit.aes = F, data = tmp.lab, aes(x = genotype, y = y, label = value),
            color = tmp.lab$Color) +
  scale_fill_manual(values = colorRampPalette(c("white","blue4"))(5)[2:4]) +
  scale_y_continuous(limits = c(0,1.0), expand = c(0,0), breaks = 0:5*0.2) +
  guides(fill = "none") +
  labs(title = "Auxin maxima num\nplast 0.8, t=100",
       y = "Proportion", x = "") +
  theme(axis.text.x = element_text(angle = 45, hjust = 1, vjust = 1))
panel <- egg::set_panel_size(p = g, file = "Plots/AuxMaxNum_plast0-8.png", 
                             margin = unit(1, "cm"), width = unit(3, "cm"), height = unit(6, "cm"))
png(); print(panel); dev.off()


# Plotting the number of auxin maxima, WT and drmy1, in different growth rates
sums.auxMax.t.WT.drmy1 <- sums.auxMax.t[sums.auxMax.t$genotype%in%c("WT","drmy1"),]
tmp <- sums.auxMax.t.WT.drmy1 %>% group_by(budID, genotype, parm.plast) %>% summarize(num = n())
tmp$num <- as.character(tmp$num)
tmp$num[tmp$num %in% c("1","2")] <- "≤2"
tmp$num[tmp$num %in% c("6","7","8","9","10")] <- "≥6"
tmp$num <- factor(tmp$num, levels = c("≤2","3","4","5","≥6"))
tmp.lab <- tmp %>%
  group_by(genotype, parm.plast) %>%
  do(data.frame(value = names(rev(table(.$num))),
                Cumsum = cumsum(rev(table(.$num)) / sum(rev(table(.$num)))))) %>%
  mutate(y = interpolate(Cumsum)) %>%
  mutate(Color = ifelse(value %in% c("≤2","3"), "black", "white")) %>%
  .[!is.na(.$y),] %>%
  .[.$y != 0 & .$y != 1,]

g <- ggplot(tmp, aes(x = genotype, fill = num)) +
  facet_wrap(~parm.plast, nrow=1) +
  geom_bar(position = "fill") +
  geom_text(inherit.aes = F, data = tmp.lab, aes(x = genotype, y = y, label = value),
            color = tmp.lab$Color) +
  scale_fill_manual(values = colorRampPalette(c("white","blue4"))(5)[2:4]) +
  scale_y_continuous(limits = c(0,1.0), expand = c(0,0), breaks = 0:5*0.2) +
  guides(fill = "none") +
  labs(title = "Auxin maxima num\nacross growth rates",
       y = "Proportion", x = "") +
  theme(axis.text.x = element_text(angle = 45, hjust = 1, vjust = 1))
panel <- egg::set_panel_size(p = g, file = "Plots/AuxMaxNum_WT-drmy1_growth-rates.png", 
                             margin = unit(1, "cm"), width = unit(1.5, "cm"), height = unit(5.5, "cm"))
png(); print(panel); dev.off()
```
