## Supplementary Dataset 2 for "Tradeoff Between Speed and Robustness in Primordium Initiation Mediated by Auxin-CUC1 Interaction": Parse_output_2023-10-10_demo.R

```
#######################################################################################

# Parsing modeling output for downstream analysis
# Using demo data

# Shuyao Kong
#
# 2023-10-10

library(ggplot2)
library(dplyr)
library(tidyr)
library(stringr)
library(progress)
setwd("~/Documents/Shuyao/R-analysis/")

#######################################################################################


# Get list of files
simfiles <- list.files("../oofs_output_analyzed/20231010", pattern = "*.txt", full.names = T, recursive = T)
numfiles <- length(simfiles); numfiles

# Desires output
dats.filenames <- data.frame(batch = character(numfiles),
                             parm.cuc = character(numfiles),
                             parm.SDaux = character(numfiles),
                             parm.plast = character(numfiles),
                             budID = character(numfiles),
                             time = numeric(numfiles),
                             budtimeID = character(numfiles),
                             simfile = character(numfiles),
                             stringsAsFactors = F) # Stores metadata and all the file names 
dats.auxMax <- list() # Information of all cells within auxin maxima
sums.auxMax <- data.frame(budTimeID = character(), # Summarized information for auxin maxima
                          auxMaxID = character(),  
                          CCIndex = list(character()),
                          wc.r = numeric(),     # Weighed center
                          wc.rnorm = numeric(), # Normalized by the max radius of the cell disk
                          wc.theta = numeric(),
                          num.cells = numeric(),
                          aux.amount= numeric(),
                          asso.crit = character(),
                          note = character(),
                          stringsAsFactors = F)

# Output parameters
auxMax.thres <- 20 # Concentration of auxin in a cell has to be 20 folds higher than tissue-wide median to be considered part of an auxin maxima
rmin <- 1/3 # Only count the middle ring
rmax <- 2/3 # Only count the middle ring
cell.dist <- 8.5   # Maximum distance for two cell centers to be considered adjacent (belonging to the same auxin maxima)

# Global parameters
Glob.auxMaxID <- 0 # current maximum auxin maxima ID (so a new ID would be Glob.auxMaxID+1)

# Function to calculate distance between two polar coordinates
dist.pol <- function(r1, theta1, r2, theta2){
  return(sqrt(r1^2 + r2^2 - 2*r1*r2*cos(theta1-theta2)))
}

# Function to get auxin maxima from an input file
GetAuxMax <- function(dat.raw){
  
  # Find cells whose auxin concentration is auxMax.thres folds above the MEDIAN
  dat.raw$auxMax <- dat.raw$aux >= auxMax.thres * median(dat.raw$aux); length(which(dat.raw$auxMax))
  
  # Trivial cases
  if(length(which(dat.raw$auxMax))==0) return(NULL)
  if(length(which(dat.raw$auxMax))==1){
    dat.auxMax <- dat.raw[dat.raw$auxMax,c("CCIndex","x","y","aux")]
    dat.auxMax$auxMaxID <- 1 + Glob.auxMaxID
    Glob.auxMaxID <- 1 + Glob.auxMaxID
    dat.auxMax$auxMaxID <- as.character(dat.auxMax$auxMaxID)
    return(list(dat.auxMax, Glob.auxMaxID))
  }
  
  # Calculate pairwise cell distances of auxMax cells, and cluster into auxin maxima
  dat.auxMax <- dat.raw[dat.raw$auxMax,c("CCIndex","x","y","aux")]
  d <- dist(dat.auxMax[,2:3], method = "euclidean")
  hc <- hclust(d, method = "single") # "Friends of friends"
  hc2 <- cutree(hc, h = cell.dist)
  dat.auxMax$auxMaxID <- hc2 + Glob.auxMaxID
  Glob.auxMaxID <- max(dat.auxMax$auxMaxID)
  dat.auxMax$auxMaxID <- as.character(dat.auxMax$auxMaxID)
  
  return(list(dat.auxMax, Glob.auxMaxID))
}

# Function to plot the results of auxin maxima classification
PlotAuxMax <- function(simfile, dat.raw, dat.auxMax){
  if(is.null(dat.auxMax)){ # No auxin maxima identified
    g <- ggplot(dat.raw, aes(x=x, y=y)) +
      geom_point(size=5, color = "gray") +
      coord_equal()
  } else{
    g <- ggplot(dat.raw, aes(x=x, y=y)) +
      geom_point(size=5, color = "gray") +
      geom_point(data = dat.auxMax, aes(color=auxMaxID), size=5) +
      coord_equal()
  }
  png(str_replace(simfile, ".txt", ".png"), width = 420, height = 400)
  print(g)
  dev.off()
}

# Function to summarize cellular information into information of auxin maxima
SummarizeAuxMax <- function(budTimeID, dat.auxMax, r.max){
  
  tmp <- dat.auxMax %>% group_by(auxMaxID) %>%
    summarize(CCIndex = list(CCIndex),
              wc.x = sum(x*aux)/sum(aux),
              wc.y = sum(y*aux)/sum(aux), # Weighed center
              num.cells = n(),
              aux.amount = sum(aux),
              asso.crit = "unassociated", # Default
              note = "") %>%
    mutate(wc.r = sqrt(wc.x^2 + wc.y^2),
           wc.theta = atan2(wc.y, wc.x)) %>% # In radians between -pi and pi
    mutate(wc.rnorm = wc.r / r.max)
  tmp <- data.frame(budTimeID, tmp[,c(1,2,9,11,10,5:8)])
  if(endsWith(budTimeID, suffix = "_0")) tmp$asso.crit <- "First time point" # First time point
  
  return(tmp)
}

# Function to associate auxin maxima with the previous time point
AssociateAuxMax <- function(sum.auxMax, dat.auxMax, sum.auxMax.prev, dat.auxMax.prev){

  # First association criteria: overlapping CCIndex
  for(i in 1:nrow(sum.auxMax)){
    for(j in 1:nrow(sum.auxMax.prev)){
      if(length(intersect(sum.auxMax$CCIndex[[i]], sum.auxMax.prev$CCIndex[[j]])) != 0){

        auxMaxID.curr <- sum.auxMax$auxMaxID[i]      # ID in the current time point
        auxMaxID.prev <- sum.auxMax.prev$auxMaxID[j] # ID in the previous time point

        # Test if this is the first auxin maxima in the previous time point that it can be associated with
        if(sum.auxMax$asso.crit[i] == "unassociated"){

          # Test if this auxin maxima in the previous time point has been associated with any other auxin maxima in the current time point
          # If so, this means that the auxin maxima in the previous time point has split into two. (Currently not supporting splitting in three)
          # Name them as such: 1a,1b
          if(auxMaxID.prev %in% sum.auxMax$auxMaxID){
            ID1 <- paste0(auxMaxID.prev, "a")
            ID2 <- paste0(auxMaxID.prev, "b")
            sum.auxMax$auxMaxID[sum.auxMax$auxMaxID == auxMaxID.prev] <- ID1
            dat.auxMax$auxMaxID[dat.auxMax$auxMaxID == auxMaxID.prev] <- ID1
            sum.auxMax$auxMaxID[i] <- ID2
            dat.auxMax$auxMaxID[dat.auxMax$auxMaxID == auxMaxID.curr] <- ID2
            sum.auxMax$note[sum.auxMax$auxMaxID %in% c(ID1,ID2)] <- "Split"
          } else{
            sum.auxMax$auxMaxID[i] <- auxMaxID.prev
            dat.auxMax$auxMaxID[dat.auxMax$auxMaxID == auxMaxID.curr] <- auxMaxID.prev
          }
        }
        # If it has been associated with other auxin maxima in the previous time point, this means multiple auxin maxima has merged
        # Name this auxin maxima as such: 1+2
        else{
          ID <- paste(auxMaxID.prev, auxMaxID.curr, sep = "+")
          sum.auxMax$auxMaxID[i] <- ID
          sum.auxMax$note[i] <- "Merged"
          dat.auxMax$auxMaxID[dat.auxMax$auxMaxID == auxMaxID.curr] <- ID
        }

        # Take a note of the association method
        sum.auxMax$asso.crit[i] <- "CCIndex"
      }
    }
  }

  # Second attempt to associate: close (unweighed) position, within 1 cell distance
  for(i in 1:nrow(sum.auxMax)){
    if(sum.auxMax$asso.crit[i] == "unassociated"){
      for(j in 1:nrow(sum.auxMax.prev)){
        if(dist.pol(sum.auxMax$wc.r[i], sum.auxMax$wc.theta[i], sum.auxMax.prev$wc.r[j], sum.auxMax.prev$wc.theta[j]) <= cell.dist){

          auxMaxID.curr <- sum.auxMax$auxMaxID[i]      # ID in the current time point
          auxMaxID.prev <- sum.auxMax.prev$auxMaxID[j] # ID in the previous time point

          # Test if this is the first auxin maxima in the previous time point that it can be associated with
          if(sum.auxMax$asso.crit[i] == "unassociated"){

            # Test if this auxin maxima in the previous time point has been associated with any other auxin maxima in the current time point
            # If so, this means that the auxin maxima in the previous time point has split into two. (Currently not supporting splitting in three)
            # Name them as such: 1a,1b
            if(auxMaxID.prev %in% sum.auxMax$auxMaxID){
              ID1 <- paste0(auxMaxID.prev, "a")
              ID2 <- paste0(auxMaxID.prev, "b")
              sum.auxMax$auxMaxID[sum.auxMax$auxMaxID == auxMaxID.prev] <- ID1
              dat.auxMax$auxMaxID[dat.auxMax$auxMaxID == auxMaxID.prev] <- ID1
              sum.auxMax$auxMaxID[i] <- ID2
              dat.auxMax$auxMaxID[dat.auxMax$auxMaxID == auxMaxID.curr] <- ID2
              sum.auxMax$note[sum.auxMax$auxMaxID %in% c(ID1,ID2)] <- "Split"
            } else{
              sum.auxMax$auxMaxID[i] <- auxMaxID.prev
              dat.auxMax$auxMaxID[dat.auxMax$auxMaxID == auxMaxID.curr] <- auxMaxID.prev
            }
          }
          # If it has been associated with other auxin maxima in the previous time point, this means multiple auxin maxima has merged
          # Name this auxin maxima as such: 1+2
          else{
            ID <- paste(auxMaxID.prev, auxMaxID.curr, sep = "+")
            sum.auxMax$auxMaxID[i] <- ID
            sum.auxMax$note[i] <- "Merged"
            dat.auxMax$auxMaxID[dat.auxMax$auxMaxID == auxMaxID.curr] <- ID
          }

          # Take a note of the association method
          sum.auxMax$asso.crit[i] <- "Distance"
        }
      }
    }
  }

  return(list(sum.auxMax,dat.auxMax))
}


# Parse the file names of each file for easy query
pb <- progress_bar$new(format = "Parsing file names [:bar] :percent, eta :eta", clear = F, total = numfiles, width = 80)
for(i in 1:numfiles){
  simfile.name <- str_split(simfiles[i], "/")[[1]] %>% .[length(.)] %>% str_remove(".txt")
  tmp <- str_split(simfile.name, "[_]")[[1]]
  dats.filenames[i,"parm.cuc"] <- tmp[3]
  dats.filenames[i,"parm.SDaux"] <- tmp[4]
  dats.filenames[i,"parm.plast"] <- tmp[7]
  dats.filenames[i,"batch"] <- tmp[10]
  dats.filenames[i,"time"] <- as.numeric(tmp[11])
  dats.filenames[i,"budID"] <- paste(tmp[2:10], collapse = "_")
  dats.filenames[i,"budtimeID"] <- paste(tmp[2:11], collapse = "_") # Corresponds one-to-one to simfile
  dats.filenames[i,"simfile"] <- simfiles[i]
  pb$tick()
}


# Parse all the CCData files 
budIDs <- unique(dats.filenames$budID)
nBuds <- length(budIDs)
pb <- progress_bar$new(format = "Parsing CCData [:bar] :percent, eta :eta", clear = F, total = nBuds, width = 80)

for(budID in budIDs){
  
  # Get all relevant files (all time points of this bud)
  tmp.dats.filenames <- dats.filenames[dats.filenames$budID==budID,] %>% arrange(time)
  
  # Loop over all the time points
  for(i in 1:nrow(tmp.dats.filenames)){
    
    # Read in a file for the current time point
    simfile <- tmp.dats.filenames$simfile[i]
    budTimeID <- tmp.dats.filenames$budtimeID[i]
    time <- str_split(budTimeID, "_")[[1]] %>% .[length(.)]
    dat.raw <- read.table(simfile, header = T, sep = '\t') # Read in raw data file
    r.max <- max(sqrt(dat.raw$x^2 + dat.raw$y^2)) # Maximum radius of the disk (for normalization)
    
    # Try to get auxin maxima
    tmp <- GetAuxMax(dat.raw) 
    dat.auxMax <- tmp[[1]]
    if(!is.null(tmp)){ # If there is any auxin maxima
      Glob.auxMaxID <- tmp[[2]]
      
      # Summarize auxin maxima information of the current time point
      sum.auxMax <- SummarizeAuxMax(budTimeID, dat.auxMax, r.max) 
      
      # Associate auxin maxima with the previous time point
      budTimeID.prev <- tmp.dats.filenames$budtimeID[i-1]
      sum.auxMax.prev <- sums.auxMax[sums.auxMax$budTimeID == budTimeID.prev,]
      if(nrow(sum.auxMax.prev)>0){
        dat.auxMax.prev <- dats.auxMax[[budTimeID.prev]]
        tmp <- AssociateAuxMax(sum.auxMax, dat.auxMax, sum.auxMax.prev, dat.auxMax.prev)
        sum.auxMax <- tmp[[1]]
        dat.auxMax <- tmp[[2]]
      }
      
      # Only retain auxin maxima whose center falls in the middle ring
      sum.auxMax <- sum.auxMax[sum.auxMax$wc.rnorm >= rmin & sum.auxMax$wc.rnorm <= rmax,]
      auxMaxIDs <- sum.auxMax$auxMaxID
      dat.auxMax <- dat.auxMax[dat.auxMax$auxMaxID %in% auxMaxIDs,]

      # Write to existing data tables
      sums.auxMax <- rbind(sums.auxMax, sum.auxMax)
      dats.auxMax <- append(dats.auxMax, list(dat.auxMax))
      names(dats.auxMax)[length(dats.auxMax)] <- budTimeID
    }
    
    # Output an image showing how auxin maxima are assigned
    PlotAuxMax(simfile, dat.raw, dat.auxMax)
    
  }
  
  pb$tick()
}


# Save RData
RData.name <- "RData_20231010.RData"
save.image(RData.name)
```
