## Supplementary Dataset 2 for "Tradeoff Between Speed and Robustness in Primordium Initiation Mediated by Auxin-CUC1 Interaction": Readme.pdf

### Computational model of auxin-CUC interaction in the floral meristem

10/10/2023

#### Summary

The floral meristem is modeled using a 2D disk of cells that are growing and dividing, based on two previous models, GridAuxin and MassSpringCells. Mechanics are modeled using a mass-spring model where cell walls (springs) are pulling cell wall junctions (masses). Each cell has auxin, CUC, and PIN, that are produced, degraded, and interacting, calculated using a differential equations solver. The model is implemented in C++, and can be run inside the VLab environment[1] and the MorphoDynamX software[2]. The model can be run either one simulation at a time, or batch-run using a python code.

#### System requirements

The model requires a Linux machine with NVIDIA CUDA version 11.40. MorphoDynamX (www.MorphoDynamX.org) and VLab (http://algorithmicbotany.org/virtual_laboratory/). We suggest using R for downstream data parsing.

For the demo, we used vlab (version 5.0, build #3609) and MorphoDynamX (version 2.0, revision 2-1395, CUDA version 11.40) in an Ubuntu 20.04.6 LTS system, equipped with an Intel Core i9-10900X 3.7 GHz 10-Core Processor, G.Skill Trident Z RGB 256 GB DDR4-3600 CL18 Memory, and a NVIDIA GeForce RTX 4090 24 GB Graphics Card. For downstream data processing, we used RStudio (R version 4.3.1 (2023-06-16) -- "Beagle Scouts").

#### The model folder

The folder “MassSpringAuxin” contains the source files for the model.

- “specifications.txt” lists all the files within the folder to be included with the model
- “CellDisk.hpp” and “CellDisk.cpp” are the model source files, and can be edited using a text editor or IDEs such as VS Code.
- “CellDisk.mdxv” can be edited using a text editor, and contain metadata of the model, including which initial condition to use.
- “Init_blank_72cells_2023-09-07.mdxm” is the initial condition used in the model. This can be changed to another MDX mesh file.

**Running the model one simulation at a time**

1. Open the VLab Browser.
2. Select the oofs repository.
3. Navigate to the model, and double click on the window to open it.
4. Right click on the model icon, and select “run”. The model would now open in MDX.
5. Set the desired parameters, mainly in Process -> Model -> 01 Cell Disk and Process -> Model -> Auxin Gradient Model. If output is desired, set “Run name”, “Output directory”, and “Output frequency”.
6. Double-click Process -> Model -> 01 Cell Disk to run it. Click on the stop button to stop when desired.

**Running the model in batch mode**

1. Using a python code, such as the provided “create_run_20231010_demo.py”, create a run file, such as the provided “run_20231010.py”. The former file specifies the parameter combinations to be tested, while the latter file contains the commands to be fed into MDX.
2. Open the model in MDX following steps 1-4 in the previous section.
3. Under Process -> Tools -> Python -> Python Script, select the run file created in step 1 (“run_20231010.py”). The model would now run automatically.
4. Screenshots are automatically saved in “output directory/run name/Screenshots/*.jpg”. Cell complex data are automatically saved in “output directory/run name/CCData/*.txt”. The following data are saved in these files: Cell index (CCIndex), x and y coordinates, cell area, and auxin and CUC concentrations. Each row represents a cell.
5. In the demo, 5 simulations each of WT, *drmy1*, *cuc1*, and *drmy1 cuc1*, with growth rates 0.4, 0.8, and 1.2 were run, and the results were saved in the directory “20231010” (in Github, these are uploaded as five zip files, 20231010 *.zip). The simulations were named as follows: “*Run_CUC Prod_SD Auxin Prod_PIN low CUC_PIN high CUC_Plasticity_Unchanging Noise_sim number_date*”.

**Parsing and analysis of CCData**

1. The code “Parse_output_2023-10-10_demo.R” parses these data files, identifies auxin maxima, tracks them over time, and summarizes them in “RData_20231010.RData”. It also plots how auxin maxima are assigned and tracked. These plots are saved in “output directory/run name/CCData/*.png”.
2. The data frames in the output RData file can be analyzed as desired. For example, the provided code “Modeling_analysis_2023-10-10_demo.R” reads in the RData file and generates three plots in the “Plots” folder.
