## Supplementary Dataset 2 for "Tradeoff Between Speed and Robustness in Primordium Initiation Mediated by Auxin-CUC1 Interaction": run_20231010.py

```
Process.Mesh__System__Open('Stack1', '/home/ahr75/Documents/Shuyao/oofs_2023-03-18/ext/02-CellModels/ext/04-MassSpring/ext/19-MassSpringAuxinQuadratic/Init_blank_72cells_2023-09-07.mdxm', 'no', 'no')
Process.Model__CCF__01_Cell_Disk('10', '0.01', '.0001', '10', '10', '10', '0.1', 'Euler', 'Bi-conjugate Gradient Stabilized', '50', '1e-6', 'No', '.0001', 'Model/CCF/02 Mass Spring Solver', 'Model/CCF/10 Cell Tissue', 'Model/CCF/04 Mass Spring Growth', 'Model/CCF/05 Divide Cells', 'Model/CCF/06 Split Edges', 'Model/CCF/Auxin Gradient Model', 'Run_1_0-1_Linear_Quadratic_0-4_False_sim1_20231010', '140', '/home/ahr75/Documents/Shuyao/oofs_output/20231010', '10', 'True', 'False')
Process.Mesh__System__Open('Stack1', '/home/ahr75/Documents/Shuyao/oofs_2023-03-18/ext/02-CellModels/ext/04-MassSpring/ext/19-MassSpringAuxinQuadratic/Init_blank_72cells_2023-09-07.mdxm', 'no', 'no')
Process.Model__CCF__01_Cell_Disk('10', '0.01', '.0001', '10', '10', '10', '0.1', 'Euler', 'Bi-conjugate Gradient Stabilized', '50', '1e-6', 'No', '.0001', 'Model/CCF/02 Mass Spring Solver', 'Model/CCF/10 Cell Tissue', 'Model/CCF/04 Mass Spring Growth', 'Model/CCF/05 Divide Cells', 'Model/CCF/06 Split Edges', 'Model/CCF/Auxin Gradient Model', 'Run_1_0-1_Linear_Quadratic_0-8_False_sim1_20231010', '140', '/home/ahr75/Documents/Shuyao/oofs_output/20231010', '10', 'True', 'False')
Process.Mesh__System__Open('Stack1', '/home/ahr75/Documents/Shuyao/oofs_2023-03-18/ext/02-CellModels/ext/04-MassSpring/ext/19-MassSpringAuxinQuadratic/Init_blank_72cells_2023-09-07.mdxm', 'no', 'no')
Process.Model__CCF__01_Cell_Disk('10', '0.01', '.0001', '10', '10', '10', '0.1', 'Euler', 'Bi-conjugate Gradient Stabilized', '50', '1e-6', 'No', '.0001', 'Model/CCF/02 Mass Spring Solver', 'Model/CCF/10 Cell Tissue', 'Model/CCF/04 Mass Spring Growth', 'Model/CCF/05 Divide Cells', 'Model/CCF/06 Split Edges', 'Model/CCF/Auxin Gradient Model', 'Run_1_0-1_Linear_Quadratic_1-2_False_sim1_20231010', '140', '/home/ahr75/Documents/Shuyao/oofs_output/20231010', '10', 'True', 'False')
Process.Mesh__System__Open('Stack1', '/home/ahr75/Documents/Shuyao/oofs_2023-03-18/ext/02-CellModels/ext/04-MassSpring/ext/19-MassSpringAuxinQuadratic/Init_blank_72cells_2023-09-07.mdxm', 'no', 'no')
Process.Model__CCF__01_Cell_Disk('10', '0.01', '.0001', '10', '10', '10', '0.1', 'Euler', 'Bi-conjugate Gradient Stabilized', '50', '1e-6', 'No', '.0001', 'Model/CCF/02 Mass Spring Solver', 'Model/CCF/10 Cell Tissue', 'Model/CCF/04 Mass Spring Growth', 'Model/CCF/05 Divide Cells', 'Model/CCF/06 Split Edges', 'Model/CCF/Auxin Gradient Model', 'Run_1_1-0_Linear_Quadratic_0-4_False_sim1_20231010', '140', '/home/ahr75/Documents/Shuyao/oofs_output/20231010', '10', 'True', 'False')
Process.Mesh__System__Open('Stack1', '/home/ahr75/Documents/Shuyao/oofs_2023-03-18/ext/02-CellModels/ext/04-MassSpring/ext/19-MassSpringAuxinQuadratic/Init_blank_72cells_2023-09-07.mdxm', 'no', 'no')
Process.Model__CCF__01_Cell_Disk('10', '0.01', '.0001', '10', '10', '10', '0.1', 'Euler', 'Bi-conjugate Gradient Stabilized', '50', '1e-6', 'No', '.0001', 'Model/CCF/02 Mass Spring Solver', 'Model/CCF/10 Cell Tissue', 'Model/CCF/04 Mass Spring Growth', 'Model/CCF/05 Divide Cells', 'Model/CCF/06 Split Edges', 'Model/CCF/Auxin Gradient Model', 'Run_1_1-0_Linear_Quadratic_0-8_False_sim1_20231010', '140', '/home/ahr75/Documents/Shuyao/oofs_output/20231010', '10', 'True', 'False')
Process.Mesh__System__Open('Stack1', '/home/ahr75/Documents/Shuyao/oofs_2023-03-18/ext/02-CellModels/ext/04-MassSpring/ext/19-MassSpringAuxinQuadratic/Init_blank_72cells_2023-09-07.mdxm', 'no', 'no')
Process.Model__CCF__01_Cell_Disk('10', '0.01', '.0001', '10', '10', '10', '0.1', 'Euler', 'Bi-conjugate Gradient Stabilized', '50', '1e-6', 'No', '.0001', 'Model/CCF/02 Mass Spring Solver', 'Model/CCF/10 Cell Tissue', 'Model/CCF/04 Mass Spring Growth', 'Model/CCF/05 Divide Cells', 'Model/CCF/06 Split Edges', 'Model/CCF/Auxin Gradient Model', 'Run_1_1-0_Linear_Quadratic_1-2_False_sim1_20231010', '140', '/home/ahr75/Documents/Shuyao/oofs_output/20231010', '10', 'True', 'False')
Process.Mesh__System__Open('Stack1', '/home/ahr75/Documents/Shuyao/oofs_2023-03-18/ext/02-CellModels/ext/04-MassSpring/ext/19-MassSpringAuxinQuadratic/Init_blank_72cells_2023-09-07.mdxm', 'no', 'no')
Process.Model__CCF__01_Cell_Disk('10', '0.01', '.0001', '10', '10', '10', '0.1', 'Euler', 'Bi-conjugate Gradient Stabilized', '50', '1e-6', 'No', '.0001', 'Model/CCF/02 Mass Spring Solver', 'Model/CCF/10 Cell Tissue', 'Model/CCF/04 Mass Spring Growth', 'Model/CCF/05 Divide Cells', 'Model/CCF/06 Split Edges', 'Model/CCF/Auxin Gradient Model', 'Run_0_0-1_Linear_Quadratic_0-4_False_sim1_20231010', '140', '/home/ahr75/Documents/Shuyao/oofs_output/20231010', '10', 'True', 'False')
Process.Mesh__System__Open('Stack1', '/home/ahr75/Documents/Shuyao/oofs_2023-03-18/ext/02-CellModels/ext/04-MassSpring/ext/19-MassSpringAuxinQuadratic/Init_blank_72cells_2023-09-07.mdxm', 'no', 'no')
Process.Model__CCF__01_Cell_Disk('10', '0.01', '.0001', '10', '10', '10', '0.1', 'Euler', 'Bi-conjugate Gradient Stabilized', '50', '1e-6', 'No', '.0001', 'Model/CCF/02 Mass Spring Solver', 'Model/CCF/10 Cell Tissue', 'Model/CCF/04 Mass Spring Growth', 'Model/CCF/05 Divide Cells', 'Model/CCF/06 Split Edges', 'Model/CCF/Auxin Gradient Model', 'Run_0_0-1_Linear_Quadratic_0-8_False_sim1_20231010', '140', '/home/ahr75/Documents/Shuyao/oofs_output/20231010', '10', 'True', 'False')
Process.Mesh__System__Open('Stack1', '/home/ahr75/Documents/Shuyao/oofs_2023-03-18/ext/02-CellModels/ext/04-MassSpring/ext/19-MassSpringAuxinQuadratic/Init_blank_72cells_2023-09-07.mdxm', 'no', 'no')
Process.Model__CCF__01_Cell_Disk('10', '0.01', '.0001', '10', '10', '10', '0.1', 'Euler', 'Bi-conjugate Gradient Stabilized', '50', '1e-6', 'No', '.0001', 'Model/CCF/02 Mass Spring Solver', 'Model/CCF/10 Cell Tissue', 'Model/CCF/04 Mass Spring Growth', 'Model/CCF/05 Divide Cells', 'Model/CCF/06 Split Edges', 'Model/CCF/Auxin Gradient Model', 'Run_0_0-1_Linear_Quadratic_1-2_False_sim1_20231010', '140', '/home/ahr75/Documents/Shuyao/oofs_output/20231010', '10', 'True', 'False')
Process.Mesh__System__Open('Stack1', '/home/ahr75/Documents/Shuyao/oofs_2023-03-18/ext/02-CellModels/ext/04-MassSpring/ext/19-MassSpringAuxinQuadratic/Init_blank_72cells_2023-09-07.mdxm', 'no', 'no')
Process.Model__CCF__01_Cell_Disk('10', '0.01', '.0001', '10', '10', '10', '0.1', 'Euler', 'Bi-conjugate Gradient Stabilized', '50', '1e-6', 'No', '.0001', 'Model/CCF/02 Mass Spring Solver', 'Model/CCF/10 Cell Tissue', 'Model/CCF/04 Mass Spring Growth', 'Model/CCF/05 Divide Cells', 'Model/CCF/06 Split Edges', 'Model/CCF/Auxin Gradient Model', 'Run_0_1-0_Linear_Quadratic_0-4_False_sim1_20231010', '140', '/home/ahr75/Documents/Shuyao/oofs_output/20231010', '10', 'True', 'False')
Process.Mesh__System__Open('Stack1', '/home/ahr75/Documents/Shuyao/oofs_2023-03-18/ext/02-CellModels/ext/04-MassSpring/ext/19-MassSpringAuxinQuadratic/Init_blank_72cells_2023-09-07.mdxm', 'no', 'no')
Process.Model__CCF__01_Cell_Disk('10', '0.01', '.0001', '10', '10', '10', '0.1', 'Euler', 'Bi-conjugate Gradient Stabilized', '50', '1e-6', 'No', '.0001', 'Model/CCF/02 Mass Spring Solver', 'Model/CCF/10 Cell Tissue', 'Model/CCF/04 Mass Spring Growth', 'Model/CCF/05 Divide Cells', 'Model/CCF/06 Split Edges', 'Model/CCF/Auxin Gradient Model', 'Run_0_1-0_Linear_Quadratic_0-8_False_sim1_20231010', '140', '/home/ahr75/Documents/Shuyao/oofs_output/20231010', '10', 'True', 'False')
Process.Mesh__System__Open('Stack1', '/home/ahr75/Documents/Shuyao/oofs_2023-03-18/ext/02-CellModels/ext/04-MassSpring/ext/19-MassSpringAuxinQuadratic/Init_blank_72cells_2023-09-07.mdxm', 'no', 'no')
Process.Model__CCF__01_Cell_Disk('10', '0.01', '.0001', '10', '10', '10', '0.1', 'Euler', 'Bi-conjugate Gradient Stabilized', '50', '1e-6', 'No', '.0001', 'Model/CCF/02 Mass Spring Solver', 'Model/CCF/10 Cell Tissue', 'Model/CCF/04 Mass Spring Growth', 'Model/CCF/05 Divide Cells', 'Model/CCF/06 Split Edges', 'Model/CCF/Auxin Gradient Model', 'Run_0_1-0_Linear_Quadratic_1-2_False_sim1_20231010', '140', '/home/ahr75/Documents/Shuyao/oofs_output/20231010', '10', 'True', 'False')
Process.Mesh__System__Open('Stack1', '/home/ahr75/Documents/Shuyao/oofs_2023-03-18/ext/02-CellModels/ext/04-MassSpring/ext/19-MassSpringAuxinQuadratic/Init_blank_72cells_2023-09-07.mdxm', 'no', 'no')
Process.Model__CCF__01_Cell_Disk('10', '0.01', '.0001', '10', '10', '10', '0.1', 'Euler', 'Bi-conjugate Gradient Stabilized', '50', '1e-6', 'No', '.0001', 'Model/CCF/02 Mass Spring Solver', 'Model/CCF/10 Cell Tissue', 'Model/CCF/04 Mass Spring Growth', 'Model/CCF/05 Divide Cells', 'Model/CCF/06 Split Edges', 'Model/CCF/Auxin Gradient Model', 'Run_1_0-1_Linear_Quadratic_0-4_False_sim2_20231010', '140', '/home/ahr75/Documents/Shuyao/oofs_output/20231010', '10', 'True', 'False')
Process.Mesh__System__Open('Stack1', '/home/ahr75/Documents/Shuyao/oofs_2023-03-18/ext/02-CellModels/ext/04-MassSpring/ext/19-MassSpringAuxinQuadratic/Init_blank_72cells_2023-09-07.mdxm', 'no', 'no')
Process.Model__CCF__01_Cell_Disk('10', '0.01', '.0001', '10', '10', '10', '0.1', 'Euler', 'Bi-conjugate Gradient Stabilized', '50', '1e-6', 'No', '.0001', 'Model/CCF/02 Mass Spring Solver', 'Model/CCF/10 Cell Tissue', 'Model/CCF/04 Mass Spring Growth', 'Model/CCF/05 Divide Cells', 'Model/CCF/06 Split Edges', 'Model/CCF/Auxin Gradient Model', 'Run_1_0-1_Linear_Quadratic_0-8_False_sim2_20231010', '140', '/home/ahr75/Documents/Shuyao/oofs_output/20231010', '10', 'True', 'False')
Process.Mesh__System__Open('Stack1', '/home/ahr75/Documents/Shuyao/oofs_2023-03-18/ext/02-CellModels/ext/04-MassSpring/ext/19-MassSpringAuxinQuadratic/Init_blank_72cells_2023-09-07.mdxm', 'no', 'no')
Process.Model__CCF__01_Cell_Disk('10', '0.01', '.0001', '10', '10', '10', '0.1', 'Euler', 'Bi-conjugate Gradient Stabilized', '50', '1e-6', 'No', '.0001', 'Model/CCF/02 Mass Spring Solver', 'Model/CCF/10 Cell Tissue', 'Model/CCF/04 Mass Spring Growth', 'Model/CCF/05 Divide Cells', 'Model/CCF/06 Split Edges', 'Model/CCF/Auxin Gradient Model', 'Run_1_0-1_Linear_Quadratic_1-2_False_sim2_20231010', '140', '/home/ahr75/Documents/Shuyao/oofs_output/20231010', '10', 'True', 'False')
Process.Mesh__System__Open('Stack1', '/home/ahr75/Documents/Shuyao/oofs_2023-03-18/ext/02-CellModels/ext/04-MassSpring/ext/19-MassSpringAuxinQuadratic/Init_blank_72cells_2023-09-07.mdxm', 'no', 'no')
Process.Model__CCF__01_Cell_Disk('10', '0.01', '.0001', '10', '10', '10', '0.1', 'Euler', 'Bi-conjugate Gradient Stabilized', '50', '1e-6', 'No', '.0001', 'Model/CCF/02 Mass Spring Solver', 'Model/CCF/10 Cell Tissue', 'Model/CCF/04 Mass Spring Growth', 'Model/CCF/05 Divide Cells', 'Model/CCF/06 Split Edges', 'Model/CCF/Auxin Gradient Model', 'Run_1_1-0_Linear_Quadratic_0-4_False_sim2_20231010', '140', '/home/ahr75/Documents/Shuyao/oofs_output/20231010', '10', 'True', 'False')
Process.Mesh__System__Open('Stack1', '/home/ahr75/Documents/Shuyao/oofs_2023-03-18/ext/02-CellModels/ext/04-MassSpring/ext/19-MassSpringAuxinQuadratic/Init_blank_72cells_2023-09-07.mdxm', 'no', 'no')
Process.Model__CCF__01_Cell_Disk('10', '0.01', '.0001', '10', '10', '10', '0.1', 'Euler', 'Bi-conjugate Gradient Stabilized', '50', '1e-6', 'No', '.0001', 'Model/CCF/02 Mass Spring Solver', 'Model/CCF/10 Cell Tissue', 'Model/CCF/04 Mass Spring Growth', 'Model/CCF/05 Divide Cells', 'Model/CCF/06 Split Edges', 'Model/CCF/Auxin Gradient Model', 'Run_1_1-0_Linear_Quadratic_0-8_False_sim2_20231010', '140', '/home/ahr75/Documents/Shuyao/oofs_output/20231010', '10', 'True', 'False')
Process.Mesh__System__Open('Stack1', '/home/ahr75/Documents/Shuyao/oofs_2023-03-18/ext/02-CellModels/ext/04-MassSpring/ext/19-MassSpringAuxinQuadratic/Init_blank_72cells_2023-09-07.mdxm', 'no', 'no')
Process.Model__CCF__01_Cell_Disk('10', '0.01', '.0001', '10', '10', '10', '0.1', 'Euler', 'Bi-conjugate Gradient Stabilized', '50', '1e-6', 'No', '.0001', 'Model/CCF/02 Mass Spring Solver', 'Model/CCF/10 Cell Tissue', 'Model/CCF/04 Mass Spring Growth', 'Model/CCF/05 Divide Cells', 'Model/CCF/06 Split Edges', 'Model/CCF/Auxin Gradient Model', 'Run_1_1-0_Linear_Quadratic_1-2_False_sim2_20231010', '140', '/home/ahr75/Documents/Shuyao/oofs_output/20231010', '10', 'True', 'False')
Process.Mesh__System__Open('Stack1', '/home/ahr75/Documents/Shuyao/oofs_2023-03-18/ext/02-CellModels/ext/04-MassSpring/ext/19-MassSpringAuxinQuadratic/Init_blank_72cells_2023-09-07.mdxm', 'no', 'no')
Process.Model__CCF__01_Cell_Disk('10', '0.01', '.0001', '10', '10', '10', '0.1', 'Euler', 'Bi-conjugate Gradient Stabilized', '50', '1e-6', 'No', '.0001', 'Model/CCF/02 Mass Spring Solver', 'Model/CCF/10 Cell Tissue', 'Model/CCF/04 Mass Spring Growth', 'Model/CCF/05 Divide Cells', 'Model/CCF/06 Split Edges', 'Model/CCF/Auxin Gradient Model', 'Run_0_0-1_Linear_Quadratic_0-4_False_sim2_20231010', '140', '/home/ahr75/Documents/Shuyao/oofs_output/20231010', '10', 'True', 'False')
Process.Mesh__System__Open('Stack1', '/home/ahr75/Documents/Shuyao/oofs_2023-03-18/ext/02-CellModels/ext/04-MassSpring/ext/19-MassSpringAuxinQuadratic/Init_blank_72cells_2023-09-07.mdxm', 'no', 'no')
Process.Model__CCF__01_Cell_Disk('10', '0.01', '.0001', '10', '10', '10', '0.1', 'Euler', 'Bi-conjugate Gradient Stabilized', '50', '1e-6', 'No', '.0001', 'Model/CCF/02 Mass Spring Solver', 'Model/CCF/10 Cell Tissue', 'Model/CCF/04 Mass Spring Growth', 'Model/CCF/05 Divide Cells', 'Model/CCF/06 Split Edges', 'Model/CCF/Auxin Gradient Model', 'Run_0_0-1_Linear_Quadratic_0-8_False_sim2_20231010', '140', '/home/ahr75/Documents/Shuyao/oofs_output/20231010', '10', 'True', 'False')
Process.Mesh__System__Open('Stack1', '/home/ahr75/Documents/Shuyao/oofs_2023-03-18/ext/02-CellModels/ext/04-MassSpring/ext/19-MassSpringAuxinQuadratic/Init_blank_72cells_2023-09-07.mdxm', 'no', 'no')
Process.Model__CCF__01_Cell_Disk('10', '0.01', '.0001', '10', '10', '10', '0.1', 'Euler', 'Bi-conjugate Gradient Stabilized', '50', '1e-6', 'No', '.0001', 'Model/CCF/02 Mass Spring Solver', 'Model/CCF/10 Cell Tissue', 'Model/CCF/04 Mass Spring Growth', 'Model/CCF/05 Divide Cells', 'Model/CCF/06 Split Edges', 'Model/CCF/Auxin Gradient Model', 'Run_0_0-1_Linear_Quadratic_1-2_False_sim2_20231010', '140', '/home/ahr75/Documents/Shuyao/oofs_output/20231010', '10', 'True', 'False')
Process.Mesh__System__Open('Stack1', '/home/ahr75/Documents/Shuyao/oofs_2023-03-18/ext/02-CellModels/ext/04-MassSpring/ext/19-MassSpringAuxinQuadratic/Init_blank_72cells_2023-09-07.mdxm', 'no', 'no')
Process.Model__CCF__01_Cell_Disk('10', '0.01', '.0001', '10', '10', '10', '0.1', 'Euler', 'Bi-conjugate Gradient Stabilized', '50', '1e-6', 'No', '.0001', 'Model/CCF/02 Mass Spring Solver', 'Model/CCF/10 Cell Tissue', 'Model/CCF/04 Mass Spring Growth', 'Model/CCF/05 Divide Cells', 'Model/CCF/06 Split Edges', 'Model/CCF/Auxin Gradient Model', 'Run_0_1-0_Linear_Quadratic_0-4_False_sim2_20231010', '140', '/home/ahr75/Documents/Shuyao/oofs_output/20231010', '10', 'True', 'False')
Process.Mesh__System__Open('Stack1', '/home/ahr75/Documents/Shuyao/oofs_2023-03-18/ext/02-CellModels/ext/04-MassSpring/ext/19-MassSpringAuxinQuadratic/Init_blank_72cells_2023-09-07.mdxm', 'no', 'no')
Process.Model__CCF__01_Cell_Disk('10', '0.01', '.0001', '10', '10', '10', '0.1', 'Euler', 'Bi-conjugate Gradient Stabilized', '50', '1e-6', 'No', '.0001', 'Model/CCF/02 Mass Spring Solver', 'Model/CCF/10 Cell Tissue', 'Model/CCF/04 Mass Spring Growth', 'Model/CCF/05 Divide Cells', 'Model/CCF/06 Split Edges', 'Model/CCF/Auxin Gradient Model', 'Run_0_1-0_Linear_Quadratic_0-8_False_sim2_20231010', '140', '/home/ahr75/Documents/Shuyao/oofs_output/20231010', '10', 'True', 'False')
Process.Mesh__System__Open('Stack1', '/home/ahr75/Documents/Shuyao/oofs_2023-03-18/ext/02-CellModels/ext/04-MassSpring/ext/19-MassSpringAuxinQuadratic/Init_blank_72cells_2023-09-07.mdxm', 'no', 'no')
Process.Model__CCF__01_Cell_Disk('10', '0.01', '.0001', '10', '10', '10', '0.1', 'Euler', 'Bi-conjugate Gradient Stabilized', '50', '1e-6', 'No', '.0001', 'Model/CCF/02 Mass Spring Solver', 'Model/CCF/10 Cell Tissue', 'Model/CCF/04 Mass Spring Growth', 'Model/CCF/05 Divide Cells', 'Model/CCF/06 Split Edges', 'Model/CCF/Auxin Gradient Model', 'Run_0_1-0_Linear_Quadratic_1-2_False_sim2_20231010', '140', '/home/ahr75/Documents/Shuyao/oofs_output/20231010', '10', 'True', 'False')
Process.Mesh__System__Open('Stack1', '/home/ahr75/Documents/Shuyao/oofs_2023-03-18/ext/02-CellModels/ext/04-MassSpring/ext/19-MassSpringAuxinQuadratic/Init_blank_72cells_2023-09-07.mdxm', 'no', 'no')
Process.Model__CCF__01_Cell_Disk('10', '0.01', '.0001', '10', '10', '10', '0.1', 'Euler', 'Bi-conjugate Gradient Stabilized', '50', '1e-6', 'No', '.0001', 'Model/CCF/02 Mass Spring Solver', 'Model/CCF/10 Cell Tissue', 'Model/CCF/04 Mass Spring Growth', 'Model/CCF/05 Divide Cells', 'Model/CCF/06 Split Edges', 'Model/CCF/Auxin Gradient Model', 'Run_1_0-1_Linear_Quadratic_0-4_False_sim3_20231010', '140', '/home/ahr75/Documents/Shuyao/oofs_output/20231010', '10', 'True', 'False')
Process.Mesh__System__Open('Stack1', '/home/ahr75/Documents/Shuyao/oofs_2023-03-18/ext/02-CellModels/ext/04-MassSpring/ext/19-MassSpringAuxinQuadratic/Init_blank_72cells_2023-09-07.mdxm', 'no', 'no')
Process.Model__CCF__01_Cell_Disk('10', '0.01', '.0001', '10', '10', '10', '0.1', 'Euler', 'Bi-conjugate Gradient Stabilized', '50', '1e-6', 'No', '.0001', 'Model/CCF/02 Mass Spring Solver', 'Model/CCF/10 Cell Tissue', 'Model/CCF/04 Mass Spring Growth', 'Model/CCF/05 Divide Cells', 'Model/CCF/06 Split Edges', 'Model/CCF/Auxin Gradient Model', 'Run_1_0-1_Linear_Quadratic_0-8_False_sim3_20231010', '140', '/home/ahr75/Documents/Shuyao/oofs_output/20231010', '10', 'True', 'False')
Process.Mesh__System__Open('Stack1', '/home/ahr75/Documents/Shuyao/oofs_2023-03-18/ext/02-CellModels/ext/04-MassSpring/ext/19-MassSpringAuxinQuadratic/Init_blank_72cells_2023-09-07.mdxm', 'no', 'no')
Process.Model__CCF__01_Cell_Disk('10', '0.01', '.0001', '10', '10', '10', '0.1', 'Euler', 'Bi-conjugate Gradient Stabilized', '50', '1e-6', 'No', '.0001', 'Model/CCF/02 Mass Spring Solver', 'Model/CCF/10 Cell Tissue', 'Model/CCF/04 Mass Spring Growth', 'Model/CCF/05 Divide Cells', 'Model/CCF/06 Split Edges', 'Model/CCF/Auxin Gradient Model', 'Run_1_0-1_Linear_Quadratic_1-2_False_sim3_20231010', '140', '/home/ahr75/Documents/Shuyao/oofs_output/20231010', '10', 'True', 'False')
Process.Mesh__System__Open('Stack1', '/home/ahr75/Documents/Shuyao/oofs_2023-03-18/ext/02-CellModels/ext/04-MassSpring/ext/19-MassSpringAuxinQuadratic/Init_blank_72cells_2023-09-07.mdxm', 'no', 'no')
Process.Model__CCF__01_Cell_Disk('10', '0.01', '.0001', '10', '10', '10', '0.1', 'Euler', 'Bi-conjugate Gradient Stabilized', '50', '1e-6', 'No', '.0001', 'Model/CCF/02 Mass Spring Solver', 'Model/CCF/10 Cell Tissue', 'Model/CCF/04 Mass Spring Growth', 'Model/CCF/05 Divide Cells', 'Model/CCF/06 Split Edges', 'Model/CCF/Auxin Gradient Model', 'Run_1_1-0_Linear_Quadratic_0-4_False_sim3_20231010', '140', '/home/ahr75/Documents/Shuyao/oofs_output/20231010', '10', 'True', 'False')
Process.Mesh__System__Open('Stack1', '/home/ahr75/Documents/Shuyao/oofs_2023-03-18/ext/02-CellModels/ext/04-MassSpring/ext/19-MassSpringAuxinQuadratic/Init_blank_72cells_2023-09-07.mdxm', 'no', 'no')
Process.Model__CCF__01_Cell_Disk('10', '0.01', '.0001', '10', '10', '10', '0.1', 'Euler', 'Bi-conjugate Gradient Stabilized', '50', '1e-6', 'No', '.0001', 'Model/CCF/02 Mass Spring Solver', 'Model/CCF/10 Cell Tissue', 'Model/CCF/04 Mass Spring Growth', 'Model/CCF/05 Divide Cells', 'Model/CCF/06 Split Edges', 'Model/CCF/Auxin Gradient Model', 'Run_1_1-0_Linear_Quadratic_0-8_False_sim3_20231010', '140', '/home/ahr75/Documents/Shuyao/oofs_output/20231010', '10', 'True', 'False')
Process.Mesh__System__Open('Stack1', '/home/ahr75/Documents/Shuyao/oofs_2023-03-18/ext/02-CellModels/ext/04-MassSpring/ext/19-MassSpringAuxinQuadratic/Init_blank_72cells_2023-09-07.mdxm', 'no', 'no')
Process.Model__CCF__01_Cell_Disk('10', '0.01', '.0001', '10', '10', '10', '0.1', 'Euler', 'Bi-conjugate Gradient Stabilized', '50', '1e-6', 'No', '.0001', 'Model/CCF/02 Mass Spring Solver', 'Model/CCF/10 Cell Tissue', 'Model/CCF/04 Mass Spring Growth', 'Model/CCF/05 Divide Cells', 'Model/CCF/06 Split Edges', 'Model/CCF/Auxin Gradient Model', 'Run_1_1-0_Linear_Quadratic_1-2_False_sim3_20231010', '140', '/home/ahr75/Documents/Shuyao/oofs_output/20231010', '10', 'True', 'False')
Process.Mesh__System__Open('Stack1', '/home/ahr75/Documents/Shuyao/oofs_2023-03-18/ext/02-CellModels/ext/04-MassSpring/ext/19-MassSpringAuxinQuadratic/Init_blank_72cells_2023-09-07.mdxm', 'no', 'no')
Process.Model__CCF__01_Cell_Disk('10', '0.01', '.0001', '10', '10', '10', '0.1', 'Euler', 'Bi-conjugate Gradient Stabilized', '50', '1e-6', 'No', '.0001', 'Model/CCF/02 Mass Spring Solver', 'Model/CCF/10 Cell Tissue', 'Model/CCF/04 Mass Spring Growth', 'Model/CCF/05 Divide Cells', 'Model/CCF/06 Split Edges', 'Model/CCF/Auxin Gradient Model', 'Run_0_0-1_Linear_Quadratic_0-4_False_sim3_20231010', '140', '/home/ahr75/Documents/Shuyao/oofs_output/20231010', '10', 'True', 'False')
Process.Mesh__System__Open('Stack1', '/home/ahr75/Documents/Shuyao/oofs_2023-03-18/ext/02-CellModels/ext/04-MassSpring/ext/19-MassSpringAuxinQuadratic/Init_blank_72cells_2023-09-07.mdxm', 'no', 'no')
Process.Model__CCF__01_Cell_Disk('10', '0.01', '.0001', '10', '10', '10', '0.1', 'Euler', 'Bi-conjugate Gradient Stabilized', '50', '1e-6', 'No', '.0001', 'Model/CCF/02 Mass Spring Solver', 'Model/CCF/10 Cell Tissue', 'Model/CCF/04 Mass Spring Growth', 'Model/CCF/05 Divide Cells', 'Model/CCF/06 Split Edges', 'Model/CCF/Auxin Gradient Model', 'Run_0_0-1_Linear_Quadratic_0-8_False_sim3_20231010', '140', '/home/ahr75/Documents/Shuyao/oofs_output/20231010', '10', 'True', 'False')
Process.Mesh__System__Open('Stack1', '/home/ahr75/Documents/Shuyao/oofs_2023-03-18/ext/02-CellModels/ext/04-MassSpring/ext/19-MassSpringAuxinQuadratic/Init_blank_72cells_2023-09-07.mdxm', 'no', 'no')
Process.Model__CCF__01_Cell_Disk('10', '0.01', '.0001', '10', '10', '10', '0.1', 'Euler', 'Bi-conjugate Gradient Stabilized', '50', '1e-6', 'No', '.0001', 'Model/CCF/02 Mass Spring Solver', 'Model/CCF/10 Cell Tissue', 'Model/CCF/04 Mass Spring Growth', 'Model/CCF/05 Divide Cells', 'Model/CCF/06 Split Edges', 'Model/CCF/Auxin Gradient Model', 'Run_0_0-1_Linear_Quadratic_1-2_False_sim3_20231010', '140', '/home/ahr75/Documents/Shuyao/oofs_output/20231010', '10', 'True', 'False')
Process.Mesh__System__Open('Stack1', '/home/ahr75/Documents/Shuyao/oofs_2023-03-18/ext/02-CellModels/ext/04-MassSpring/ext/19-MassSpringAuxinQuadratic/Init_blank_72cells_2023-09-07.mdxm', 'no', 'no')
Process.Model__CCF__01_Cell_Disk('10', '0.01', '.0001', '10', '10', '10', '0.1', 'Euler', 'Bi-conjugate Gradient Stabilized', '50', '1e-6', 'No', '.0001', 'Model/CCF/02 Mass Spring Solver', 'Model/CCF/10 Cell Tissue', 'Model/CCF/04 Mass Spring Growth', 'Model/CCF/05 Divide Cells', 'Model/CCF/06 Split Edges', 'Model/CCF/Auxin Gradient Model', 'Run_0_1-0_Linear_Quadratic_0-4_False_sim3_20231010', '140', '/home/ahr75/Documents/Shuyao/oofs_output/20231010', '10', 'True', 'False')
Process.Mesh__System__Open('Stack1', '/home/ahr75/Documents/Shuyao/oofs_2023-03-18/ext/02-CellModels/ext/04-MassSpring/ext/19-MassSpringAuxinQuadratic/Init_blank_72cells_2023-09-07.mdxm', 'no', 'no')
Process.Model__CCF__01_Cell_Disk('10', '0.01', '.0001', '10', '10', '10', '0.1', 'Euler', 'Bi-conjugate Gradient Stabilized', '50', '1e-6', 'No', '.0001', 'Model/CCF/02 Mass Spring Solver', 'Model/CCF/10 Cell Tissue', 'Model/CCF/04 Mass Spring Growth', 'Model/CCF/05 Divide Cells', 'Model/CCF/06 Split Edges', 'Model/CCF/Auxin Gradient Model', 'Run_0_1-0_Linear_Quadratic_0-8_False_sim3_20231010', '140', '/home/ahr75/Documents/Shuyao/oofs_output/20231010', '10', 'True', 'False')
Process.Mesh__System__Open('Stack1', '/home/ahr75/Documents/Shuyao/oofs_2023-03-18/ext/02-CellModels/ext/04-MassSpring/ext/19-MassSpringAuxinQuadratic/Init_blank_72cells_2023-09-07.mdxm', 'no', 'no')
Process.Model__CCF__01_Cell_Disk('10', '0.01', '.0001', '10', '10', '10', '0.1', 'Euler', 'Bi-conjugate Gradient Stabilized', '50', '1e-6', 'No', '.0001', 'Model/CCF/02 Mass Spring Solver', 'Model/CCF/10 Cell Tissue', 'Model/CCF/04 Mass Spring Growth', 'Model/CCF/05 Divide Cells', 'Model/CCF/06 Split Edges', 'Model/CCF/Auxin Gradient Model', 'Run_0_1-0_Linear_Quadratic_1-2_False_sim3_20231010', '140', '/home/ahr75/Documents/Shuyao/oofs_output/20231010', '10', 'True', 'False')
Process.Mesh__System__Open('Stack1', '/home/ahr75/Documents/Shuyao/oofs_2023-03-18/ext/02-CellModels/ext/04-MassSpring/ext/19-MassSpringAuxinQuadratic/Init_blank_72cells_2023-09-07.mdxm', 'no', 'no')
Process.Model__CCF__01_Cell_Disk('10', '0.01', '.0001', '10', '10', '10', '0.1', 'Euler', 'Bi-conjugate Gradient Stabilized', '50', '1e-6', 'No', '.0001', 'Model/CCF/02 Mass Spring Solver', 'Model/CCF/10 Cell Tissue', 'Model/CCF/04 Mass Spring Growth', 'Model/CCF/05 Divide Cells', 'Model/CCF/06 Split Edges', 'Model/CCF/Auxin Gradient Model', 'Run_1_0-1_Linear_Quadratic_0-4_False_sim4_20231010', '140', '/home/ahr75/Documents/Shuyao/oofs_output/20231010', '10', 'True', 'False')
Process.Mesh__System__Open('Stack1', '/home/ahr75/Documents/Shuyao/oofs_2023-03-18/ext/02-CellModels/ext/04-MassSpring/ext/19-MassSpringAuxinQuadratic/Init_blank_72cells_2023-09-07.mdxm', 'no', 'no')
Process.Model__CCF__01_Cell_Disk('10', '0.01', '.0001', '10', '10', '10', '0.1', 'Euler', 'Bi-conjugate Gradient Stabilized', '50', '1e-6', 'No', '.0001', 'Model/CCF/02 Mass Spring Solver', 'Model/CCF/10 Cell Tissue', 'Model/CCF/04 Mass Spring Growth', 'Model/CCF/05 Divide Cells', 'Model/CCF/06 Split Edges', 'Model/CCF/Auxin Gradient Model', 'Run_1_0-1_Linear_Quadratic_0-8_False_sim4_20231010', '140', '/home/ahr75/Documents/Shuyao/oofs_output/20231010', '10', 'True', 'False')
Process.Mesh__System__Open('Stack1', '/home/ahr75/Documents/Shuyao/oofs_2023-03-18/ext/02-CellModels/ext/04-MassSpring/ext/19-MassSpringAuxinQuadratic/Init_blank_72cells_2023-09-07.mdxm', 'no', 'no')
Process.Model__CCF__01_Cell_Disk('10', '0.01', '.0001', '10', '10', '10', '0.1', 'Euler', 'Bi-conjugate Gradient Stabilized', '50', '1e-6', 'No', '.0001', 'Model/CCF/02 Mass Spring Solver', 'Model/CCF/10 Cell Tissue', 'Model/CCF/04 Mass Spring Growth', 'Model/CCF/05 Divide Cells', 'Model/CCF/06 Split Edges', 'Model/CCF/Auxin Gradient Model', 'Run_1_0-1_Linear_Quadratic_1-2_False_sim4_20231010', '140', '/home/ahr75/Documents/Shuyao/oofs_output/20231010', '10', 'True', 'False')
Process.Mesh__System__Open('Stack1', '/home/ahr75/Documents/Shuyao/oofs_2023-03-18/ext/02-CellModels/ext/04-MassSpring/ext/19-MassSpringAuxinQuadratic/Init_blank_72cells_2023-09-07.mdxm', 'no', 'no')
Process.Model__CCF__01_Cell_Disk('10', '0.01', '.0001', '10', '10', '10', '0.1', 'Euler', 'Bi-conjugate Gradient Stabilized', '50', '1e-6', 'No', '.0001', 'Model/CCF/02 Mass Spring Solver', 'Model/CCF/10 Cell Tissue', 'Model/CCF/04 Mass Spring Growth', 'Model/CCF/05 Divide Cells', 'Model/CCF/06 Split Edges', 'Model/CCF/Auxin Gradient Model', 'Run_1_1-0_Linear_Quadratic_0-4_False_sim4_20231010', '140', '/home/ahr75/Documents/Shuyao/oofs_output/20231010', '10', 'True', 'False')
Process.Mesh__System__Open('Stack1', '/home/ahr75/Documents/Shuyao/oofs_2023-03-18/ext/02-CellModels/ext/04-MassSpring/ext/19-MassSpringAuxinQuadratic/Init_blank_72cells_2023-09-07.mdxm', 'no', 'no')
Process.Model__CCF__01_Cell_Disk('10', '0.01', '.0001', '10', '10', '10', '0.1', 'Euler', 'Bi-conjugate Gradient Stabilized', '50', '1e-6', 'No', '.0001', 'Model/CCF/02 Mass Spring Solver', 'Model/CCF/10 Cell Tissue', 'Model/CCF/04 Mass Spring Growth', 'Model/CCF/05 Divide Cells', 'Model/CCF/06 Split Edges', 'Model/CCF/Auxin Gradient Model', 'Run_1_1-0_Linear_Quadratic_0-8_False_sim4_20231010', '140', '/home/ahr75/Documents/Shuyao/oofs_output/20231010', '10', 'True', 'False')
Process.Mesh__System__Open('Stack1', '/home/ahr75/Documents/Shuyao/oofs_2023-03-18/ext/02-CellModels/ext/04-MassSpring/ext/19-MassSpringAuxinQuadratic/Init_blank_72cells_2023-09-07.mdxm', 'no', 'no')
Process.Model__CCF__01_Cell_Disk('10', '0.01', '.0001', '10', '10', '10', '0.1', 'Euler', 'Bi-conjugate Gradient Stabilized', '50', '1e-6', 'No', '.0001', 'Model/CCF/02 Mass Spring Solver', 'Model/CCF/10 Cell Tissue', 'Model/CCF/04 Mass Spring Growth', 'Model/CCF/05 Divide Cells', 'Model/CCF/06 Split Edges', 'Model/CCF/Auxin Gradient Model', 'Run_1_1-0_Linear_Quadratic_1-2_False_sim4_20231010', '140', '/home/ahr75/Documents/Shuyao/oofs_output/20231010', '10', 'True', 'False')
Process.Mesh__System__Open('Stack1', '/home/ahr75/Documents/Shuyao/oofs_2023-03-18/ext/02-CellModels/ext/04-MassSpring/ext/19-MassSpringAuxinQuadratic/Init_blank_72cells_2023-09-07.mdxm', 'no', 'no')
Process.Model__CCF__01_Cell_Disk('10', '0.01', '.0001', '10', '10', '10', '0.1', 'Euler', 'Bi-conjugate Gradient Stabilized', '50', '1e-6', 'No', '.0001', 'Model/CCF/02 Mass Spring Solver', 'Model/CCF/10 Cell Tissue', 'Model/CCF/04 Mass Spring Growth', 'Model/CCF/05 Divide Cells', 'Model/CCF/06 Split Edges', 'Model/CCF/Auxin Gradient Model', 'Run_0_0-1_Linear_Quadratic_0-4_False_sim4_20231010', '140', '/home/ahr75/Documents/Shuyao/oofs_output/20231010', '10', 'True', 'False')
Process.Mesh__System__Open('Stack1', '/home/ahr75/Documents/Shuyao/oofs_2023-03-18/ext/02-CellModels/ext/04-MassSpring/ext/19-MassSpringAuxinQuadratic/Init_blank_72cells_2023-09-07.mdxm', 'no', 'no')
Process.Model__CCF__01_Cell_Disk('10', '0.01', '.0001', '10', '10', '10', '0.1', 'Euler', 'Bi-conjugate Gradient Stabilized', '50', '1e-6', 'No', '.0001', 'Model/CCF/02 Mass Spring Solver', 'Model/CCF/10 Cell Tissue', 'Model/CCF/04 Mass Spring Growth', 'Model/CCF/05 Divide Cells', 'Model/CCF/06 Split Edges', 'Model/CCF/Auxin Gradient Model', 'Run_0_0-1_Linear_Quadratic_0-8_False_sim4_20231010', '140', '/home/ahr75/Documents/Shuyao/oofs_output/20231010', '10', 'True', 'False')
Process.Mesh__System__Open('Stack1', '/home/ahr75/Documents/Shuyao/oofs_2023-03-18/ext/02-CellModels/ext/04-MassSpring/ext/19-MassSpringAuxinQuadratic/Init_blank_72cells_2023-09-07.mdxm', 'no', 'no')
Process.Model__CCF__01_Cell_Disk('10', '0.01', '.0001', '10', '10', '10', '0.1', 'Euler', 'Bi-conjugate Gradient Stabilized', '50', '1e-6', 'No', '.0001', 'Model/CCF/02 Mass Spring Solver', 'Model/CCF/10 Cell Tissue', 'Model/CCF/04 Mass Spring Growth', 'Model/CCF/05 Divide Cells', 'Model/CCF/06 Split Edges', 'Model/CCF/Auxin Gradient Model', 'Run_0_0-1_Linear_Quadratic_1-2_False_sim4_20231010', '140', '/home/ahr75/Documents/Shuyao/oofs_output/20231010', '10', 'True', 'False')
Process.Mesh__System__Open('Stack1', '/home/ahr75/Documents/Shuyao/oofs_2023-03-18/ext/02-CellModels/ext/04-MassSpring/ext/19-MassSpringAuxinQuadratic/Init_blank_72cells_2023-09-07.mdxm', 'no', 'no')
Process.Model__CCF__01_Cell_Disk('10', '0.01', '.0001', '10', '10', '10', '0.1', 'Euler', 'Bi-conjugate Gradient Stabilized', '50', '1e-6', 'No', '.0001', 'Model/CCF/02 Mass Spring Solver', 'Model/CCF/10 Cell Tissue', 'Model/CCF/04 Mass Spring Growth', 'Model/CCF/05 Divide Cells', 'Model/CCF/06 Split Edges', 'Model/CCF/Auxin Gradient Model', 'Run_0_1-0_Linear_Quadratic_0-4_False_sim4_20231010', '140', '/home/ahr75/Documents/Shuyao/oofs_output/20231010', '10', 'True', 'False')
Process.Mesh__System__Open('Stack1', '/home/ahr75/Documents/Shuyao/oofs_2023-03-18/ext/02-CellModels/ext/04-MassSpring/ext/19-MassSpringAuxinQuadratic/Init_blank_72cells_2023-09-07.mdxm', 'no', 'no')
Process.Model__CCF__01_Cell_Disk('10', '0.01', '.0001', '10', '10', '10', '0.1', 'Euler', 'Bi-conjugate Gradient Stabilized', '50', '1e-6', 'No', '.0001', 'Model/CCF/02 Mass Spring Solver', 'Model/CCF/10 Cell Tissue', 'Model/CCF/04 Mass Spring Growth', 'Model/CCF/05 Divide Cells', 'Model/CCF/06 Split Edges', 'Model/CCF/Auxin Gradient Model', 'Run_0_1-0_Linear_Quadratic_0-8_False_sim4_20231010', '140', '/home/ahr75/Documents/Shuyao/oofs_output/20231010', '10', 'True', 'False')
Process.Mesh__System__Open('Stack1', '/home/ahr75/Documents/Shuyao/oofs_2023-03-18/ext/02-CellModels/ext/04-MassSpring/ext/19-MassSpringAuxinQuadratic/Init_blank_72cells_2023-09-07.mdxm', 'no', 'no')
Process.Model__CCF__01_Cell_Disk('10', '0.01', '.0001', '10', '10', '10', '0.1', 'Euler', 'Bi-conjugate Gradient Stabilized', '50', '1e-6', 'No', '.0001', 'Model/CCF/02 Mass Spring Solver', 'Model/CCF/10 Cell Tissue', 'Model/CCF/04 Mass Spring Growth', 'Model/CCF/05 Divide Cells', 'Model/CCF/06 Split Edges', 'Model/CCF/Auxin Gradient Model', 'Run_0_1-0_Linear_Quadratic_1-2_False_sim4_20231010', '140', '/home/ahr75/Documents/Shuyao/oofs_output/20231010', '10', 'True', 'False')
Process.Mesh__System__Open('Stack1', '/home/ahr75/Documents/Shuyao/oofs_2023-03-18/ext/02-CellModels/ext/04-MassSpring/ext/19-MassSpringAuxinQuadratic/Init_blank_72cells_2023-09-07.mdxm', 'no', 'no')
Process.Model__CCF__01_Cell_Disk('10', '0.01', '.0001', '10', '10', '10', '0.1', 'Euler', 'Bi-conjugate Gradient Stabilized', '50', '1e-6', 'No', '.0001', 'Model/CCF/02 Mass Spring Solver', 'Model/CCF/10 Cell Tissue', 'Model/CCF/04 Mass Spring Growth', 'Model/CCF/05 Divide Cells', 'Model/CCF/06 Split Edges', 'Model/CCF/Auxin Gradient Model', 'Run_1_0-1_Linear_Quadratic_0-4_False_sim5_20231010', '140', '/home/ahr75/Documents/Shuyao/oofs_output/20231010', '10', 'True', 'False')
Process.Mesh__System__Open('Stack1', '/home/ahr75/Documents/Shuyao/oofs_2023-03-18/ext/02-CellModels/ext/04-MassSpring/ext/19-MassSpringAuxinQuadratic/Init_blank_72cells_2023-09-07.mdxm', 'no', 'no')
Process.Model__CCF__01_Cell_Disk('10', '0.01', '.0001', '10', '10', '10', '0.1', 'Euler', 'Bi-conjugate Gradient Stabilized', '50', '1e-6', 'No', '.0001', 'Model/CCF/02 Mass Spring Solver', 'Model/CCF/10 Cell Tissue', 'Model/CCF/04 Mass Spring Growth', 'Model/CCF/05 Divide Cells', 'Model/CCF/06 Split Edges', 'Model/CCF/Auxin Gradient Model', 'Run_1_0-1_Linear_Quadratic_0-8_False_sim5_20231010', '140', '/home/ahr75/Documents/Shuyao/oofs_output/20231010', '10', 'True', 'False')
Process.Mesh__System__Open('Stack1', '/home/ahr75/Documents/Shuyao/oofs_2023-03-18/ext/02-CellModels/ext/04-MassSpring/ext/19-MassSpringAuxinQuadratic/Init_blank_72cells_2023-09-07.mdxm', 'no', 'no')
Process.Model__CCF__01_Cell_Disk('10', '0.01', '.0001', '10', '10', '10', '0.1', 'Euler', 'Bi-conjugate Gradient Stabilized', '50', '1e-6', 'No', '.0001', 'Model/CCF/02 Mass Spring Solver', 'Model/CCF/10 Cell Tissue', 'Model/CCF/04 Mass Spring Growth', 'Model/CCF/05 Divide Cells', 'Model/CCF/06 Split Edges', 'Model/CCF/Auxin Gradient Model', 'Run_1_0-1_Linear_Quadratic_1-2_False_sim5_20231010', '140', '/home/ahr75/Documents/Shuyao/oofs_output/20231010', '10', 'True', 'False')
Process.Mesh__System__Open('Stack1', '/home/ahr75/Documents/Shuyao/oofs_2023-03-18/ext/02-CellModels/ext/04-MassSpring/ext/19-MassSpringAuxinQuadratic/Init_blank_72cells_2023-09-07.mdxm', 'no', 'no')
Process.Model__CCF__01_Cell_Disk('10', '0.01', '.0001', '10', '10', '10', '0.1', 'Euler', 'Bi-conjugate Gradient Stabilized', '50', '1e-6', 'No', '.0001', 'Model/CCF/02 Mass Spring Solver', 'Model/CCF/10 Cell Tissue', 'Model/CCF/04 Mass Spring Growth', 'Model/CCF/05 Divide Cells', 'Model/CCF/06 Split Edges', 'Model/CCF/Auxin Gradient Model', 'Run_1_1-0_Linear_Quadratic_0-4_False_sim5_20231010', '140', '/home/ahr75/Documents/Shuyao/oofs_output/20231010', '10', 'True', 'False')
Process.Mesh__System__Open('Stack1', '/home/ahr75/Documents/Shuyao/oofs_2023-03-18/ext/02-CellModels/ext/04-MassSpring/ext/19-MassSpringAuxinQuadratic/Init_blank_72cells_2023-09-07.mdxm', 'no', 'no')
Process.Model__CCF__01_Cell_Disk('10', '0.01', '.0001', '10', '10', '10', '0.1', 'Euler', 'Bi-conjugate Gradient Stabilized', '50', '1e-6', 'No', '.0001', 'Model/CCF/02 Mass Spring Solver', 'Model/CCF/10 Cell Tissue', 'Model/CCF/04 Mass Spring Growth', 'Model/CCF/05 Divide Cells', 'Model/CCF/06 Split Edges', 'Model/CCF/Auxin Gradient Model', 'Run_1_1-0_Linear_Quadratic_0-8_False_sim5_20231010', '140', '/home/ahr75/Documents/Shuyao/oofs_output/20231010', '10', 'True', 'False')
Process.Mesh__System__Open('Stack1', '/home/ahr75/Documents/Shuyao/oofs_2023-03-18/ext/02-CellModels/ext/04-MassSpring/ext/19-MassSpringAuxinQuadratic/Init_blank_72cells_2023-09-07.mdxm', 'no', 'no')
Process.Model__CCF__01_Cell_Disk('10', '0.01', '.0001', '10', '10', '10', '0.1', 'Euler', 'Bi-conjugate Gradient Stabilized', '50', '1e-6', 'No', '.0001', 'Model/CCF/02 Mass Spring Solver', 'Model/CCF/10 Cell Tissue', 'Model/CCF/04 Mass Spring Growth', 'Model/CCF/05 Divide Cells', 'Model/CCF/06 Split Edges', 'Model/CCF/Auxin Gradient Model', 'Run_1_1-0_Linear_Quadratic_1-2_False_sim5_20231010', '140', '/home/ahr75/Documents/Shuyao/oofs_output/20231010', '10', 'True', 'False')
Process.Mesh__System__Open('Stack1', '/home/ahr75/Documents/Shuyao/oofs_2023-03-18/ext/02-CellModels/ext/04-MassSpring/ext/19-MassSpringAuxinQuadratic/Init_blank_72cells_2023-09-07.mdxm', 'no', 'no')
Process.Model__CCF__01_Cell_Disk('10', '0.01', '.0001', '10', '10', '10', '0.1', 'Euler', 'Bi-conjugate Gradient Stabilized', '50', '1e-6', 'No', '.0001', 'Model/CCF/02 Mass Spring Solver', 'Model/CCF/10 Cell Tissue', 'Model/CCF/04 Mass Spring Growth', 'Model/CCF/05 Divide Cells', 'Model/CCF/06 Split Edges', 'Model/CCF/Auxin Gradient Model', 'Run_0_0-1_Linear_Quadratic_0-4_False_sim5_20231010', '140', '/home/ahr75/Documents/Shuyao/oofs_output/20231010', '10', 'True', 'False')
Process.Mesh__System__Open('Stack1', '/home/ahr75/Documents/Shuyao/oofs_2023-03-18/ext/02-CellModels/ext/04-MassSpring/ext/19-MassSpringAuxinQuadratic/Init_blank_72cells_2023-09-07.mdxm', 'no', 'no')
Process.Model__CCF__01_Cell_Disk('10', '0.01', '.0001', '10', '10', '10', '0.1', 'Euler', 'Bi-conjugate Gradient Stabilized', '50', '1e-6', 'No', '.0001', 'Model/CCF/02 Mass Spring Solver', 'Model/CCF/10 Cell Tissue', 'Model/CCF/04 Mass Spring Growth', 'Model/CCF/05 Divide Cells', 'Model/CCF/06 Split Edges', 'Model/CCF/Auxin Gradient Model', 'Run_0_0-1_Linear_Quadratic_0-8_False_sim5_20231010', '140', '/home/ahr75/Documents/Shuyao/oofs_output/20231010', '10', 'True', 'False')
Process.Mesh__System__Open('Stack1', '/home/ahr75/Documents/Shuyao/oofs_2023-03-18/ext/02-CellModels/ext/04-MassSpring/ext/19-MassSpringAuxinQuadratic/Init_blank_72cells_2023-09-07.mdxm', 'no', 'no')
Process.Model__CCF__01_Cell_Disk('10', '0.01', '.0001', '10', '10', '10', '0.1', 'Euler', 'Bi-conjugate Gradient Stabilized', '50', '1e-6', 'No', '.0001', 'Model/CCF/02 Mass Spring Solver', 'Model/CCF/10 Cell Tissue', 'Model/CCF/04 Mass Spring Growth', 'Model/CCF/05 Divide Cells', 'Model/CCF/06 Split Edges', 'Model/CCF/Auxin Gradient Model', 'Run_0_0-1_Linear_Quadratic_1-2_False_sim5_20231010', '140', '/home/ahr75/Documents/Shuyao/oofs_output/20231010', '10', 'True', 'False')
Process.Mesh__System__Open('Stack1', '/home/ahr75/Documents/Shuyao/oofs_2023-03-18/ext/02-CellModels/ext/04-MassSpring/ext/19-MassSpringAuxinQuadratic/Init_blank_72cells_2023-09-07.mdxm', 'no', 'no')
Process.Model__CCF__01_Cell_Disk('10', '0.01', '.0001', '10', '10', '10', '0.1', 'Euler', 'Bi-conjugate Gradient Stabilized', '50', '1e-6', 'No', '.0001', 'Model/CCF/02 Mass Spring Solver', 'Model/CCF/10 Cell Tissue', 'Model/CCF/04 Mass Spring Growth', 'Model/CCF/05 Divide Cells', 'Model/CCF/06 Split Edges', 'Model/CCF/Auxin Gradient Model', 'Run_0_1-0_Linear_Quadratic_0-4_False_sim5_20231010', '140', '/home/ahr75/Documents/Shuyao/oofs_output/20231010', '10', 'True', 'False')
Process.Mesh__System__Open('Stack1', '/home/ahr75/Documents/Shuyao/oofs_2023-03-18/ext/02-CellModels/ext/04-MassSpring/ext/19-MassSpringAuxinQuadratic/Init_blank_72cells_2023-09-07.mdxm', 'no', 'no')
Process.Model__CCF__01_Cell_Disk('10', '0.01', '.0001', '10', '10', '10', '0.1', 'Euler', 'Bi-conjugate Gradient Stabilized', '50', '1e-6', 'No', '.0001', 'Model/CCF/02 Mass Spring Solver', 'Model/CCF/10 Cell Tissue', 'Model/CCF/04 Mass Spring Growth', 'Model/CCF/05 Divide Cells', 'Model/CCF/06 Split Edges', 'Model/CCF/Auxin Gradient Model', 'Run_0_1-0_Linear_Quadratic_0-8_False_sim5_20231010', '140', '/home/ahr75/Documents/Shuyao/oofs_output/20231010', '10', 'True', 'False')
Process.Mesh__System__Open('Stack1', '/home/ahr75/Documents/Shuyao/oofs_2023-03-18/ext/02-CellModels/ext/04-MassSpring/ext/19-MassSpringAuxinQuadratic/Init_blank_72cells_2023-09-07.mdxm', 'no', 'no')
Process.Model__CCF__01_Cell_Disk('10', '0.01', '.0001', '10', '10', '10', '0.1', 'Euler', 'Bi-conjugate Gradient Stabilized', '50', '1e-6', 'No', '.0001', 'Model/CCF/02 Mass Spring Solver', 'Model/CCF/10 Cell Tissue', 'Model/CCF/04 Mass Spring Growth', 'Model/CCF/05 Divide Cells', 'Model/CCF/06 Split Edges', 'Model/CCF/Auxin Gradient Model', 'Run_0_1-0_Linear_Quadratic_1-2_False_sim5_20231010', '140', '/home/ahr75/Documents/Shuyao/oofs_output/20231010', '10', 'True', 'False')
```
