## Supplementary figures and images for "Tradeoff Between Speed and Robustness in Primordium Initiation Mediated by Auxin-CUC1 Interaction"

### auxConc_plast_0-8_t100.png

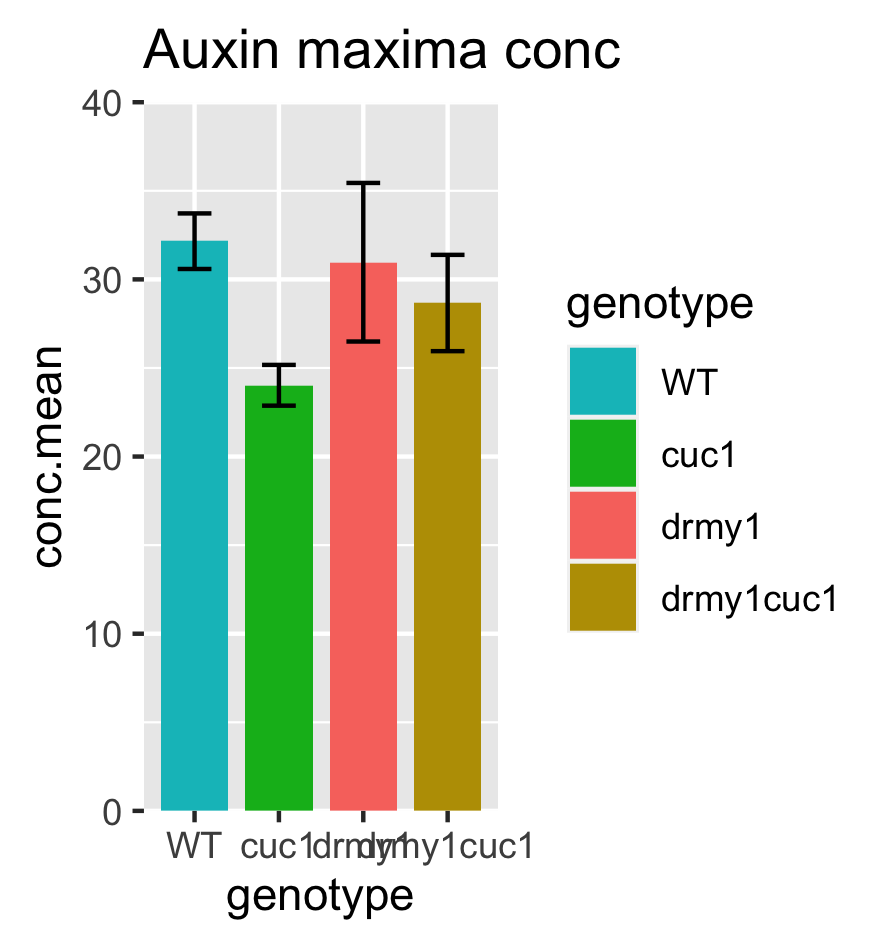

### AuxMaxNum_plast0-8.png

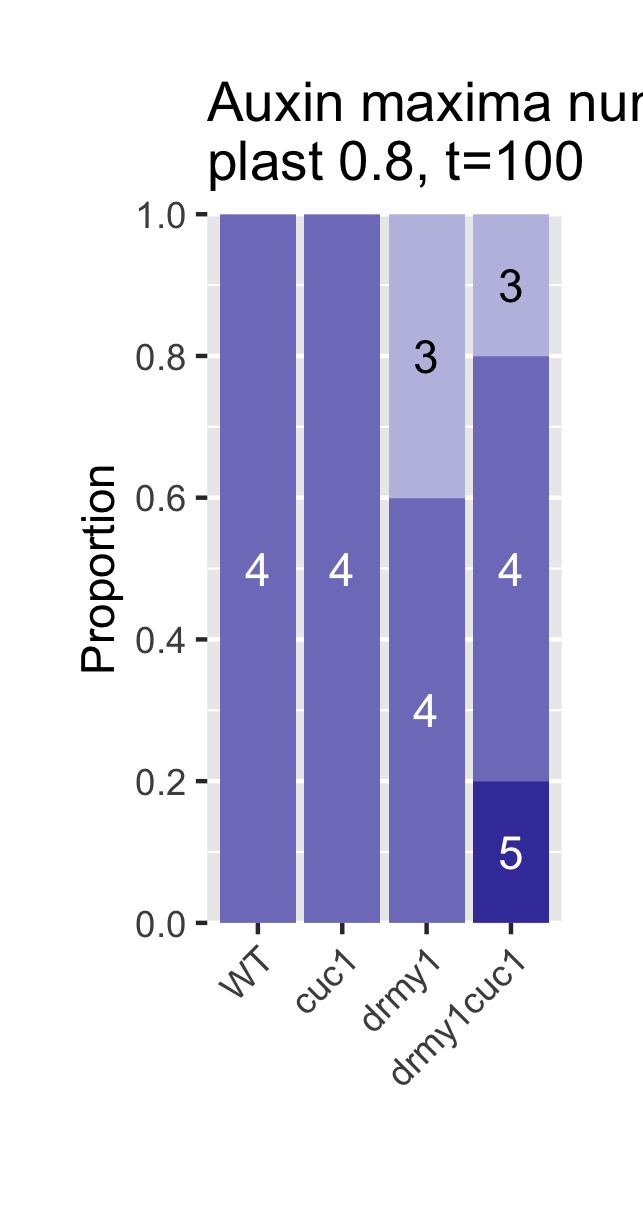

### AuxMaxNum_WT-drmy1_growth-rates.png

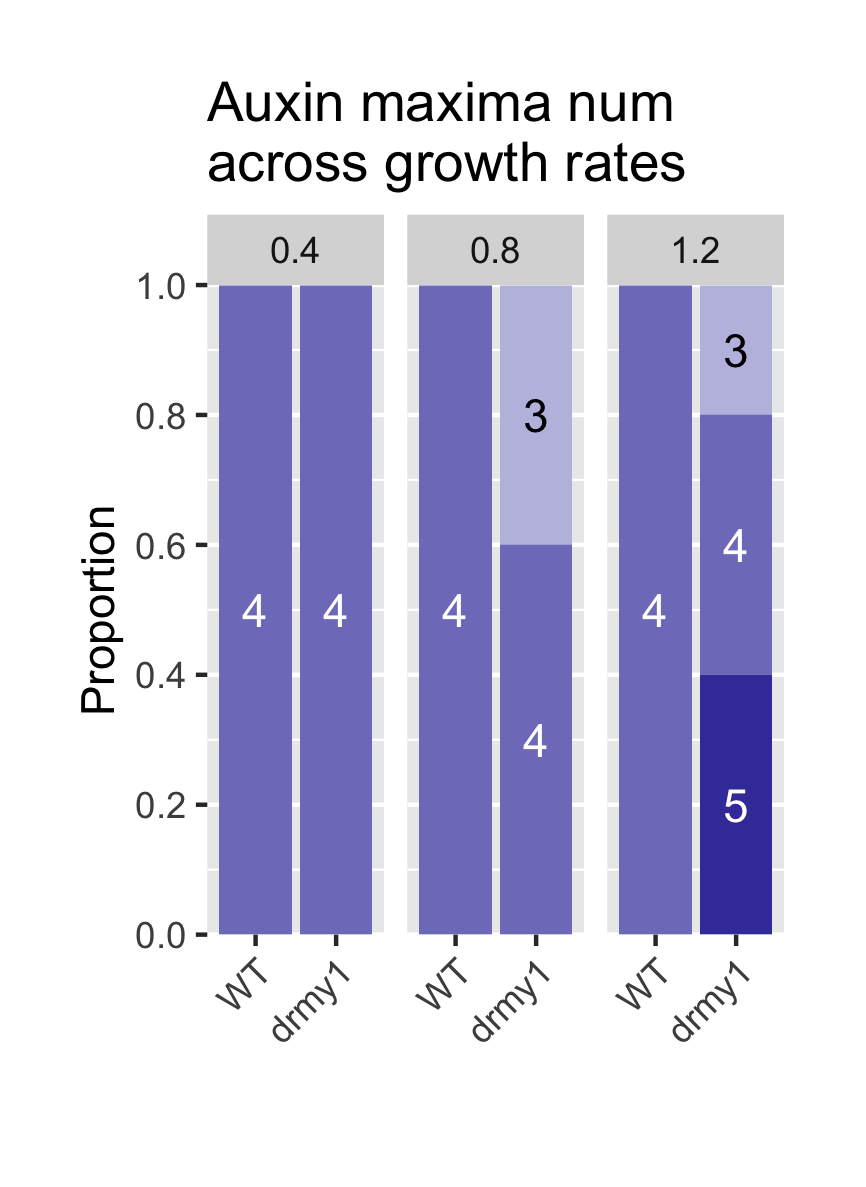

### CellDisk.png

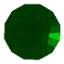
